# Evolutionary-Scale Enzymology Enables Biochemical Constant Prediction Across a Multi-Peaked Catalytic Landscape

**DOI:** 10.1101/2024.10.23.619915

**Authors:** Duncan F. Muir, Garrison P. R. Asper, Pascal Notin, Jacob A. Posner, Debora S. Marks, Michael J. Keiser, Margaux M. Pinney

## Abstract

Quantitatively mapping enzyme sequence-catalysis landscapes remains a critical challenge in understanding enzyme function, evolution, and design. Here, we expand an emerging microfluidic platform to measure catalytic constants—*k*_cat_ and *K*_M_—for hundreds of diverse naturally occurring sequences and mutants of the model enzyme Adenylate Kinase (ADK). This enables us to dissect the sequence-catalysis landscape’s topology, navigability, and mechanistic underpinnings, revealing distinct catalytic peaks organized by structural motifs. These results challenge long-standing hypotheses in enzyme adaptation, demonstrating that thermophilic enzymes are not slower than their mesophilic counterparts. Combining the rich representations of protein sequences provided by deep-learning models with our custom high-throughput kinetic data yields semi-supervised models that significantly outperform existing models at predicting catalytic parameters of naturally occurring ADK sequences. Our work demonstrates a promising strategy for dissecting sequence-catalysis landscapes across enzymatic evolution and building family-specific models capable of accurately predicting catalytic constants, opening new avenues for enzyme engineering and functional prediction.

## Introduction

Natural selection has shaped the catalytic parameters of enzymatic reactions, determining the rates and specificities that govern nearly all biological processes. The relationship between enzyme sequence and catalytic parameters is often conceptualized as a landscape, as first described by Fisher (*1*) and Wright (*2*), that is traversed through mutational “walks” (Fig. 1A). The topologies of sequence-catalysis landscapes, their responses to environmental changes, and their underlying molecular mechanisms are central to understanding how enzymes evolved to perform catalysis in distinct physicochemical environments (*3–7*). Furthermore, a quantitative mapping of these enzyme sequence-catalysis relationships can aid in developing predictive models for enzyme function and guide enzyme optimization.

**Figure 1.**
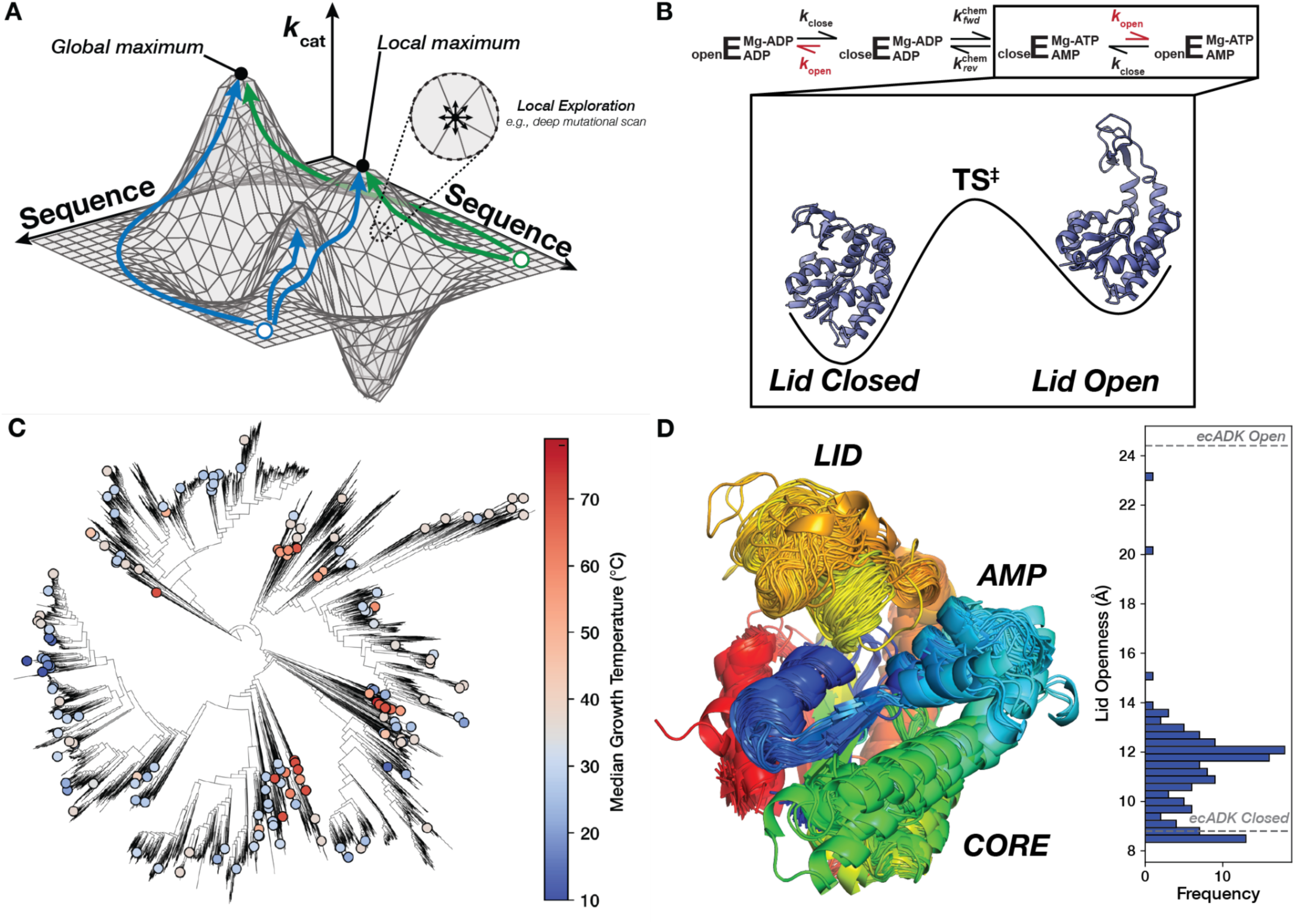
Mapping the sequence-catalysis of Adenylate Kinase across evolution. **(A)** Similar to a physical landscape, individual positions along the surface correspond to different enzyme sequences, with the height at each position representing their respective catalytic parameters. Sequences that occupy the highest “peaks” have the highest catalytic activities. The sequence space is effectively continuous for many sequences, allowing us to conceptualize the adaptive landscape as a two-dimensional surface in a three-dimensional space (*2*, *5*, *7*). While Deep-Mutational Scanning exercises assay many sequences, they explore a narrow region of sequence space (one mutation from wild-type) and are liable to be stuck in local optima. Green and blue paths show possible paths explorable by evolution that reach unique optima. **(B)** Schematic of the ADK reaction, where E represents the ADK enzyme, with the rate-limiting step, *k*_open_, shown in red (*46*). Experimental structures of closed (PDB: 4AKE) and open ecADK (PDB: 1AKE) are depicted on a simplified energy landscape for this reaction step. **(C)** The ADK sequence library characterized herein (dots) spans the bacterial tree of life, with organisms adapted to optimal growth temperatures ranging from the coldest to the hottest environments on Earth (*114*). **(D)** Superimposition of AlphaFold2 predictions of the ADK orthologs, with sequences trimmed to align with ecADK in an MSA. Measurements between the c-alpha of posistions 30 and 130 are histogrammed to display the range of observed LID “openness”, with measurements from experimental structures of ecADK in the open (PDB: 1AKE) and closed (PDB: 4AKE) conformations (*61*, *115*).

This inherent complexity of protein sequence space has often restricted sequence-function landscapes to simplified illustrations to conceptualize—but not quantify—evolutionary processes (e.g., Fig. 1A) (*7*). Current experimental explorations of enzyme sequence-catalysis landscapes are limited to narrow regions of the underlying protein sequence space. Combinatorial mutational studies have uncovered the role of intramolecular epistasis in shaping the evolutionary paths available to enzymes (*8–12*) but typically explore fewer than five sites (*9*, *10*, *13–20*). Alternatively, deep mutational scanning (DMS) studies measure a large number of sequence-fitness relationships (*21–23*). However, this approach explores a dense, local region of sequence space—all single mutations from wild-type—missing alternative peaks separated by multiple mutations. Furthermore, most mutational effects in DMS explorations are neutral or deleterious (*24–29*), limiting the potential to discover enzymes with improved catalytic parameters. In contrast, genomic sequencing has shown that orthologous enzymes exhibit significantly more sequence variation than is typically explored experimentally (*30*), potentially yielding substantial differences in catalytic properties from adaptation to diverse environments (e.g., temperature, [salt], pH). Nevertheless, catalytic activities are typically measured for few (typically <10) orthologs, leaving the topology of sequence-catalysis landscapes at the scale of naturally occurring sequences biochemically underexplored and underdetermined.

Existing approaches to model sequence-catalysis relationships are limited by a dearth of catalytic parameters for a given enzyme and narrow explorations of sequence space. In the absence of large-scale catalytic datasets, unsupervised deep-learning methods can learn complex distributions that may approximate these high-dimensional landscapes (*31*). Protein Language Models (PLMs), trained on the millions to billions of publicly available naturally occurring protein sequences, can often predict protein structure (*32*, *33*). Beyond structure, functional properties of enzymes, such as thermodynamic stability, folding kinetics, solubility, substrate specificity, and catalytic rate are presumably encoded in sequence, suggesting that sequence-only models may also learn sequence-function relationships. Furthermore, early PLM work demonstrated that the models learned an underlying representation of protein sequence that organized amino acids by biochemical property (*34*), and recent work has suggested their utility in optimizing protein function directly (*35*, *36*). Nevertheless, while PLM likelihoods (*37–40*), supervision on top of PLMs (*41*), and other unsupervised sequence-only models (*42*, *43*) can predict single mutation fitness effects, evaluating these representations of sequence-function landscapes remains challenging due to fundamental data challenges. Namely, these models are often trained or evaluated on DMS datasets, which typically use a fitness-linked proxy to estimate enzyme function. Various molecular properties can influence these fitness values, such as catalytic activity, stability, and inhibition, and these multiple low-level biochemical properties may buffer or amplify each other (*44*). This reliance on complex, indirect labels for protein function prediction may underlie the “yawning chasm” in our ability to accurately predict the relationships between sequence and specific enzyme properties (*6*). Consequently, there is a pressing need for quantitative datasets of gold-standard catalytic parameters collected under consistent conditions to test whether sequence-only models can accurately predict enzyme properties across diverse evolutionary and environmental contexts.

Here, we quantitatively map and dissect the topology of an evolutionary-scale sequence-catalysis landscape. We leverage an emerging microfluidics platform, High-Throughput Microfluidic Enzyme Kinetics (HT-MEK) (*45*), to measure the Michaelis-Menten parameters *k*_cat_ (catalytic constant), *K*_M_ (Michaelis-Menten constant), and *k*_cat_/*K*_M_ (catalytic efficiency) for hundreds of diverse naturally occurring and mutant sequences of the model enzyme adenylate kinase (ADK) (Fig. 1B, Fig. 1C). We dissect the topology and navigability of this sequence-catalysis landscape, showing that it is rugged, with at least three global peaks organized by structural motifs, yet this landscape remains navigable over long evolutionary timescales through path-dependent mechanisms. We address long-standing evolutionary hypotheses, for example, demonstrating that environmental temperature is not a primary determinant of enzyme catalytic rates and that thermophilic enzymes are not slower than their mesophilic counterparts. Finally, we show that a state-of-the-art unsupervised PLM organizes ADK sequence space by structure but not catalytic activity. Supervised and semi-supervised deep-learning models trained on the dataset newly collected herein to predict *k_cat_* outperform prior models trained on only public databases, with performance improving as the training dataset size grows. We provide a strategy to develop models that directly learn from family-specific sequence-catalysis landscapes and accurately predict fundamental biochemical constants of interest.

## Results and Discussion

### High-throughput measurement of ADK catalytic parameters across the bacterial and archaeal tree of life

The enzyme ADK is essential and ubiquitous across the tree of life. ADK regulates the cellular balance of adenylate currencies (ATP, ADP, and AMP), reversibly catalyzing the phosphoryl transfer between two ADP molecules (or ATP and AMP) using a Mg^2+^ cofactor that acts as an electrostatic “pivot” during the chemical step (Fig. 1B) (*46*). ADK consists of three domains: CORE, AMP-binding (AMP), and LID (Fig. 1D). The CORE domain forms most of the nucleotide-binding site and provides many catalytic residues needed for phosphoryl transfer. Prior NMR experiments with ADKs from diverse bacteria and archaea have revealed that the rate-limiting step of ADK catalysis is a large conformational change that opens two domains, LID and AMP, enabling product release (*47–49*). These conformational dynamics have made ADK a valuable model for studying the relationship between enzyme dynamics and catalytic turnover (Fig. 1B) (*46–48*, *50–53*).

To map the naturally occurring sequence-catalysis landscape of ADK, we gathered orthologous ADK sequences from bacteria and archaea from protein sequence databases (see Methods). We selected 193 orthologs based on sequence diversity and environmental adaptations. Specifically, we included ADK sequences from organisms adapted to divergent temperatures, as temperature adaptation has been suggested to drive differences in ADK catalytic parameters (*47*, *54–58*) (see Methods, Table S1). The resulting ADK sequence library has an average pairwise sequence identity of 42% and median optimal growth temperatures (*T*_Growth_) ranging from 9 to 96 °C (Fig. 1C, Fig. S1). Despite low average sequence identity, most substrate-contacting residues are >90% conserved, including the key catalytic arginine residues R36, R88, R123, R156, and R167 (*E. coli* ADK [ecADK] numbering used throughout) (*59*) (Fig. S2). Since most of our orthologs lack experimentally derived structures, we used AlphaFold2 (*60*) to predict their structures (Fig. 1D). Most ortholog structural differences were found in the predicted conformation of the LID domain, which varied in its openness. However, none of the predictions were as open as the fully open conformation of ecADK (*61*) (Fig. 1B, D).

Measuring Michaelis-Menten kinetics for >10^2^ enzyme sequences is intractable with traditional bench-top biochemistry methods. To overcome this challenge, we redeveloped an emerging microfluidic technology, HT-MEK (*45*), to assay all orthologous ADK enzyme kinetics in parallel under identical conditions. Briefly, we recombinantly expressed, purified, and assayed all 193 ADK orthologs on a single HT-MEK device (Fig. 2A). ADK enzymatic activity was monitored “on-chip” by coupling the formation of ATP to the production of NADPH (*62*), which was detected using time-lapse microscopy (Fig. 2B). All ADK orthologs were tagged at the C-terminus with a flexible Ser-Gly linker and eGFP, which enabled the purification of each ADK ortholog and subsequent quantification of per-chamber ADK concentrations (Fig. 2B). Observed initial rates collected over a range of ADP concentrations were normalized by ADK concentration and fit to the Michaelis-Menten equation, obtaining *k_cat_* and *K_M_* values for each ADK ortholog (Fig. 2C-D). Across the 1792 chambers in our device, we obtain an average of 6-7 biological replicates for each ADK ortholog per on-chip experiment. Out of the 193 ADK orthologs, 181 expressed and displayed catalytic activity above background, and of those, 175 had bounded *K*_M_ values under our assay conditions (Table S2, Methods). For ADK orthologs in our library that have been previously characterized in the literature under common conditions and assays, we observe a strong correlation between those *k*_cat_ measurements and our on-chip measurements (r^2^ = 0.96, Fig. 2E). Multiple controls provide confidence that these naturally occurring ADKs are natively-folded on-chip (Supplementary Text S1, Fig. S3-6).

**Figure 2.**
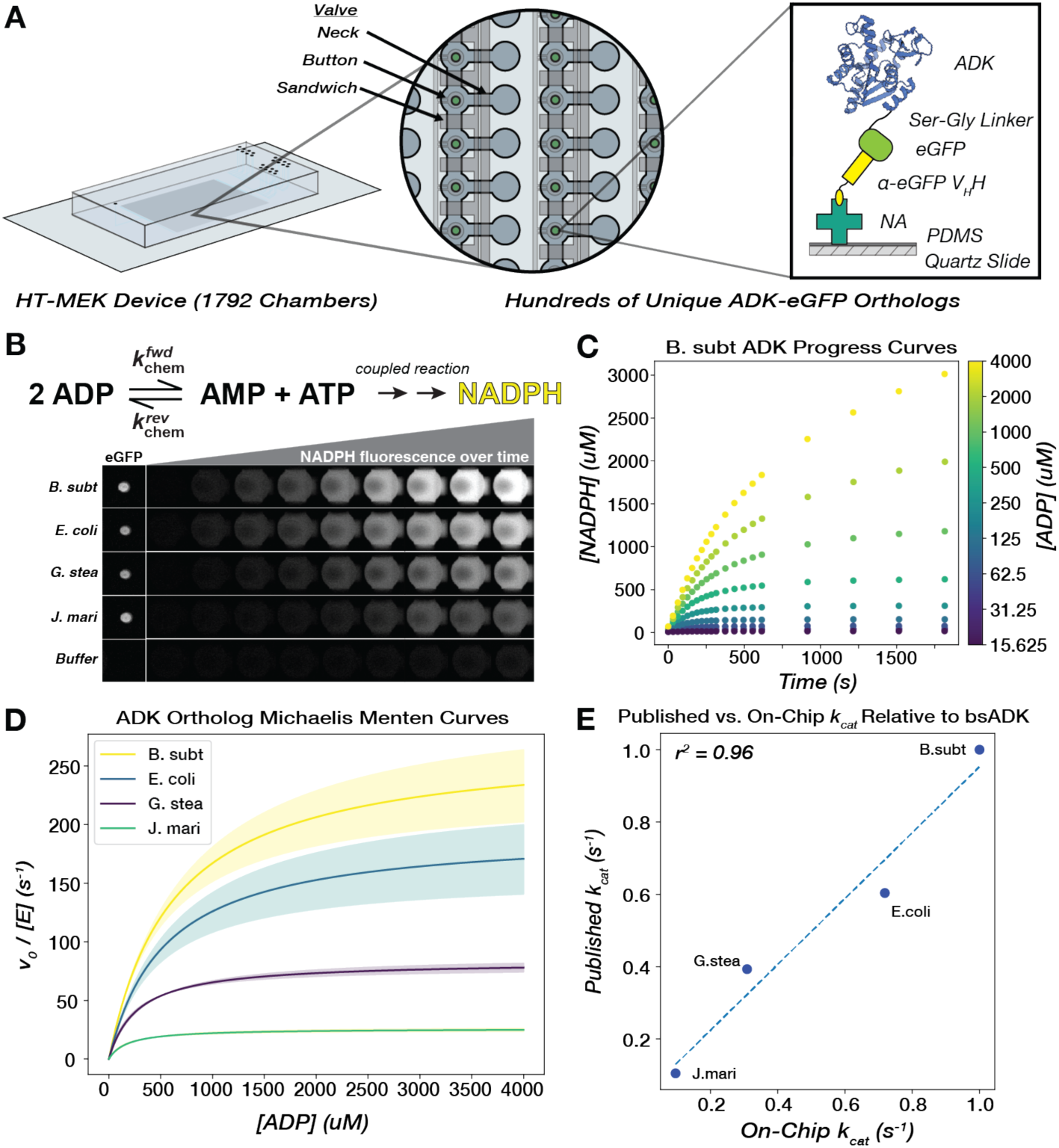
Michaelis-Menten parameters for hundreds of naturally occurring ADK sequences can be measured in parallel via high-throughput microfluidic devices. **(A)** A single High-Throughput Microfluidic Enzyme Kinetics (HT-MEK) device enabled the expression and purification of up to 1792 enzyme variants in a single experiment. All enzyme variants are tagged with a C-terminal eGFP construct to facilitate capture on a functionalized “pedestal.” This pedestal consists of neutravidin proteins (NA) non-specifically bound to a PDMS-coated quartz slide, which in turn binds a biotinylated anti-eGFP VHH nanobody, pulling down the eGFP-tagged ADK in each chamber. **(B)** ADK activity in the direction of ATP formation is monitored on-chip through coupled production of NADPH, and product formation is measured over the course of the assay with time-lapse inverted fluorescent microscopy. Four chambers containing exemplary orthologs are highlighted, as well as a control chamber that did not contain an ADK-encoding plasmid and thus did not show any expressed enzyme or detectable catalysis. **(C)** Scatter plot of progress curves for bsADK across multiple substrate concentrations (encoded by color). **(D)** Mean fits of initial rates to the Michaelis-Menten equation for four ADK orthologs. Shaded regions represent the standard deviation of *k_cat_* across biological replicates for each ortholog. **(E)** On-chip catalytic measurements correlate with previously published values for orthologs, relative to *bs*ADK, measured in the same reaction direction under comparable conditions (Methods).

Remarkably, while the *k*_cat_ values of all natural enzymes vary by ∼5 orders of magnitude (*63*), the *k*_cat_ values within the ADK family alone span at least three orders of magnitude, ranging from 1–803 s^-1^ (Fig. 3A, B, Fig. S7). Thus, while the slowest measurable ADK in our library had a *k*_cat_ of 1 s^-1^, certain sequence combinations found in nature achieve catalytic turnover at least two orders of magnitude higher, approaching the activities of some of the fastest known enzymes (*63*). Thus, naturally occurring sequences performing analogous functions across different organisms can exhibit catalytic activities spanning orders of magnitude despite having superimposable structures, experimental and predicted, and nearly identical active sites (Fig. 1D, Fig. S2). This finding underscores the challenge of predicting catalytic function from sequence alone.

**Figure 3.**
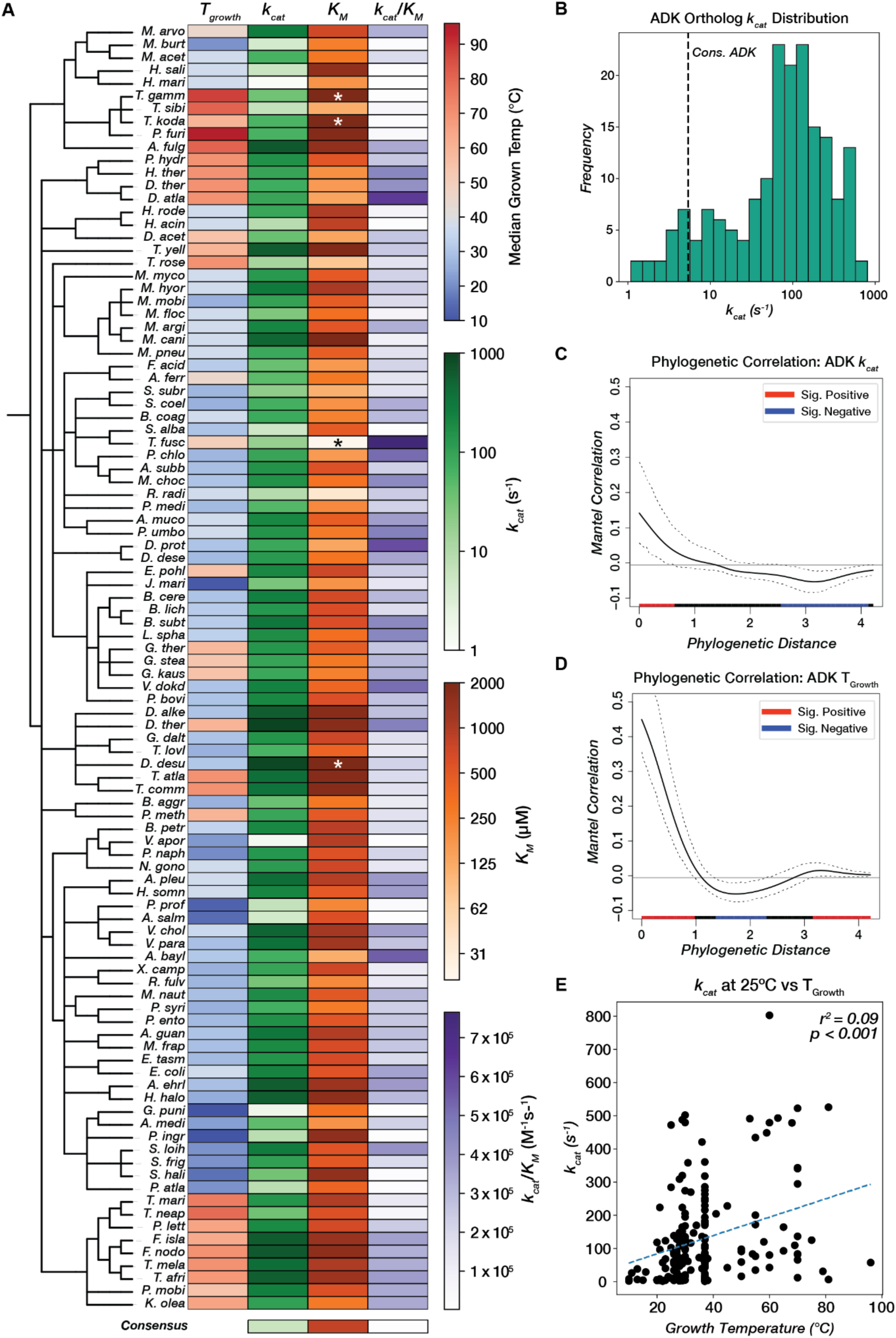
Adenylate kinase catalytic parameters span three orders of magnitude and correlate weakly with phylogeny and environmental conditions. **(A)** Catalytic parameters and *T_Growth_* values for 100/181 ADK orthologs (including those with *K_M_* outside the range of the assay) are displayed as a heatmap across a taxonomic tree (Methods). *k_cat_* and *K_M_* are colored on a log scale. Orthologs with *K_M_* outside the range of the assay are labeled with asterisks (*). See Fig. S7 for a heatmap of all 181 ADK orthologs. The consensus ADK sequence (*94*) is plotted at the bottom (*k_cat_* = 5.4 *s^-1^*, *K_M_* = 804 uM). **(B)** Measured *k_cat_* values for 175 orthologs span three orders of magnitude. The vertical dashed line represents the *k*_cat_ for the ADK consensus sequence (*94*). **(C,D)** Phylogenetic signal analysis of (**C**) *k_cat_* and (**D**) *T_Growth_*. Moran’s I index of autocorrelation is plotted as a solid black line, with the 95% confidence interval outlined by dashed black lines. The colored bar at the bottom encodes the significance of autocorrelation, with red and blue representing positive and negative significant autocorrelation, respectively, and black representing nonsignificant autocorrelation. **(E)** Linear regression analysis between optimal growth temperature and *k_cat_*. The regression line is plotted as a dashed blue line (r^2^ = 0.09, p < 0.001).

Given the wide variation in catalytic rates, we next explored whether ADK variants close in evolutionary space exhibit similar activities. Mapping the catalytic parameters *k*_cat_, *K*_M_, and *k*_cat_/*K*_M_ to the organismal taxonomic tree shows very little visual organization (Fig. 3A). To quantify this relationship between phylogenetic distance and catalytic rate, we computed the phylogenetic signal, which quantifies the autocorrelation between distance on a phylogenetic tree and continuous traits (Methods) (Fig. 3C-D, Fig. S8) (*64*). We quantified this relationship for *k*_cat_, *K*_M_, *k*_cat_/*K*_M_, and *T_Growth_* and observed that *k*_cat_ values show a significant positive correlation over short phylogenetic distance (Fig. 3D), albeit weaker than *T_Growth_* phylogenetic signal (Fig 3D). However, across medium to long phylogenetic distances, *k*_cat_ is decorrelated with phylogeny (Fig 3C), with high *k*_cat_ values often interspersed among comparatively low ones. Similar behaviors are observed for *K*_M_ and *k*_cat_/*K*_M_ (Fig. S8). This result suggests that high catalytic activity has independently evolved multiple times during ADK evolution along distinct lineages. Thus, computational models for predicting catalytic activity must encompass multiple evolutionary—and potentially mechanistic—routes to high activity.

### Many thermophilic orthologs remain highly active at mesophilic temperatures, and psychrophilic ADKs are not catalytically superior

Next, we explored whether the distinct environmental niches and selection pressures of each organism were responsible for the variation and evolutionary distribution we observed in ADK catalytic rates. We focused on temperature, a pervasive environmental factor reported to drive adaptive changes in enzyme catalytic rates (*54*, *62*, *65–73*). Discussions of activity-stability trade-offs propose that stabilizing mutations in thermophilic enzymes increase rigidity during natural evolution, suppressing activity-promoting dynamics (*65–68*, *71*, *74–78*). Accordingly, psychrophilic enzymes are frequently reported to have higher *k*_cat_ values relative to their mesophilic and thermophilic counterparts (*65–68*, *76*). However, these findings typically rely on comparisons of just a pair of sequences, and recent analyses of enzyme kinetic data in BRENDA (*79*) question the generality of this model (*80*).

We systematically tested this model across a wide range of growth temperatures under consistent conditions. When *k*_cat_ is plotted against the optimal growth temperature of the corresponding organisms, we do not observe the expected negative correlation but rather a weak positive correlation (Fig. 3E). Similarly weak trends are observed for *K*_M_ versus growth temperature, providing evidence against related hypotheses that enzymatic *K*_M_ values are increased in cold-adapted enzymes (Fig. S9) (*65*, *68*, *71*, *81*). Since growth temperature correlates with melting temperature and thermodynamic stability, our data show that ADK activity and stability *do not* universally trade off during natural evolution. Consequently, psychrophilic ADKs are not universally catalytically superior, and thermophilic ADKs are not catalytically limited. In fact, some of the fastest ADKs come from thermophilic organisms, potentially because high ADK activity is needed at increased temperatures to regenerate the ATP pool to combat its thermal lability (*82*). These findings are particularly intriguing for ADK, which has been central to the long-standing debate on the trade-off between thermodynamic stability and catalytic activity, and show that natural evolution can jointly optimize stability and activity (*47*, *54*, *62*, *66*, *83–85*). The joint optimization of both properties is perhaps facilitated by the independent folding of the CORE and LID domains, allowing the CORE to evolve high thermodynamic stability while the LID retains the necessary flexibility for domain opening along the reaction path (*86*). Our findings underscore the importance of examining sequence-catalysis landscapes on a broad scale. While small, localized studies may suggest the existence of an activity-stability trade-off (*65*, *66*, *68*, *71*, *74–78*), the larger sequence space reveals that this trend does not hold universally for natural ADK sequences. Furthermore, our results strongly indicate that enzymatic catalytic rates are not under strong selection during temperature adaptation. Instead, recent bioinformatic trends observed in enzyme sequences across a wide range of *T*_Growth_ values are likely driven primarily by changes in stability rather than activity (*87*).

### The evolutionary-scale ADK sequence-catalysis landscape is rugged, with multiple structural peaks achieving distinct activity levels

While protein “sequence space” is often discussed in conceptual terms, its high dimensionality (20 x sequence length) makes its concrete visualization challenging. To further dissect the topology of this landscape, we visualized the sequence space as a graph, where each node corresponds to an orthologous ADK sequence, and edges connect these nodes, weighted by the number of edits in a multiple sequence alignment (MSA). Using our 175 observed sequences, we traversed this subset of ADK sequence space by minimizing the total edit distance, forming a Minimum Spanning Tree (MST). This MST visually represents the rugged sequence-catalysis landscape of naturally occurring ADKs (Fig. 4A).

**Figure 4.**
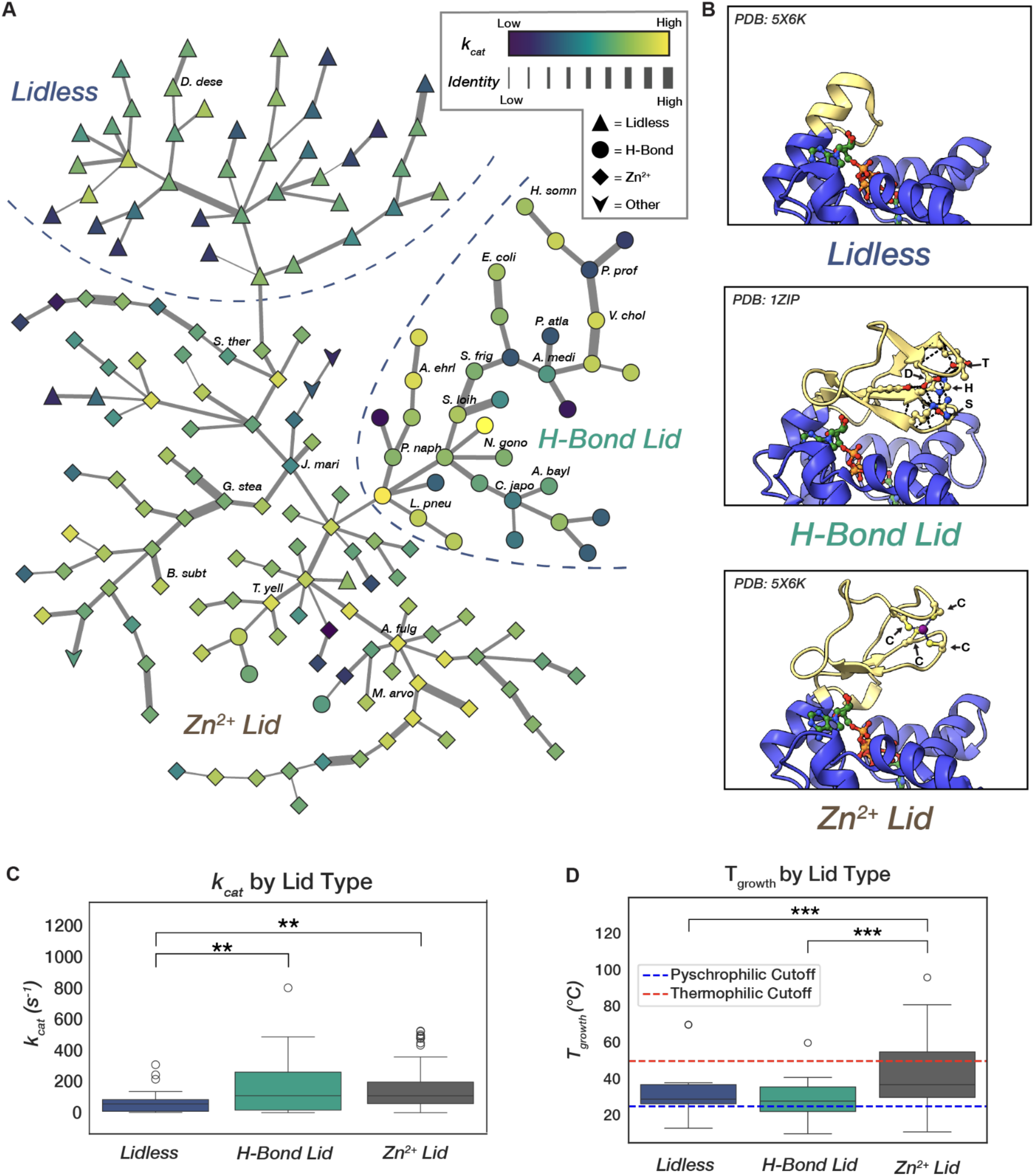
Adenylate Kinase sequence space features multiple “peaks” linked to different lid types, each associated with distinct growth temperatures and reaching varying “heights” in activity. **(A)** A Minimum-Spanning Tree (MST) from the all-by-all graph of ADK sequences with edges weighted by edit distance in an MSA. Edge thickness in the MST encodes increasing edit distance on a log scale. Node color encodes measured *k_cat_* value in log scale, and node shape corresponds to lid type. Rough partitions between lid-type neighborhoods are labeled and outlined with dashed lines. Selected orthologs discussed in depth in this study are labeled. **(B)** The three major types of LID domains found in ADKs: “Lidless” containing a short loop that still provides catalytic Arg residues, and two lidded variations, one containing hydrogen-bonding network consisting of a conserved His-Ser-Asp-Thr tetrad (H-Bond Lid) and the other containing a cysteine tetrad chelating a Zn^2+^ ion. **(C)** H-Bond and Zn^2+^ Binding Lid ADKs are significantly faster than lidless counterparts but have similar activity distributions to one another (ANOVA, F=6.92, p-val=0.002; Tukey-HSD, Zn^2+^ vs. Lidless p-adj=0.002, H-Bond vs. Lidless p-adj=0.006, Zn^2+^ vs. H-Bond p-adj=0.963). **(D)** Zn^2+^ Binding Lid ADKs have a significantly higher associated growth temperature than the other lid-types (ANOVA, F=16.39, p-val=3.11e-7; Tukey-HSD, Zn^2+^ vs. Lidless p-adj=0.0005, H-Bond vs. Lidless p-adj=0.370, Zn^2+^ vs. H-Bond p-adj<1.0e-16). Cutoffs for psychrophilicity and thermophilicity are shown as dashed lines (*103*).

Since the ADK MST shows little organization by catalytic activity, we next explored whether ADK structural differences might organize it. The LID of ADK exists in three general forms: (1) “Lidless” ADKs, characterized by unstructured loops of varying lengths, present across all domains of life (Fig. 4B, top); (2) “Zn²⁺-binding” lids, which feature a structural Zn²⁺-binding site typically formed by four Cys residues, found in bacteria and archaea (Fig. 4B, middle); and (3) “H-Bond” lids, which possess an extended hydrogen bond network and are found almost exclusively in bacteria (Fig. 4B, bottom) (*88–90*). Visualizing the ADK MST reveals that these three lid types—Zn²⁺-binding, H-Bond, and Lidless—each cluster into distinct neighborhoods, or “peaks” (Fig. 4A). A closer examination of these structural peaks reveals that while the two lidded peaks span similar dynamic ranges of catalytic activity, encompassing both the fastest and slowest ADKs, the Lidless sequences are significantly slower (Fig. 4C). Given that Lidless ADKs span all domains of life, while Zn²⁺-binding and H-Bond lids are limited to bacteria and archaea or bacteria alone, respectively (*90*), lid domains may have been acquired later in ADK evolution, potentially as a mechanism to enhance catalytic rates.

ADKs adapted to different environmental temperatures inhabit different structural peaks within the landscape. ADKs from psychrophilic organisms (*T*_Growth_ < 25 °C) have mostly H-Bond lid types, whereas ADKs from thermophilic organisms (*T*_Growth_ > 50 °C) have nearly exclusively Zn²⁺-binding lids (Fig. 4D, Fig. S10). The Zn²⁺-binding lid may be favored in thermophilic ADKs because it provides extra stability to the domain at high temperatures, consistent with prior studies that show an increase in melting temperature (T_m_) by 15 °C when installing a Zn²⁺ binding site in the H-Bond lid of *E. coli* ADK (*91*). In contrast, Zn²⁺ can be limiting in marine environments where many psychrophilic organisms live, potentially favoring the H-Bond over the Zn²⁺-binding lid (*92*). Intriguingly, thermophilic ADKs are predominantly located at internal nodes within the MST, indicating they share more sequence similarity with other ADKs compared to their mesophilic and psychrophilic counterparts (Fig. S10). This result aligns with previous observations of highly thermostable consensus sequences in enzyme families (*93–98*). Thus, while the temperature of an organism’s environment does not appear to directly drive changes in catalytic properties (Fig. 3E), it does influence the distribution of organisms across this evolutionary-scale sequence-catalysis landscape, confining psychrophilic and thermophilic ADKs to specific neighborhoods of distinct lid structures (Fig. 4D, Fig. S10).

### Navigating between peaks: mutational walks and extra-dimensional bypasses

Given that all three lid types appear in nature, these peaks must have been accessible to one another throughout evolution. To explore the viability of transitioning between these lid-type peaks in our landscape, we first focused on the Zn²⁺-binding and H-Bond peaks. We generated combinatorial mutations to swap the four key residues that constitute the Zn²-binding motif–Cys130, Cys133, Cys150, and Cys153 (CCCC) in *G. stearothermophilus* (gsADK)–with the corresponding residues in the H-Bond motif–His126, Ser129, Asp146, and Thr149 (HSDT) in ecADK (Fig. 4B, Fig. S11). We measured *k*_cat_ and *K*_M_ for all possible mutations that interconvert the Zn²⁺-binding and H-Bond motifs in gsADK and ecADK backgrounds in a single high-throughput experiment. When installing a Zn²⁺-binding motif in the *E. coli* ADK LID, the final Zn²⁺-binding motif (CCCC) retains 74% of the activity of the wild-type H-Bond motif (HSDT), consistent with previous studies (*91*), and there are mutational pathways without highly deleterious intermediates (Fig. 5A). Given the feasibility of a mutational walk from the H-Bond to the Zn²⁺-binding lid, we anticipated that the reciprocal walk in gsADK—starting with the Zn²⁺-binding motif (CCCC) and ending with the H-Bond motif (HSDT)—should also be possible. Surprisingly, nearly all mutational trajectories in this direction encounter an unfavorable intermediate or “pit,” leading to a large loss of activity or an inexpressible variant (Fig. 5B, Table S3). This non-reciprocal behavior aligns with differences in the three-dimensional context of the two structural motifs. While the CCCC motif coordinates the Zn²⁺ ion independently of neighboring residues, the H-Bond motif is more complex, involving additional interactions with second-shell residues, presumably contributing to the observed epistasis of the HSDT motif (Fig. S12). Supporting this model, the H-Bond lid occupies a narrower region of sequence space compared to the Zn²⁺-binding lid (79% vs. 65% average sequence identity, respectively, Fig. S13). This indicates that the sequence contexts available for the lid beyond the HSDT/CCCC motifs are more constrained in H-Bond lids than in Zn²⁺-binding lids.

**Figure 5.**
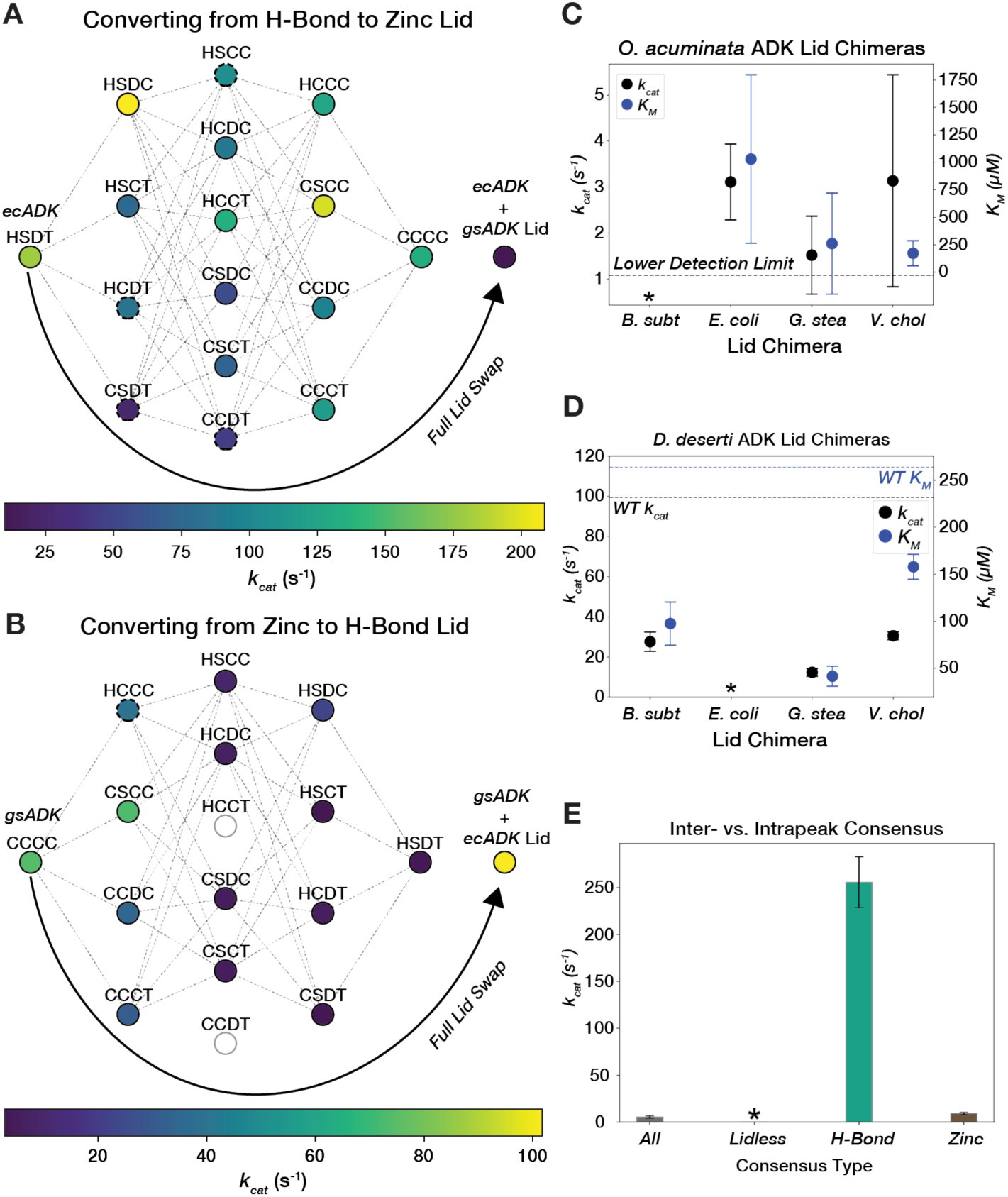
Traversing the ADK sequence-catalysis landscape through mutational walks and “extra-dimensional bypasses”. **(A)** A graph showing mutational pathways of swapping the chelating cysteine tetrad into the H-Bond lid of ecADK. Nodes represent variants along the pathway, with their *k_cat_* value encoded by color. Dashed lines connect variants that are one mutation away from each other. Dashed circles represent variants with *K_M_* fit outside of the bounds of the assay. Fully swapping the ecADK H-Bond LID for gsADK’s Zn^2+^-binding LID is shown as an arrow. **(B)** A graph showing mutational pathways of swapping the hydrogen-bond motif into the Zn^2+^ lid of gsADK. Nodes represent variants along the pathway, with their *k_cat_* value encoded by color. Dashed lines connect variants that are one mutation away from each other. Dashed circles represent variants with *K_M_* fit outside of the bounds of the assay. Fully swapping the gsADK Zn^2+^-binding LID for ecADK’s H-Bond LID is shown as an arrow. Empty circles represent variants that expressed poorly or displayed activity below the limit of detections and are not connected by edges in the graph. **(C)** *k_cat_* and *K_M_* are plotted for LID chimeras of ocADK. Error bars represent standard deviation across biological replicates. The lower detection limit for *k_cat_* is plotted as a black dashed line. The bsADK LID chimera displayed activity below the lower limit of detection and is marked with an asterisk. **(D)** *k_cat_* and *K_M_* are plotted for LID chimeras of ddADK. Error bars represent standard deviation across biological replicates. The lower detection limit for *k_cat_* is plotted as a black dashed line. The ecADK LID chimera exhibited a *K_M_* below the lower bound of the assay and is marked with an asterisk. **(E)** Barplot of *k_cat_* for consensus sequences, with error bars representing standard deviation across biological replicates. The lidless consensus sequence did not exhibit activity above background.

We next explored whether navigation between these peaks could be achieved through “extra-dimensional bypasses” (i.e., whole LID swaps), thereby avoiding unfavorable valleys (*5*). Substituting the ecADK H-Bond LID into gsADK results in a highly active ADK that is even faster than wild-type gsADK (Fig. 5B). Interestingly, the reverse experiment, where the entire LID of ecADK is swapped into gsADK, leads to a dramatic decrease in activity, dropping below the level of any of the incremental cysteine mutants. These results suggest that to avoid inactive intermediates, whole LID swaps are necessary to convert a Zn²⁺-binding lid into an H-Bond lid, whereas, in the reverse direction, it is more favorable to convert an H-Bond lid to a Zn²⁺-binding lid through single mutational steps.

Considering the existence of Lidless ADKs, which may represent the ancestral form of all ADKs given their presence across all domains of life (*90*), we explored navigating our landscapes through the insertion or deletion of the LID. Since we found Lidless ADKs to be significantly slower than their lidded counterparts (Fig. 4C), we asked whether their activity could be “rescued” by inserting a LID. Given the variability in the degree of “lidlessness”, we selected two Lidless ADKs for our study: *O. acuminata* ADK (ocADK), which has a seven amino-acid loop and expressed well but showed activity below our detection limit in the initial assay, and *D. deserti* ADK (ddADK), which has a longer 17 amino-acid loop and displayed above-average activity among Lidless ADKs. We generated chimeras incorporating the LIDs from gsADK, bsADK, ecADK, and vcADK into both ocADK and ddADK backgrounds. For ocADK, we found that three out of four LID chimeras successfully rescued catalytic activity above our detection limit (Fig. 5C). In contrast, for ddADK, which already had modest activity, we instead observed a decrease in *k*_cat_ with a commensurate decrease in *K*_M_ for measurable lid insertions (Fig. 5E). Therefore, LID insertion does not always increase *k*_cat_, complicating the prediction of ADK activity, which depends on LID dynamics but not solely on the LID sequence due to potential functional coupling between these domains (*55*).

### Contributions from multiple peaks catalytically impair the ADK consensus sequence

Consensus sequences have garnered interest for engineering proteins with enhanced stability while preserving catalytic activity, though the retention of activity depends on the specific natural enzyme used as a benchmark (*93–98*). For ADK, we anticipated that a consensus sequence incorporating elements from all three peaks would produce suboptimal amino acid combinations. Indeed, a previously constructed ADK consensus sequence that spans the landscape has a *k*_cat_ value of 5 s^-1^, making it slower than 91% of the naturally occurring sequences we measured (Fig. 3A) (*94*). To further test this hypothesis, we independently constructed consensus sequences from each of the three peaks. We observed that the internal consensus sequences for ADKs with H-Bond and Zn²⁺-binding LIDs were 47- and 1.6-fold faster in *k*_cat_ compared to the consensus sequence for the entire family (Fig. 5E). The larger increase in *k*_cat_ for the H-Bond lid consensus sequence aligns with the narrower sequence space for this lid type (62% vs. 45% internal pairwise sequence identity for H-Bond and Zn²⁺-binding LID orthologs, respectively, Fig. S14). The internal consensus sequence for the Lidless ADKs exhibited activity below our detection limit, perhaps due to the variability in loop length among Lidless ADKs (Fig. 5E). These results emphasize the importance of functional coupling between residue positions and the role of intramolecular epistasis in shaping sequence-catalysis landscapes (*8–12*).

### Changes in ADK dynamics tune activity across billions of years of evolution

Since high ADK activity has evolved multiple times in distinct structural contexts, we investigated the molecular mechanisms underlying high activity in each peak to determine if they are general or unique. Prior work in ecADK has shown that interconversion between the closed and open states involves the local unfolding of the LID and that this conformational equilibrium can be tuned with osmolytes and mutations (Fig. 1B) (*48*, *50*, *90*, *99*, *100*). In particular, urea, which can stabilize the unfolded state of proteins, favors the more expanded, open forms of ecADK, increasing *k*_cat_ by 1.7-fold at 2 M urea (*99*). We hypothesized that if changes in conformational dynamics were driving the differences in ADK catalysis, we would observe this capacity for conformational tuning across most naturally occurring ADK sequences. A closer examination of ADK activity under gradually increasing concentrations of urea (0.0–2.0 M) supports this hypothesis: most naturally occurring ADKs exhibited activation by urea, including for all three lid types, suggesting that the capacity for conformational tuning has been conserved in multiple structural contexts (Fig. 6A, 6B, Fig. S6). Previous reports of activation of thermophilic enzymes by denaturants were interpreted as increasing motions of these presumably overly rigid enzymes (*101–103*). Here, we systematically demonstrate that low concentrations of urea activate most naturally occurring ADKs, regardless of their parent organisms’ growth temperature, challenging the model of rigidity-activity trade-offs during temperature adaptation (*65*, *66*, *68*, *71*, *74–78*).

**Figure 6.**
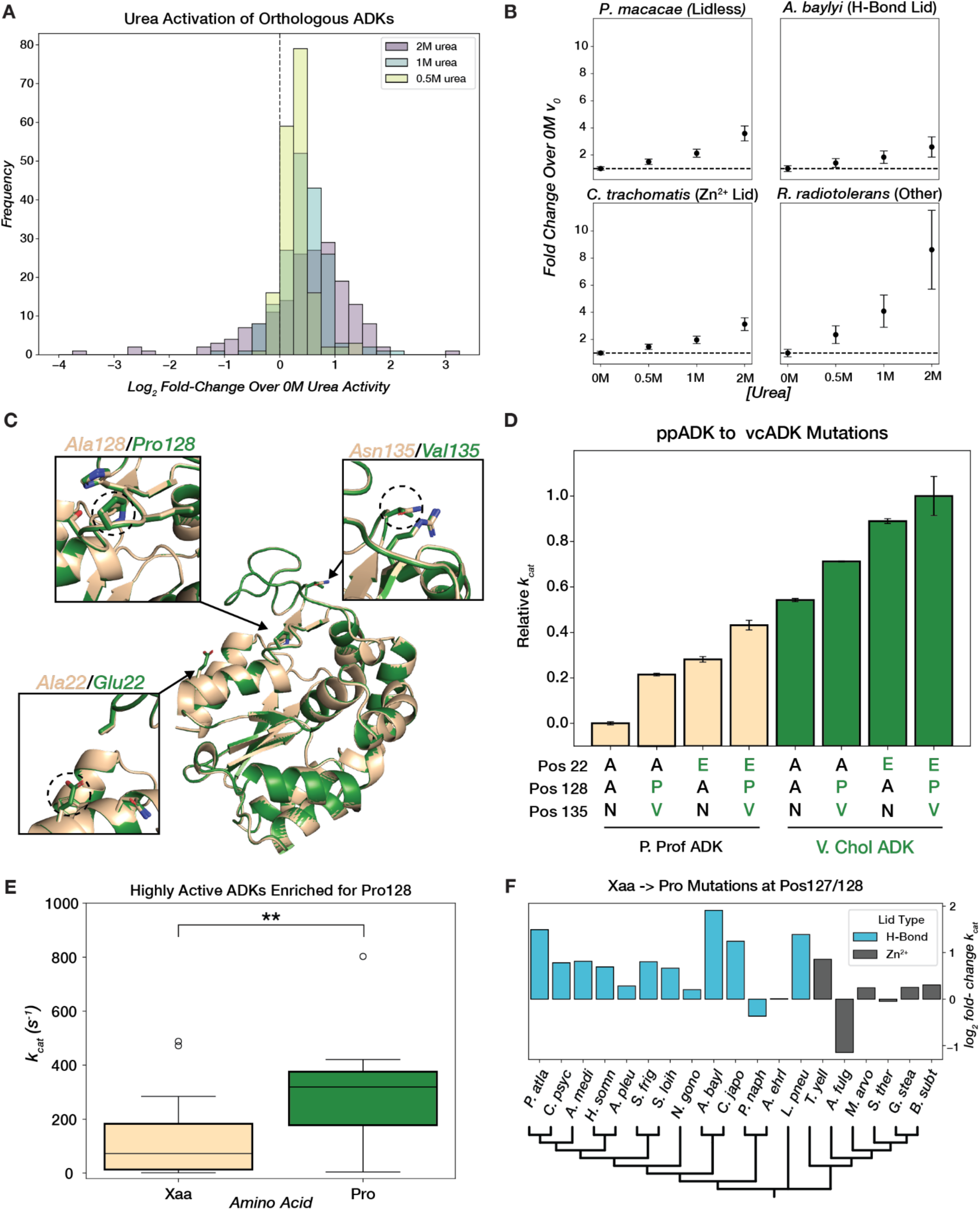
ADK conformational tuning with osmolytes and mutations across evolution. **(A)** Distribution of log_2_ fold-change in initial reaction rate at saturating substrate concentration relative to 0M urea for 0.5M, 1M, and 2M urea**. (B)** Mean fold-change in initial rate over 0M urea at 4mM [substrate]. Error bars represent standard deviation across replicates. The black dashed line represents no change in initial rate. **(C)** Superimposed AF2 models of ppADK (tan) and vcADK (green) with positions 22, 128, and 135 highlighted. **(D)** Barplot of mean relative catalytic effects (technical replicates, n=2) of mutations at key positions that differ between ppADK and vcADK. Error bars represent standard deviation across replicates. Variants in ppADK background are plotted in tan and vcADK background in green. Amino acid identity at positions 22, 128, and 135 are displayed below each bar. *k*_cat_ values for ppADK and vcADK mutants were collected off-chip (see Methods). **(E)** Boxplot of *k_cat_* for H-Bond LID ADKs that have a proline (green) or different amino acid (tan) at position 128. t(37)=-3.162, p=0.003. **(F)** Barplot of log_2_ fold-change in *k_cat_* for Xaa→Pro mutations in selected ADKs with either an H-Bond or Zn^2+^ LID, organized by a phylogenetic tree (plotted with arbitrary branch lengths).

To identify mutations that influence ADK conformational dynamics throughout evolution, we turned to orthologs with high sequence similarity but large differences in catalytic rates. We focused on ADKs from *V. cholerae* (vcADK) and *P. profundum* (ppADK), as they exhibited the largest ratio of fold-change in *k*_cat_ to edit-distance–a >50-fold difference in activity over 23 sequence differences–indicating the steepest activity “cliff” in our landscape (Fig. 4A, Fig. 6C). We hypothesized that functionally-relevant sequence positions would be in the LID. There are two key residue differences in the LID of ppADK and vcADK: Ala/Pro at position 128 and Asn/Val at position 135 (Fig. 6C). Indeed, the double mutation Ala128Pro/Asn135Val increases ppADK *k*_cat_ by 21%, while the reverse mutation, Pro128Ala/Val135Asn, decreases vcADK *k*_cat_ by 11% (Fig. 6D, Methods). An additional CORE mutation, Ala22Glu, was also selected because its proximity to the LID led us to hypothesize that the increased bulkiness and charge of the Glu side chain could destabilize the enzyme’s closed state by electrostatic repulsion (Fig. 6C). The Ala22Glu mutation increased ppADK *k*_cat_ by 28%, while the reverse Glu22Ala mutation decreased activity in vcADK by a reciprocal 29% (Fig. 6D). Together, the Ala22Glu/Pro128Ala/Val135Asn mutations account for 49% of the activity difference between ppADK and vcADK (Fig. 6D).

Next, we assessed the generality of these activation mechanisms across the ADK landscape. Pro128 is frequently found in ADKs and shows a statistically significant association with high *k*_cat_ across diverse ADK sequences (Fig. 6E). We selected 20 H-Bond and Zn²⁺-binding ADKs and either mutated the residue at position 128 to proline or, in cases where position 128 was already proline, we mutated P128 to alanine. In the Xaa-to-Pro mutations, we primarily observed activating effects in ADKs with H-Bond lids (Fig. 6F). In Zn²⁺-binding ADKs, the activating effect of Pro was less pronounced, although most still showed increased activity (Fig. 6F). This smaller effect may be due to Pro128’s position between Cys130 and Cys133 (*B. subtilis* ADK numbering), which coordinate the structural Zn²⁺ (Fig. S12). In contrast, mutational effects at position 22 do not generalize to other orthologs (Fig. S15). Thus, conformational tuning can regulate ADKs separated by billions of years of evolution, whether through mutations or small molecule solutes.

### PLMs organize ADK sequence space by structure but not by catalytic activity

We next explored the landscape of our naturally occurring ADK sequences as learned by the state-of-the-art model ESM-2 (*32*) to evaluate the sequence-catalysis relationship of PLMs. We fed representative ADK sequences from sequence databases (∼5,000 sequences, see Methods) into the pre-trained 650-million-parameter ESM-2 model to obtain fixed-length embeddings for each ADK ortholog after mean-pooling (Methods). The continuous nature of these embeddings allows us to visualize the traditional concept of landscape using dimensionality reduction techniques like UMAP (*104*). We observed that the ADKs measured in our library broadly cover the landscape generated from ADK ESM-2 embeddings, with the various lid types forming distinct visual clusters (Fig. 7A). This structural organization aligns with our MST-derived sequence-catalysis landscape, with distinct lid-type peaks (Fig. 4A). Consistent with our MSA, H-Bond ADKs had the highest internal sequence identity among the major lid types, and occupy a narrower distribution in this dimensionality reduction (Fig. 7A, Fig. S14). Nevertheless, UMAP notoriously distorts distances between true clusters during dimensionality reduction (*105*), so we sought to further characterize the organization of ADK ESM-2 embeddings using quantitative metrics. To quantify the lid-type organization, we hierarchically clustered the embeddings based on euclidean distance (Fig. S16) and computed the adjusted mutual information score (AMI) by lid-type label (Fig. 7B). We found that while ESM-2 embedding clusters exhibit higher lid-type AMI than random, they retain less information than a one-hot encoded MSA (One-Hot MSA) clustered by Hamming distance (Fig. 7C, Fig. S17).

**Figure 7.**
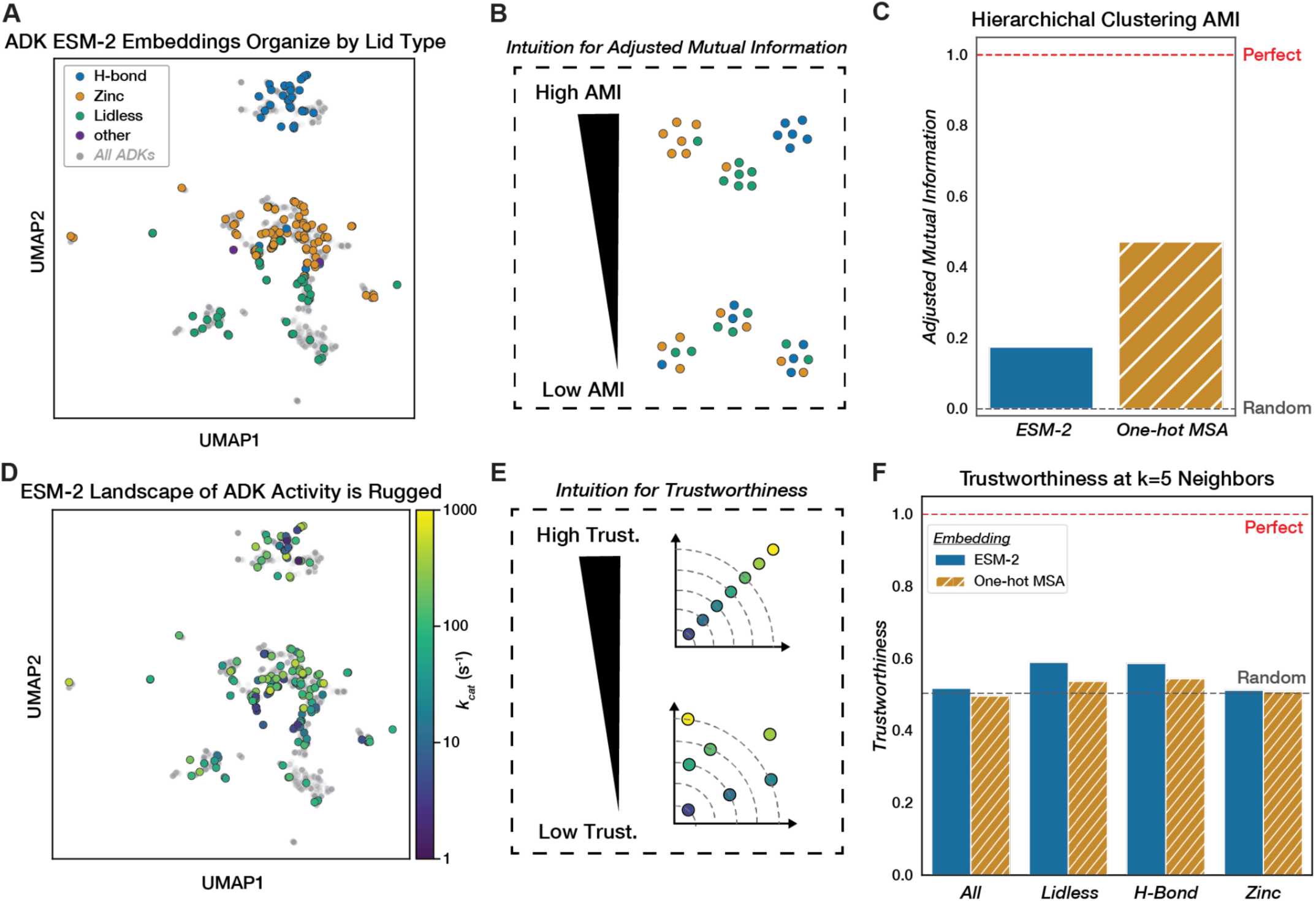
Evaluating the structural and catalytic organization of ADK sequence space learned by a protein language model. **(A)** UMAP of ADK ESM-2 embeddings, with all representative ADK sequences (∼5,000, Methods) plotted (gray). Sequences with measured catalytic parameters are encoded with color by lid type. **(B)** AMI reflects the agreement of a clustering method with respect to another label. In this case, low AMI would suggest poor clustering by Lid-type and vice versa. **(C)** Barplot comparing the AMI of hierarchical clustering on ESM-2 embeddings or a one-hot encoded MSA using euclidean and hamming distance, respectively, with respect to lid type. Perfect AMI (1.0) is plotted as a red dashed line, and random AMI (0.0) is plotted as a gray dashed line. **(D)** UMAP of ADK ESM-2 embeddings with *k_cat_* values encoded by color. Grey points represent all representative ADK sequences (∼5,000, Methods). **(E)** Trustworthiness quantifies the level of organization retained between two embeddings: in this case, it quantifies how similar neighboring sequences in ESM-2 space are in “*k_cat_* space”. **(F)** Grouped barplot of trustworthiness of ESM-2 ADK embeddings (by euclidean distance) and one-hot encoded MSA (by hamming distance) with respect to *k_cat_* computed at k=5 neighbors. Trustworthiness is computed for all sequences, as well as by lid type. Perfect trustworthiness is plotted as a red dashed line (*k_cat_* vs. *k_cat_*) and random as a gray dashed line (*k_cat_* vs. an average of 30 shuffles of corresponding embeddings).

We next investigated whether ESM-2 had instead learned a representation that reflects similarity in enzymatic activity. We find that the ESM-2 landscape appears visually rugged when colored by *k*_cat_ (Fig. 7D). Furthermore, quantifying *k*_cat_ organization using the continuous manifold metric of trustworthiness (Methods) reveals that both ESM-2 and One-Hot MSA exhibit essentially random organization in their nearest five neighbors with respect to *k*_cat_ (0.52 and 0.50 respectively) (Fig. 7F). Consequently, naturally occurring sequences close in the ESM-2 landscape often have vastly different *k*_cat_ values. Similar relationships apply to *K*_M_ and *k*_cat_/*K*_M_ (Fig. S18). The trustworthiness of ESM-2 embeddings relative to One-Hot MSA remains largely unchanged when the number of neighbors increases (Fig. S19). Given that ESM-2 has retained some lid-type organization, we also computed trustworthiness within lid types and found that this improves trustworthiness slightly (0.59 for Lidless and H-Bond) when not considering the global landscape (Fig. 7F). In a principal component analysis (PCA) on the ESM-2 embeddings (Fig. S20), we observed weak explained variance between the first principal component and *k_cat_* (r^2^ = 0.097) (Fig. S21). PC1 may reflect lid-type differences, as removing the significantly slower Lidless ADKs (Fig. 4C) weakens the regression (Fig. S21). Thus, while ESM-2 captures some high-level structural organization of ADKs, it fails to meaningfully encode the complex relationship between sequence and catalytic activity, perhaps because the co-evolutionary relationships learned are stronger for structure than for catalysis. These results highlight the limitations of current PLMs in predicting enzyme function from naturally occurring sequences alone.

### ADK k_cat_ and K_M_ prediction improve with increasing experimental data and supervision on top of PLMs

Supervised models built on top of pre-trained PLMs can enhance the prediction of mutational fitness effects compared to a zero-shot regime (*106–108*). However, fitness labels represent an aggregate of multiple underlying biochemical properties, such as catalytic activity and thermodynamic stability, and many models are specifically trained to predict the effect of a single mutation (*37*). Recently, there have been attempts to model sequence-catalysis relationships directly, with models like DLKcat (*109*) and TurNuP (*110*) trained on catalytic turnover rates collated from published literature in the BRENDA database (*79*). Our dataset of naturally occurring ADK sequences with measured *k*_cat_ values provides a valuable test set to evaluate the extent to which models like DLKcat have learned the sequence-*k*_cat_ relationship. When predicting *k*_cat_ from our sequences and substrate information (Methods), DLKcat performs poorly (spearman rho = −0.09, Fig. S22). We speculate that this poor performance may arise from challenges inherent to the BRENDA training data, collected under many experimental conditions, requiring scaling to standard temperature and pH. Furthermore, BRENDA has suffered data consistency issues from mis-annotations (*111*).

Apart from dataset inconsistencies, predicting *k*_cat_ for any naturally occurring enzyme sequence is inherently challenging, given the vast range of enzymatic activities (e.g., phosphorylation, hydrolysis, oxidation) and mechanisms. Instead, we considered whether we could use our newly collected experimental data to build an ADK-specific model that would be more predictive than a large model like DLKcat. We trained classic lightweight machine learning models–Random Forest (RF), Support Vector Regressor (SVR)–on increasing sub-samples of our dataset (20, 40, 60… 140 examples) and evaluated performance on a held-out 20% of our dataset for *k_cat_* and *K_M_* (Methods). For *k*_cat_ prediction, both SVR and RF outperform DLkcat with as few as 20 data points (mean RF spearman rho=0.14, mean SVR spearman rho=0.23, across 30 bootstrapped training set samplings). Performance steadily improved with increasing training data (Fig. 8A).

**Figure 8.**
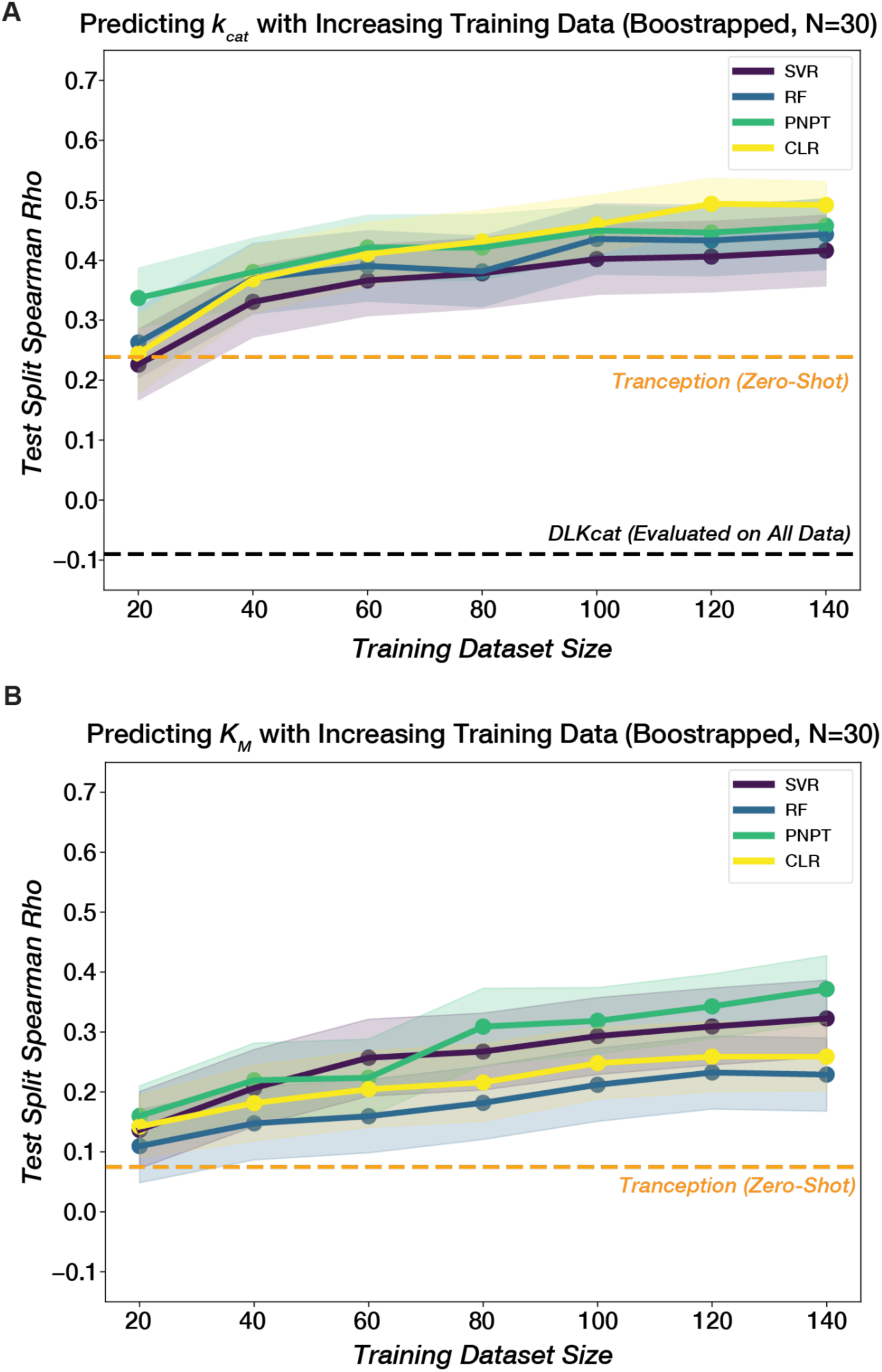
Improving *k_cat_* and *K_M_* prediction for ADK sequences with semi-supervised learning. The mean test spearman rho for different models across 30 samplings is plotted against training dataset size for (**A**) *k_cat_* and (**B**) *K_M_*. Models include Random Forest (RF), Support Vector Regression (SVR), ProteinNPT(PNPT) (*107*), and Convolutional Linear Regression (CLR) (*107*). For embeddings, SVR used a One-hot encoded MSA, RF used ESM-2 embeddings, and CLR and PNPT used Tranception(*112*) embeddings. Embeddings and other model hyperparameters were selected based on aggregate (mean) performance for both *k_cat_* and *K_M_* prediction. Shaded regions represent 95% confidence intervals across 30 training/test set samplings at each dataset size (Methods). A zero-shot evaluation of the Tranception PLM (*112*) is plotted as a dashed orange line. DLKcat (*109*) performance evaluated on all 175 sequences is plotted in (**A**) as a dashed black line.

Having demonstrated that our newly collected experimental data enables lightweight ML models to outperform an existing deep-learning model trained on large literature databases for predicting *k*_cat_, we next evaluated state-of-the-art semi-supervised approaches that combine pLMs with our experimental labels to further improve predictions. Specifically, we trained several variants of ProteinNPT (PNPT) (*107*), a pseudo-generative model that learns a joint representation of protein sequences and property annotations, as well as a convolutional linear regression (CLR) baseline(*107*). We experimented with different underlying pLM embeddings–ESM2 (*32*), Tranception (*112*), MSA Transformer (*113*)–and prediction targets–*k*_cat_ only, *K*_M_ only, or both plus growth temperature and lid type. ProteinNPT outperformed the lightweight regressors in all scenarios, particularly when leveraging Tranception embeddings, and its performance also improved when trained on a greater number of experimental labels (Fig. 8A-B). Additionally, the model variant predicting all targets simultaneously generally outperformed single-target models for *k*_cat_ and *K*_M_ (Fig. S23), demonstrating its ability to leverage relationships between different properties, as *k*_cat_ and *K*_M_ are correlated for naturally occurring ADKs (Fig. S24), a trend previously observed for mutants of ecADK (*48*).

## Conclusions and implications

We redeveloped an emerging microfluidic method, HT-MEK, to measure the catalytic constants *k*_cat_ and *K*_M_ for hundreds of naturally occurring and mutant ADK sequences. This dataset offered a unique view of the sequence-catalysis landscape on an evolutionary scale, revealing highly diverse catalytic parameters for extant ADKs. Analysis of this data uncovered a rugged topology with at least three global peaks, which remain navigable over long evolutionary timescales through distinct, path-dependent mechanisms, including single mutations and extra-dimensional bypasses. Our findings underscore the importance of examining sequence-catalysis landscapes on a broad scale. While small-scale comparisons have led to the longstanding hypothesis that adaptation to environmental temperature drives differences in catalytic rates, we show that thermophilic enzymes are not universally slower than their psychrophilic and mesophilic counterparts. A similar approach could be applied to assess the generality of other mechanistic and evolutionary hypotheses in enzymology.

PLMs predict protein structure by training on millions of protein sequences. Nevertheless, we show that enzymes with highly similar structures and analogous biochemical functions can exhibit vastly different catalytic parameters, hindering machine and deep-learning models that predict enzyme catalytic function from sequence alone. While current PLMs capture structural distinctions between naturally occurring variants of the same enzyme, they fail to accurately represent catalytic activity, perhaps because distinct structural solutions to high ADK activity have evolved over billions of years. This work underscores the need to integrate experimental data across a sequence-catalysis landscape with machine and deep-learning models to improve the prediction of catalytic parameters.

Supervised models trained on this study’s kinetic dataset outperform models using broad but sparse existing catalytic databases. Strategically selecting sequences to experimentally characterize catalytically for model training with Bayesian optimization or active learning approaches could improve the trade-off between model performance and training set size. Furthermore, we envision training semi-supervised generative models of sequence and function on data from many experimental assays at once to learn diverse enzyme sequence-catalysis landscapes. These models could generate novel sequences that potentially surpass the limits of enzyme activity that evolution has explored thus far.

## Acknowledgments

We thank members of the Pinney and Keiser laboratories, Siyuan Du, Stephanie Crilly, and Tony Capra for discussion and comments on the manuscript. We thank members of the Fordyce and Herschlag laboratories for helpful discussions and experimental advice. We acknowledge P. Suzuki for the design of the PS1.8K devices. D.F.M. thanks T. Bangalter, G.M. de Homem-Christo, D. Smith, and J. Vernon for additional support. **Funding:** This work was funded by a National Institutes of Health (NIH) grant (DP5OD033413), support from the Valhalla Foundation, and CZI grant DAF2018-191905 (DOI 10.37921/550142lkcjzw; M.J.K.) from the Chan Zuckerberg Initiative DAF, an advised fund of Silicon Valley Community Foundation (DOI 10.13039/100014989). D.F.M. was supported by a UCSF Discovery Fellowship. P.N. was supported by a Chan Zuckerberg Initiative Award (Neurodegeneration Challenge Network, CZI2018-191853). D.S.M. holds a Ben Barres Early Career Award from the Chan Zuckerberg Initiative as part of the Neurodegeneration Challenge Network (CZI2018-191853) and is supported by a NIH Transformational Research Award (TR01 1R01CA260415).

## Author contributions

D.F.M. M.M.P. and M.J.K designed the study. D.F.M. G.P.R.A. and J.A.P. cloned the ADK library. D.F.M. and G.P.R.A. performed *in vitro* experiments. D.F.M processed and analyzed kinetic data, and performed downstream computational, phylogenetic and statistical analysis. D.F.M. performed analyses of ESM-2 embeddings and machine learning experiments. P.N. performed deep-learning experiments with ProteinNPT with help from D.S.M. D.F.M. and M.M.P. wrote the manuscript with input from all authors.

## Competing interests

D.S.M. is an advisor for Dyno Therapeutics, Octant, Jura Bio, Tectonic Therapeutic and Genentech and a cofounder of Seismic.

## Data and materials availability

Summary tables of all measured kinetic parameters for each ADK variant are provided in the Supplementary Materials. All kinetic data generated in this study will be available in a registered Open Science Foundation repository upon publication. Code used for image processing and fitting kinetic parameters, as well as computational analyses will be made available in a public GitHub repository upon publication. Raw images from time-lapse microscopy will be made available upon request.

