## Supplementary Materials for "Evolutionary-Scale Enzymology Enables Biochemical Constant Prediction Across a Multi-Peaked Catalytic Landscape"

Contents:

- Materials and Methods
- Supplementary Text
- Tables S1-3
- Figs. S1-28

### Materials and Methods

#### Materials

All reagents were of the highest purity commercially available (≥97%). Any materials not specified below were purchased from Millipore-Sigma or ThermoFisher Scientific.

#### ADK Sequence Selection and Cloning and Mutagenesis

ADK amino-acid sequences were obtained through the Non-Redundant Protein Sequence Database (https://www.ncbi.nlm.nih.gov/refseq/about/nonredundantproteins/) for all Bacterial and Archaeal organisms with a reported optimal growth temperature from culture experiments (https://zenodo.org/records/1175609)(114). When multiple ADK sequences existed for a single organism, representative ADK sequences were selected on the basis of intra-genus pairwise sequence identity. Briefly, ADK sequences within a given genus were aligned all-by-all pairwise using Biopython.pairwise2.align.globalms with the following parameters: match=2, mismatch=-1, gap_open=-0.5, gap_extend=-0.1. For each sequence, the summed alignment score against all others within a genus was recorded. Sequences in the 1st and 99th length percentiles of all ADK sequences were excluded. After scoring, sequence selections were further refined by taking the max summed score sequence (see above) for an organism with a length within the fewest standard deviations of the mean sequence length for the corresponding genus. If multiple sequences had equal summed scores, the mode sequence was taken. The result of this workflow was 5,149 representative ADK sequences for bacteria and archaea.

Libraries of ADK sequences to assay on-chip were then selected to maximize diversity in sequence and growth temperature. Simple random sequence selection would bias our resulting library toward highly-sequenced organisms. To overcome this challenge and more evenly sample the phylogenetic tree of bacteria (see Fig. 1C), we used sci-kit learn to cluster representative ADK sequences based on the resulting alignment bitscores into four clusters, which were each clustered into four subclusters. 6 ADK sequences were randomly selected from each of the resulting 16 subclusters to yield 96 orthologous ADK sequences [Given biases towards mesophilic organisms in protein sequence databases, additional sequences in each cluster from psychrophiles (TGrowth ≤ 20 °C) and thermophiles (TGrowth ≥ 55 °C) were selected manually to ensure their representation]. A second batch of 96 orthologous ADK sequences was selected randomly from clusters based on hierarchical clustering on a one-hot encoded MSA or ESM-2 embeddings (using hamming and euclidean distance metrics, respectively). 192 genes encoding orthologous ADK sequences were then ordered from Twist Biosciences and cloned into a PurExpress vector containing a C-terminal eGFP tag connected by a ten amino acid serine/glycine linker (45) using the Gibson Assembly Protocol (New England Biolabs). Mutant constructs were generated either via Q5 Site-Directed Mutagenesis Kit (New England Biolabs) or ordered as a gene block and cloned with the Gibson Assembly Protocol as above.

Plasmids were purified with Monarch Plasmid Miniprep Kits (New England Biolabs) or Wizard SV 96 Plasmid DNA Purification Systems (Promega). All cloning and mutations were confirmed by Sanger sequencing miniprep DNA from NEB 5-alpha Competent E. coli (High Efficiency or Subcloning) cells (New England Biolabs) on an ABI3730xl capillary sequencer (Elim Biopharmaceuticals).

#### Adaptation of High-Throughput Microfluidic Enzyme Kinetics (HT-MEK) Platform and Specifications

We adapted the HT-MEK protocol (45) to collect kinetic data for Adenylate Kinase with the below-noted modifications.

##### *Photolithographic Mold Production.*

Briefly, flow and control layer mastermolds for PS1.8K MITOMI 2.0 PDMS devices containing 1792 chambers (https://www.fordycelab.com/microfluidic-design-files) were fabricated on 4” single-sided silicon test grade wafers (University Wafer) utilizing 30000 DPI transparency masks (Fineline Imaging) via the standard photolithography procedures employed for other MITOMI 2.0 device molds (45, 116–118).

##### *Fabrication of Two-Layer, Push-Down MITOMI Polydimethylsiloxane (PDMS) Devices*

PS1.8K PDMS (RTV615 010-Pail Kit Batch No.: 23BWFA015, Momentive) devices were fabricated as previously described for other two-layer, “push-down” MITOMI devices (Fig. S25)(45, 116, 117). 2” x 3” quartz slides (Electron Microscopy Sciences) were spin-coated (10s, 500 RPM, 75s 1820 RPM, using a Laurell Model WS-400B-6NPP/LITE) with PDMS at a 20:1 A:B ratio and baked at 80°C for 40 minutes. Quartz slides were chosen over glass due to their enhanced

UV light transmissibility for detecting NADPH fluorescence. All device fabrications were performed in a laminar flow cabinet to minimize dust contamination to our devices (AirScience Purair FLOW-36 Cabinet).

***Description of Device Architecture.***

The two-layer push-down MITOMI devices used herein (PS1.8K) contain 1792 individual chambers in a 32 x 56 array (Fig. S25). Integrated pneumatic valves enable chambers and chamber regions to be isolated for user-defined intervals. Each chamber is divided into two compartments: the DNA compartment that contains an arrayed spot of plasmid DNA and the reaction compartment where the expressed eGFP-tagged enzyme is pulled-down with a neutravidin-biotin-anti-eGFP nanobody linkage. These chambers are separated by a valve ("Neck") that isolates the plasmid DNA until the enzyme expression step (see Surface Patterning and On-Chip ADK Expression below). Each DNA compartment within every chamber contains a specific, sequence-verified and user-defined ADK-eGFP variant or buffer as a negative control. (see ADK Plasmid Arraying and Device Alignment below). To isolate chambers from one another and to prevent mixing between chambers during expression or the enzymatic reaction, chambers are separated by an additional valve ("Sandwich"). Finally, a third valve ("Button") protects a circular patch in the center of the reaction chambers. This valve allows for surface patterning that facilitates on-chip protein immobilization and purification, followed by the concurrent initiation of sequential on-chip reactions throughout the device (see Activity Measurement on Adapted HT-MEK Platform, below).

***ADK Plasmid Arraying and Device Alignment.***

5-10 µL of sequence-verified ADK plasmids were transferred to a 384-well plate (50-400 ng/µL), dried down at 60 °C on a hot plate, and resuspended in 20 µL of "print buffer" (5 mg/mL BSA, 12.5 mg/mL Trehalose-Dihydrate, 75mM NaCl). Plasmids were randomly assigned positions within a 32 x 56 array (code available upon request), with controls containing only print buffer distributed throughout (typically, 7-10 replicates of each plasmid variant were included in each array). 360-400 pL spots of ADK plasmids were arrayed on PDMS-coated quartz slides (see above) using a SciFLEXARRAYER S3 (Scieneon) outfitted with a PDC-70 Piezo Dispense Capillary nozzle (catalog no. P-2030, Scienion) with a Type 4 PDC coating (catalog no. S-6041, Scienion). Between every sample, the nozzle was washed using the S3 Software wash_flush_narrow method to ensure no cross-contamination. To prevent clogging, the nozzle was washed using 0.1% sciCLEAN 8 solution (catalog no. C-5283, Scieneon) in ultrapure water every 20 samples. Consistent with minimal cross-contamination, control chambers containing only buffer spots typically displayed no discernible eGFP fluorescence or ADK activity above background (Fig. 2B). Fabricated devices were then aligned on top of this plasmid array by eye, placing one plasmid spot per DNA chamber, using a standard dissecting microscope. Devices assembled around plasmid libraries were baked on a hot plate (Torrey Pines) for 8 hrs at 80 °C.

***Device Valve Manipulation and Optics.***

Integrated valves within the microfluidic devices were manipulated via a custom-built pneumatic manifold and controlled through a custom Python codebase (manifold design and code available upon request). Time-lapse inverted fluorescence microscopy images (see ADK Activity Measurement on Adapted HT-MEK Platform, below) were taken on a Ti-2E Large Field of View Inverted Fluorescence Microscope (Nikon) outfitted with a sCMOS camera (Photometrics Kinetix Scientific CMOS). NADPH images were taken with a solid-state light source (312 mW, configured at 15% intensity) outfitted with a 340 nm output with an integrated bandpass filter (Lumencor RETRA Light Engine) and a FURA filter set (FF01-510/84-25 bandpass emission filter; FF409-Di03 dichroic beamsplitter) using 500 ms exposure times. eGFP images were taken using a Lumencor Spectra Light Source using the GFP/FITC output (475/28 nm; 500 mW) and a GFP/FITC/CY2 filter set (FF01-466/40 bandpass excitation filter; FF495-Di03 dichroic beamsplitter, and FF03-525/50 bandpass emission filter) using 5 ms exposure times.

***Surface Patterning and On-Chip ADK Expression***

Surface functionalization was carried out as previously described with several modifications (45). Initial flow and control line pressures were set to 3-4 psi and 40 psi, respectively, and Neck and Button valves were initially closed to prevent premature DNA solubilization and to protect the contact surface of the Button valves (Fig. S25-26). To passivate the PDMS on the walls, ceiling, and floor of the device chamber (excluding the contact surface of the Button valve), a 5 mg/mL solution of ultra-pure bovine serum albumin (BSA; Invitrogen, AM2618) was flowed through all inlet channels for 30 minutes (keeping Neck and Button Valves pressurized). All channels were then washed with 1X phosphate-buffered saline (PBS; Corning 21-040-CM) for 30 minutes, again keeping the Neck and Button Valves pressurized. To specifically modify surfaces on and under the Button valve to pull down expressed eGFP-tagged ADK variants, we introduced 1 mg/mL of neutravidin (NA, Thermo Scientific catalog no. PI31050) for 55 minutes with the Button valves open to capture NA specifically on the contact surfaces of the Button valve. We then closed Button valves to protect NA functionalized surfaces and washed out unbound NA with 1X PBS for 30 minutes. To quench any NA nonspecifically bound to chamber and channel walls, we washed with a 2 mg/mL solution of biotinylated BSA (bBSA; ThermoFisher Pierce catalog no. 29130) for 30 minutes with Button valves closed. Again, unbound bBSA was washed out by flowing 1X PBS for 30 minutes. Finally, we depressurized the Button valve and introduced a variable solution of a biotinylated anti-eGFP nanobody (ProteinTech, catalog no. GTB-250) and bBSA. For binding assays requiring high amounts of pulled-down product, we flowed on a 100 µg/mL solution of biotinylated anti-eGFP nanobody (ProteinTech catalog no. GTB-250) for 15 minutes. For kinetic assays, the effective concentration of pulled-down eGFP-ADK was reduced by including 500 µg/mL (final concentration) of bBSA along with 50 µg/mL anti-eGFP nanobody during this step (see Activity Measurement on Adapted HT-MEK Platform, below). Finally, Button valves were pressurized again to protect the pulled-down anti-eGFP nanobody, and any unbound nanobody was washed out with 1X PBS for 30 minutes. The surface patterning procedure ultimately produced an NA-biotin-anti-eGFP nanobody stack ("pedestal") exclusively on the contact surface of the Button valve capable of pulling down expressed eGFP-tagged ADK variants (Fig. 2B).

ADK-eGFP constructs were expressed "on-chip" via in vitro transcription and translation (IVTT). We utilized a reconstituted IVTT system, PurExpress (New England Biolabs), supplemented with RNAse-inhibitor (Promega, N2511), Ultra Pure Water (Invitrogen, 10977-015), and 50 uM zinc chloride. This mixture was then flowed onto the chip with Button valves closed and Neck valves closed for 3 minutes to replace existing buffers, with 2 additional minutes of flow with Button valves open to increase chip volume. Then Sandwich valves were closed to prevent mixing between chambers, Necks were opened to allow the IVTT machinery to access the plasmid in the neighboring DNA chamber, and Button valves were toggled, then closed to encourage the DNA chamber to fill completely. Devices were incubated at 37 °C on a hot plate (Torrey Pines) for 1-2 hrs. After 1 hour of incubation, Button valves were opened to begin capturing expressed ADK-eGFP variants with the anti-eGFP functionalized pedestal described above. Expression was visualized by imaging the device in the eGFP channel (see Device Valve Manipulation and Optics above). ADK-eGFP variants were purified by flowing ADK assay buffer (50 mM HEPES 100 mM potassium chloride pH 8.0) over the whole device with Button valves closed to remove expression components and any uncaptured ADK-eGFP. ADK-eGFP concentrations were calculated following the procedure of Markin et al., which utilizes an eGFP standard curve to relate eGFP intensities to eGFP concentration (45).

**Activity Measurement on Modified HT-MEK Platform**
ADK activity was assayed in the direction of ATP formation coupled to the production of NADPH using an excess of the enzymes Glucose-6-phosphate Dehydrogenase from Leuconostoc mesenteroides (Sigma-Aldrich, G5885) and Hexokinase from Saccharomyces cerevisiae (Sigma-Aldrich, H4502) (62). Reaction progress was monitored by a gain in NADPH fluorescence over time (see Device Valve Manipulation and Optics above). Fluorescence intensities were related to NADPH concentration using a NADPH standard curve collected for each chamber at the outset of each on-chip experiment. Substrate (ADP) was titrated two-fold from 15.625uM to 4 mM, and the activity at each concentration was monitored using time-lapse imaging over 30 minutes (see Device Valve Manipulation and Optics, above). Before initiating reactions, the substrate solution was flowed onto the device for 5 minutes to completely replace the existing solution within the device, keeping the Button valves closed to prevent enzyme turnover. To commence the reactions, the Sandwich valves were sealed to isolate the chambers and prevent product leakage, and then the Button valves were opened simultaneously for all chambers, allowing the immobilized enzyme to come into contact with the substrate, thereby initializing the enzymatic reaction in all chambers at the same time.

The buffer for all assays and standard curves was 50 mM HEPES 100 mM potassium chloride pH 8.0. All microscopy image data was processed with a custom codebase (available upon request). Briefly, rastered images (25 per timepoint, in a 5x5 grid) were stitched together to obtain a complete whole-chip-image (WCI). eGFP and NADPH quantification WCIs were background-subtracted using images taken with identical settings immediately prior to expression initiation (in the case of eGFP) or prior to the standard curve and kinetic assays (in the case of NADPH). Individual chambers and buttons were identified by manually selecting upper-leftmost, upper-rightmost, lower-leftmost, and lower-rightmost chamber centers and then interpolating the rough position of the 32x56 array. These rough positions were refined using opencv.HoughCircles, and then summed pixel intensity was obtained across all images within each chamber or button, then converted to [NADPH] (uM) or [eGFP] (nM) using the appropriate standard curve. To obtain kcat and KM values, initial rates were determined from the processed kinetic data and fit to the Michaelis-Menten equation using SciPy (see code).

Activity parameters were fit individually for each chamber on the microfluidic device. Lower quartile background rates from buffer (empty) chambers were subtracted from each initial rate at the appropriate substrate concentration to isolate the enzyme-catalyzed rate of reaction. The mean and standard deviation across chamber (biological) replicates of each unique sequence were reported (typically, the standard deviation of kcat values between biological replicates was ≤20% of the mean; Fig. S27). A lower limit of detection for enzyme concentration was set to 0.5 nM, and data from chambers with ADK expression below this threshold were not analyzed further. kcat values from well-behaved orthologous ADKs in the literature (Bacillus subtilis, Geobacillus stearothermophilus, Jeotgalibacillus marinus, and Escherichia coli), measured under similar conditions in the same reaction direction (ATP formation) (51, 54) were used to further scale fit VMax values to convert eGRP RFUs to concentration (nM) for other orthologs. For the mutant on-chip experiment, corresponding wild-type sequences were included as cross-chip internal standards and used to normalize kcat values to ensure consistency with the full wild-type dataset (r2 = 0.92 for Vmax normalized by eGFP between wild-type sequences in ortholog and mutant on-chip experiments). KM values outside of the range of [ADP] used in our experiments were flagged and reported as upper or lower "limits" of this value. C. tepidum ADK was selected as a candidate for mutagenesis experiments, but as we consistently observed KM values above the limit of our assay, it was excluded from downstream analyses.

On-chip ADK kinetic values were found to be highly technically reproducible between different on-chip experiments collected on different days (Fig. S28; R2 = 0.960 between independent replicates), and relative kcat values correlate well with orthologous ADKs previously reported in the literature (Fig. 2E).

**Urea Challenge of ADK Activity**

The effect of urea on ADK activity was determined in a single high-throughput experiment on an identical library print as the Michaelis-Menten ortholog experiment (see Surface Patterning and On-Chip ADK Expression and Activity Measurement on Modified HT-MEK Platform above). Activity measurements were performed using the same coupled reaction scheme as described above, with increasing amounts of urea (0, 0.5, 1, and 2M) and saturating amounts of ADP (4 mM) in each condition. Off-chip controls confirmed that the coupled enzymes maintain their activity at these low urea concentrations. Loss of activity was defined as no overlap between mean and 95% confidence interval (across biological replicates) from subsequent urea conditions.

**ATP-647 Binding On-Chip**

Binding measurements of a fluorescent analog of ATP, AlexaFluor 647 ATP (ATP-647; ThermoFisher, Catalog No. A22362), were made in a single high-throughput experiment on an identical library print as the Michaelis-Menten ortholog experiment (see Surface Patterning and On-Chip ADK Expression and Activity Measurement on Modified HT-MEK Platform above) with the following modification: during patterning, the anti-eGFP nanobody was not diluted with bBSA, so as to allow for higher amounts pulled-down enzyme and greater binding signal from the fluorescent probe. ATP-647 was diluted in 50 mM HEPES 100 mM potassium chloride pH 8.0 to make a two-fold dilution series from 7.8 µM to 1000 µM. Increasing concentrations of ATP-647, starting at 0M, were flowed on the device to chambers with Buttons down and Necks closed for 3 minutes. Sandwiches were then closed, and Buttons were opened to allow for ATP-647 binding to ADK variants for 5 minutes. Buttons were then closed, and excess probe was washed out with buffer for 5 minutes prior to imaging in both eGFP and CY5 (ex/em: 618/698 nm) channels. Afterward, the next concentration of ATP-647 was introduced, and the process iterated.

**ADK Off-Chip Expression and Kinetics**

Variants were expressed off-chip following the PurExpress protocol (New England Biolabs), with zinc and RNAse-inhibitor added as described above. Expressed protein was exchanged into the previously described HEPES/KCl buffer using Amicon Ultra 0.5mL 10kDa Centrifugal Filters. ADK-eGFP concentration was determined on a BioTek Synergy H1 Plate Reader (Agilent) using an eGFP standard curve (Proteintech, EGFP-250?) (ex/em: 490/510). The same activity assay from on-chip experiments was used off-chip, and the gain of fluorescence from NADPH production was monitored via plate reader (ex/em: 350/450). kcat and KM values were obtained by fitting measured initial rates to the Michaelis-Menten equation using SciPy and averaged across two technical replicates.

**Computational Analyses and Machine Learning**

***Generation of Taxonomic Trees and Multiple Sequence Alignments***

The ADK taxonomical tree was obtained by inputting corresponding NCBI Taxonomy IDs into PhyloT (https://phylot.biobyte.de/terms.cgi) and then visualized in ITOL (https://itol.embl.de/)(119). Protein Multiple Sequence Alignments were generated via ClustalOmega (https://www.ebi.ac.uk/jdispatcher/msa/clustalo) using default parameters.

***AlphaFold2 Structure Prediction***
AlphaFold2 models were generated using the ColabFold(120) alphafold2_batch.ipynb notebook with msa_mode=MMseqs2 (UniRef + Environmental) and default parameters, and top-ranked models based on pLDDT were used in subsequent analyses.

***Phylogenetic Correlation***
Phylogenetic signal was computed using the phylosignal R package(64). The input phylogenetic tree was created using MEGA(121) based on maximum-likelihood from a MSA (described above: Generation of Taxonomic Trees and Multiple Sequence Alignments).

***ADK Isoform Assignment***
Lid types for ADK (Lidless, Zinc Lid, H-Bond Lid, Other) were assigned based on the identity of a four residue motif within the multiple sequence alignment of our ADK sequence library. The classic motifs are CCCC and HSDT for Zinc-Lid and H-Bond, respectively. After this motif was extracted from the MSA for each sequence, Lid Types were assigned with the following heuristic: 3 or more cysteines and 0 gaps = Zinc Lid, 1 or fewer cysteines and 0 gaps = H-Bond Lid, 2 or more gaps = Lidless, and any remaining were labeled "Other".

***Generation of Edit Distance Minimum Spanning Tree***
Sequence graphs and minimum spanning trees were generated in Python and NetworkX (see code). Sequence graphs contained a node corresponding to each ADK sequence and were fully connected with edges weighted by the number of amino-acid differences (including gaps) between sequences in an MSA.

***ADK Sequence Embedding with ESM-2***
Protein language model embeddings from ESM (https://github.com/facebookresearch/esm) were obtained by inputting ADK sequences in .fasta format to the esm-extract command(32). The esm2_t33_650M_UR50D was used, and fixed-length (mean-pooled) embedding vectors (L=1280) were obtained from sequence-averaged tokens.

***Clustering and Manifold Metrics***
Hierarchical clustering of ESM-2 embeddings was performed on a pairwise euclidean distance matrix of the embeddings using SciPy's hierarchy.fcluster function, with criterion="maxclust" and t=30. A similar procedure was used for one-hot-encodings, but instead, a pairwise hamming distance matrix was used as an input to fcluster. Adjusted mutual information and trustworthiness scores for clusters and embeddings, respectively, were both implemented with Sci-Kit Learn.

***Machine Learning***
Random Forest and Support Vector Regression models were imported from Sci-Kit Learn. Input features were one-hot encodings of ADK sequences from an MSA, or ESM-2 embeddings (see ADK Sequence Embedding with ESM-2), and output labels (Mean kcat and KM) were log10 transformed. To gain a robust sense of model performance on unseen sequences in our low-N data regime, we performed the dataset ablation experiment in a modified bootstrap procedure. Briefly, we split our dataset into 80:20 train:test splits 30 times, with different random seeding each time. Within each split, we sampled smaller training subsets in increments of 20 (i.e., 20, 40, 60, etc.) one time per split. Models were then fitted on each subset and evaluated on the corresponding test set, reporting the Spearman Correlation. The performance metrics were averaged for each subset size across the 30 train:test splits. The DLKcat baseline was obtained by inputting unaligned ADK sequences and the Canonical SMILES for ADP from PubChem into the published model, as described in https://github.com/SysBioChalmers/DLKcat/tree/master/DeeplearningApproach (109). The performance of DLKcat was evaluated as Spearman correlation between true values (unnormalized) and predicted output of DLKcat for all 175 ADK sequences.

***Deep-Learning with ProteinNPT***
Following the methodology outlined in Notin et al. (107), we trained various ProteinNPT model variants to jointly learn the distribution of ADK sequences and their associated catalytic activity parameters (kcat, KM). We evaluated several protein

sequence embedding models, including ESM2 (650M), MSA Transformer, and Tranception. Our experiments revealed that Tranception yielded optimal performance for both kcat and KM prediction (Fig. S23).

ProteinNPT's architecture readily accommodates multi-property prediction by jointly embedding each target of interest with its corresponding sequence. Our ablation studies demonstrated superior performance with a model exposed to all available annotation types (kcat, KM, growth temperature, and lid type) compared to models focused on single targets or subsets of the four possible targets. This improvement was particularly pronounced for KM prediction (Fig. S23) All other training hyperparameters strictly adhered to the defaults specified in Notin et al. (107).

**Supplementary Text S1.**

***Catalytic differences are independent of enzyme foldedness.***

Considering the temperatures to which our orthologous sequences are adapted, a subset (21/175) has $T_{Growth}$ values below our assay temperature of 25°C, prompting questions about their foldedness during our assays. Nevertheless, studies on psychrophilic (cold-loving) enzymes have shown that many remain stable at temperatures significantly above their host organisms' $T_{Growth}$ value, with activity often measured near 25°C (*54*, *65*, *84*, *122–127*). This excess stability may buffer against transient temperature increases in their environments. We conducted several experiments to test whether our ADK orthologs were natively folded on-chip. First, we tested whether our ADKs had well-formed ATP binding sites, as expected for a natively-folded ADK, by assessing the binding of a fluorescent ATP analog, ATP-Alexa Fluor 647 (ATP-647). We performed ATP-647 titration experiments on-chip and observed dose-dependent binding of ATP-647 (Fig. S3, which could be competed off with the two-substrate mimicking inhibitor, P1,P5-bis(adenosine-5'-)pentaphosphate (AP5) (Fig. S4). Most of our ADK orthologs show a binding signal for ATP-647 (Fig. S5). Nevertheless, the subset of ADK orthologs that did not show ATP-647 binding still exhibited ADK activity above background (Table S2). This suggests that these orthologs have well-folded ATP binding sites, but factors such as fluorophore quenching upon binding or steric hindrance of the larger ATP-647 relative to ATP may have prevented the observation of ATP-647 binding for these variants. Second, nearly all (91%) of the orthologs in our library are faster than the consensus sequence for the family, suggesting these naturally occurring ADKs are not only folded but contain activity-enhancing residue combinations that are uncoupled in the consensus sequence (Fig. 3A, 3B; see also *Contributions from multiple peaks catalytically impair the ADK consensus sequence*). Third, for the ADK orthologs in our library that have been previously characterized in the literature under common conditions and assays, we observe a strong correlation between those $k_{cat}$ measurements and our on-chip measurements ($r^2 = 0.96$, Fig. 2E), which includes the psychrophilic ortholog from *J. marinus* ($T_{growth}$ = 11 °C, $T_M$ = 45 °C) (*54*). Finally, we conducted activity measurements in the presence of the denaturant urea to test if a variant was already partially unfolded without urea, as small amounts of denaturant would cause substantial additional unfolding and proportional decreases in activity. Supporting the foldedness of our orthologs, none of the psychrophilic ADK orthologs lost activity upon the addition of 0.5 M urea (Fig. S6). Considering all ADK orthologs, only six lost activity in 0.5 M urea, and just 14 orthologs lost activity in 1.0 M urea (Fig. S7, Methods). Taken together, these results strongly support the native foldedness of these diverse ADK orthologs on-chip.

### Supplementary Tables

#### Table S1. Median Optimal Growth Temperature for 193 Organisms Associated with ADK Library

| organism_name | growth_temperature |
|---|---|
| desulfovibrio_desulfuricans | 31 |
| azorhizobium_caulinodans | 30 |
| pseudarthrobacter_chlorophenolicus | 27 |
| leptospira_interrogans | 29 |
| thermotoga_maritima | 75 |
| pseudothermotoga_lettingae | 65 |
| anaeromyxobacter_dehalogenans | 28 |
| thermococcus_gammatolerans | 88 |
| erwinia_tasmaniensis | 28 |
| shewanella_halifaxensis | 16 |
| aliivibrio_salmonicida | 15 |
| histophilus_somni | 37 |
| pseudomonas_syringae | 26 |
| pseudomonas_entomophila | 28 |
| lactobacillus_delbrueckii | 37 |
| streptococcus_pneumoniae | 37 |
| geobacillus_thermodenitrificans | 58 |
| bacillus_licheniformis | 32 |
| heliomicrobium_modesticaldum | 50 |
| desulfitobacterium_hafniense | 37 |
| syntrophomonas_wolfei | 36 |
| symbiobacterium_thermophilum | 60 |
| halobacterium_salinarum | 37 |
| psychromonas_ingrahamii | 10 |
| methylocella_silvestris | 23 |
| bartonella_henselae | 37 |
| thermobifida_fusca | 50 |
| corynebacterium_urealyticum | 37 |
| kosmotoga_olearia | 65 |
| petrotoga_mobilis | 55 |
| thermococcus_kodakarensis | 60 |
| thermococcus_sibiricus | 81 |
| shewanella_frigidimarina | 21 |
| pseudoalteromonas_atlantica | 21 |
| actinobacillus_pleuropneumoniae | 37 |
| legionella_pneumophila | 37 |
| acinetobacter_baylyi | 31 |
| chlorobaculum_parvum | 25 |
| streptococcus_thermophilus | 38 |
| lactococcus_lactis | 31 |
| lysinibacillus_sphaericus | 33 |
| clostridium_botulinum | 37 |
| methanocella_arvoryzae | 45 |
| thermodesulfovibrio_yellowstonii | 60 |
| moorella_thermoacetica | 59 |
| natranaerobius_thermophilus | 50 |
| shewanella_loihica | 21 |
| alteromonas_mediterranea | 23 |
| sorangium_cellulosum | 28 |
| elusimicrobium_minutum | 26 |
| paramagnetospirillum_magneticum | 30 |
| gluconacetobacter_diazotrophicus | 30 |

| | |
|---|---|
| fervidobacterium_nodosum | 70 |
| thermosipho_melanesiensis | 70 |
| pyrococcus_furiosus | 96 |
| chlamydia_trachomatis | 37 |
| colwellia_psychrerythraea | 9 |
| vibrio_cholerae | 36 |
| leptothrix_cholodnii | 23 |
| polaromonas_naphthalenivorans | 20 |
| chlorobaculum_tepidum | 46 |
| alkalilimnicola_ehrlichii | 29 |
| enterococcus_faecalis | 36 |
| limosilactobacillus_reuteri | 36 |
| methanosarcina_acetivorans | 37 |
| geotalea_daltonii | 30 |
| archaeoglobus_fulgidus | 81 |
| alkaliphilus_oremlandii | 29 |
| caldanaerobacter_subterraneus | 69 |
| salinispora_arenicola | 27 |
| marinobacter_nauticus | 29 |
| bacillus_subtilis | 32 |
| deinococcus_deserti | 28 |
| xanthomonas_campestris | 26 |
| sphingopyxis_alaskensis | 28 |
| thermotoga_neapolitana | 79 |
| thermosipho_africanus | 68 |
| bdellovibrio_bacteriovorus | 28 |
| mycoplasma_mobile | 26 |
| mycoplasma_mycoides | 37 |
| photobacterium_profundum | 13 |
| vibrio_parahaemolyticus | 29 |
| bordetella_petrii | 34 |
| neisseria_gonorrhoeae | 37 |
| methylococcus_capsulatus | 41 |
| cellvibrio_japonicus | 37 |
| bacillus_cereus | 30 |
| geobacillus_kaustophilus | 55 |
| methanococcoides_burtonii | 21 |
| trichlorobacter_lovleyi | 26 |
| caldicellulosiruptor_saccharolyticus | 68 |
| caldicellulosiruptor_bescii | 73 |
| streptomyces_coelicolor | 26 |
| haloarcula_marismortui | 37 |
| geobacillus_stearothermophilus | 54 |
| escherichia_coli | 37 |
| sphaerochaeta_pleomorpha | 25 |
| parafannyhessea_umbonata | 37 |
| deinococcus_proteolyticus | 30 |
| spirulina_subsalsa | 23 |
| rhodanobacter_fulvus | 27 |
| dietzia_alimentaria | 28 |
| bartonella_quintana | 37 |
| porphyromonas_macacae | 37 |
| mycoplasmopsis_arginini | 37 |
| thermodesulfatator_atlanticus | 70 |
| acidimicrobium_ferrooxidans | 45 |
| thermomicrobium_roseum | 70 |

| | |
|---|---|
| mesomycoplasma_flocculare | 37 |
| prochlorococcus_marinus | 20 |
| acaryochloris_marina | 20 |
| asanoa_hainanensis | 28 |
| lachnospira_multipara | 38 |
| halorhodospira_halophila | 25 |
| fervidobacterium_islandicum | 63 |
| pseudobdellovibrio_exovorus | 29 |
| oleiphilus_messinensis | 27 |
| virgibacillus_dokdonensis | 30 |
| pyrinomonas_methylaliphatogenes | 60 |
| effusibacillus_pohliae | 55 |
| desulfatibacillum_alkenivorans | 30 |
| mycoplasmopsis_canis | 37 |
| kiritimatiella_glycovorans | 28 |
| actinocatenispora_sera | 28 |
| mesomycoplasma_hyorhinis | 37 |
| mycoplasmoides_genitalium | 37 |
| agromyces_subbeticus | 29 |
| dactylosporangium_aurantiacum | 29 |
| streptomyces_sampsonii | 28 |
| mesonia_phycicola | 28 |
| acetomicrobium_flavidum | 53 |
| treponema_putidum | 37 |
| allofrancisella_guangzhouensis | 30 |
| sporolituus_thermophilus | 55 |
| ferrimicrobium_acidiphilum | 32 |
| streptomonospora_alba | 30 |
| pseudoroseomonas_rhizosphaerae | 30 |
| gemmobacter_aquatilis | 30 |
| oscillatoria_acuminata | 22 |
| glaciecola_punicea | 10 |
| microbacterium_chocolatum | 30 |
| veillonella_seminalis | 37 |
| chitinophaga_pinensis | 23 |
| flavobacterium_hydatis | 25 |
| nostoc_punctiforme | 20 |
| salinibacter_ruber | 37 |
| helicobacter_acinonychis | 37 |
| rubinisphaera_brasiliensis | 30 |
| falsiroseomonas_stagni | 30 |
| chondromyces_apiculatus | 29 |
| thermodesulfobacterium_commune | 70 |
| acetohalobium_arabaticum | 37 |
| streptomyces_subrutilus | 27 |
| methylophaga_frappieri | 29 |
| lactobacillus_crispatus | 37 |
| desulfurella_acetivorans | 55 |
| marinimicrobium_agarilyticum | 28 |
| capnocytophaga_ochracea | 37 |
| megasphaera_paucivorans | 30 |
| pedobacter_hartonius | 13 |
| oenococcus_oeni | 30 |
| brucella_vulpis | 37 |
| desulfurobacterium_thermolithotrophum | 70 |
| synechococcus_elongatus | 24 |

| | |
|---|---|
| legionella_erythra | 37 |
| verminephrobacter_aporrectodeae | 22 |
| rubrobacter_radiotolerans | 37 |
| patulibacter_medicamentivorans | 28 |
| brachyspira_pilosicoli | 37 |
| nonlabens_ulvanivorans | 22 |
| bombiscardovia_coagulans | 37 |
| desulfurobacterium_atlanticum | 70 |
| dissulfuribacter_thermophilus | 60 |
| kushneria_avicenniae | 29 |
| salinispira_pacifica | 35 |
| adlercreutzia_mucosicola | 37 |
| prosthecobacter_debontii | 26 |
| mycoplasmoides_pneumoniae | 37 |
| fibrobacter_intestinalis | 37 |
| bryobacter_aggregatus | 26 |
| salipiger_mucosus | 30 |
| rickettsiella_massiliensis | 28 |
| persephonella_hydrogeniphila | 70 |
| hydrogenobacter_thermophilus | 70 |
| micromonospora_viridifaciens | 27 |
| chrysiogenes_arsenatis | 28 |
| paenibacillus_bovis | 30 |
| eggerthia_catenaformis | 37 |
| rubritalea_squalenifaciens | 28 |
| faecalibaculum_rodentium | 37 |
| rubritepida_flocculans | 50 |
| helicobacter_rodentium | 37 |
| jeotgalibacillus_marinus | 11 |

### Table S2. Catalytic Parameters for ADK Orthologs

| Organism | k_cat (s^-1) | K_M (uM) | k_cat/K_M (s^-1 M^-1) | Replicates |
|---|---|---|---|---|
| thermobifida_fusca | 16.0 +/- 2.0* | 20.9 +/- 4.0* | 764964.3 +/- 175366.2* | 6 |
| rubrobacter_radiotolerans | 8.6 +/- 2.1 | 35.1 +/- 16.5 | 245481.9 +/- 130353.5 | 7 |
| porphyromonas_macacae | 42.4 +/- 10.6 | 50.2 +/- 11.0 | 845385.4 +/- 281236.1 | 6 |
| thermomicrobium_roseum | 11.8 +/- 1.7 | 74.6 +/- 23.5 | 157827.6 +/- 55003.0 | 8 |
| synechococcus_elongatus | 14.7 +/- 2.3 | 79.4 +/- 18.1 | 185677.7 +/- 51308.7 | 7 |
| cellvibrio_japonicus | 18.4 +/- 1.4 | 93.7 +/- 19.7 | 196471.7 +/- 43973.7 | 6 |
| streptomyces_subrutilus | 18.1 +/- 3.0 | 108.4 +/- 26.2 | 166872.3 +/- 48829.4 | 6 |
| acinetobacter_baylyi | 61.2 +/- 14.6 | 112.0 +/- 16.0 | 546705.0 +/- 151968.1 | 6 |
| thermococcus_sibiricus | 6.3 +/- 4.2 | 113.3 +/- 90.8 | 55495.3 +/- 57718.7 | 6 |
| bartonella_quintana | 8.0 +/- 1.5 | 116.0 +/- 39.4 | 69183.4 +/- 26635.5 | 6 |
| syntrophomonas_wolfei | 70.2 +/- 6.8 | 123.0 +/- 15.1 | 570663.6 +/- 89258.8 | 7 |
| deinococcus_proteolyticus | 76.1 +/- 8.3 | 129.8 +/- 15.0 | 586601.7 +/- 92971.5 | 7 |
| desulfurobacterium_atlanticum | 81.7 +/- 7.3 | 130.0 +/- 10.2 | 628982.7 +/- 74821.8 | 6 |
| desulfurella_acetivorans | 34.7 +/- 2.9 | 132.1 +/- 26.8 | 262304.7 +/- 57567.2 | 6 |
| acetohalobium_arabaticum | 14.6 +/- 3.6 | 133.9 +/- 55.2 | 109071.3 +/- 52302.6 | 7 |
| eggerthia_catenaformis | 14.5 +/- 1.4 | 134.7 +/- 28.9 | 107873.7 +/- 25340.1 | 7 |
| desulfurobacterium_thermolithotrophum | 65.5 +/- 3.5 | 157.3 +/- 37.0 | 416536.6 +/- 100508.8 | 8 |
| pseudarthrobacter_chlorophenolicus | 81.6 +/- 8.0 | 158.7 +/- 23.6 | 514482.4 +/- 91604.8 | 8 |
| ferrimicrobium_acidiphilum | 34.8 +/- 6.2 | 162.8 +/- 20.5 | 213718.9 +/- 46645.5 | 6 |

| | | | | |
|---|---|---|---|---|
| haloarcula_marismortui | 1.1 +/- 0.4 | 175.5 +/- 133.3 | 6180.1 +/- 5192.3 | 4 |
| brucella_vulpis | 60.8 +/- 5.6 | 176.5 +/- 45.9 | 344431.8 +/- 94953.2 | 9 |
| jeotgalibacillus_marinus | 26.0 +/- 1.6 | 176.5 +/- 33.9 | 147113.7 +/- 29641.6 | 5 |
| alteromonas_mediterranea | 39.8 +/- 11.6 | 176.9 +/- 28.6 | 225226.2 +/- 75107.2 | 7 |
| streptomyces_coelicolor | 60.5 +/- 10.4 | 182.4 +/- 26.9 | 331737.4 +/- 75136.7 | 6 |
| hydrogenobacter_thermophilus | 83.5 +/- 4.7 | 188.2 +/- 29.1 | 443729.6 +/- 73050.4 | 6 |
| lactobacillus_delbrueckii | 83.1 +/- 12.1 | 200.2 +/- 49.8 | 414823.3 +/- 119660.0 | 6 |
| chondromyces_apiculatus | 2.8 +/- 1.7 | 204.0 +/- 204.5 | 13922.0 +/- 16219.5 | 6 |
| lactobacillus_crispatus | 79.2 +/- 4.5 | 204.2 +/- 45.5 | 387803.7 +/- 89167.0 | 8 |
| patulibacter_medicamentivorans | 46.4 +/- 2.6 | 208.5 +/- 29.1 | 222412.5 +/- 33448.0 | 7 |
| rhodanobacter_fulvus | 28.0 +/- 3.1 | 214.5 +/- 48.3 | 130616.9 +/- 32836.8 | 6 |
| sorangium_cellulosum | 12.5 +/- 3.2 | 216.3 +/- 69.1 | 57876.1 +/- 23761.7 | 6 |
| photobacterium_profundum | 6.3 +/- 2.1 | 220.1 +/- 98.8 | 28731.7 +/- 16154.5 | 8 |
| azorhizobium_caulinodans | 20.3 +/- 2.7 | 228.3 +/- 59.0 | 88866.9 +/- 25930.6 | 5 |
| bombiscardovia_coagulans | 69.8 +/- 4.2 | 238.5 +/- 67.4 | 292655.0 +/- 84576.7 | 5 |
| geobacillus_kaustophilus | 76.4 +/- 14.3 | 240.3 +/- 53.8 | 318022.4 +/- 92751.4 | 8 |
| leptospira_interrogans | 53.9 +/- 8.2 | 244.0 +/- 37.4 | 220774.2 +/- 47760.9 | 6 |
| methanococcoides_burtonii | 4.0 +/- 4.2 | 250.5 +/- 276.4 | 15865.3 +/- 24175.6 | 3 |
| chlamydia_trachomatis | 59.5 +/- 14.5 | 254.6 +/- 44.9 | 233725.9 +/- 70336.2 | 6 |
| salinibacter_ruber | 2.7 +/- 2.0 | 258.4 +/- 175.7 | 10331.7 +/- 10534.8 | 4 |
| pseudomonas_syringae | 60.8 +/- 19.4 | 260.4 +/- 74.7 | 233608.1 +/- 100210.4 | 7 |
| actinocatenispora_sera | 2.0 +/- 1.6 | 269.6 +/- 388.1 | 7359.3 +/- 12102.2 | 3 |
| kosmotoga_olearia | 92.7 +/- 3.6 | 270.2 +/- 52.4 | 342986.5 +/- 67839.3 | 7 |
| parafannyhessea_umbonata | 126.2 +/- 8.6 | 271.2 +/- 52.8 | 465346.6 +/- 95938.3 | 6 |
| limosilactobacillus_reuteri | 39.3 +/- 4.1 | 273.3 +/- 62.4 | 143896.5 +/- 36042.6 | 6 |
| geobacillus_stearothermophilus | 83.3 +/- 5.0 | 273.8 +/- 48.6 | 304196.8 +/- 57022.8 | 6 |
| acidimicrobium_ferrooxidans | 41.8 +/- 3.3 | 274.9 +/- 36.0 | 152160.5 +/- 23329.8 | 7 |
| micromonospora_viridifaciens | 91.3 +/- 9.1 | 275.9 +/- 58.1 | 330930.2 +/- 77102.6 | 7 |
| bryobacter_aggregatus | 33.9 +/- 8.2 | 281.6 +/- 47.7 | 120522.6 +/- 35473.1 | 7 |
| methanosarcina_acetivorans | 54.9 +/- 5.2 | 283.6 +/- 65.6 | 193557.9 +/- 48366.7 | 6 |
| corynebacterium_urealyticum | 86.4 +/- 11.4 | 284.4 +/- 42.4 | 303810.0 +/- 60449.0 | 6 |
| natranaerobius_thermophilus | 97.6 +/- 15.5 | 293.5 +/- 56.3 | 332651.6 +/- 82744.8 | 6 |
| salinispora_arenicola | 128.4 +/- 16.3 | 296.3 +/- 53.6 | 433359.9 +/- 95778.7 | 6 |
| deinococcus_deserti | 107.3 +/- 12.2 | 300.5 +/- 38.2 | 356975.5 +/- 60923.6 | 8 |
| microbacterium_chocolatum | 134.9 +/- 12.1 | 309.5 +/- 67.8 | 435813.6 +/- 103166.7 | 7 |
| faecalibaculum_rodentium | 3.4 +/- 2.0 | 317.9 +/- 506.9 | 10588.3 +/- 17968.0 | 5 |
| mesomycoplasma_flocculare | 24.6 +/- 3.6 | 320.4 +/- 53.5 | 76622.7 +/- 16945.1 | 6 |
| glaciecola_punicea | 1.8 +/- 0.9 | 320.7 +/- 274.6 | 5621.8 +/- 5516.4 | 4 |
| lysinibacillus_sphaericus | 146.6 +/- 13.1 | 329.1 +/- 75.7 | 445460.7 +/- 109951.5 | 6 |
| dietzia_alimentaria | 56.9 +/- 2.7 | 336.8 +/- 41.1 | 169066.1 +/- 22089.1 | 6 |
| bartonella_henselae | 77.9 +/- 15.6 | 342.3 +/- 40.0 | 227462.4 +/- 52773.2 | 6 |
| veillonella_seminalis | 132.2 +/- 12.8 | 343.7 +/- 81.9 | 384776.0 +/- 98934.2 | 6 |
| falsiroseomonas_stagni | 96.0 +/- 5.2 | 345.6 +/- 80.9 | 277768.5 +/- 66799.7 | 5 |
| chrysiogenes_arsenatis | 71.4 +/- 22.4 | 358.8 +/- 128.4 | 199034.4 +/- 94643.3 | 6 |
| symbiobacterium_thermophilum | 42.5 +/- 7.7 | 374.1 +/- 88.9 | 113530.7 +/- 33963.3 | 9 |
| virgibacillus_dokdonensis | 195.0 +/- 16.2 | 379.1 +/- 104.2 | 514294.1 +/- 147691.7 | 6 |
| pseudoalteromonas_atlantica | 7.7 +/- 1.3 | 397.0 +/- 452.1 | 19300.9 +/- 22231.5 | 5 |
| chitinophaga_pinensis | 68.2 +/- 6.1 | 401.4 +/- 72.5 | 169831.2 +/- 34182.6 | 7 |
| trichlorobacter_lovleyi | 53.9 +/- 12.6 | 405.7 +/- 123.8 | 132834.7 +/- 51125.0 | 7 |
| gemmobacter_aquatilis | 68.2 +/- 8.5 | 417.4 +/- 47.5 | 163283.2 +/- 27559.3 | 7 |
| prosthecobacter_debontii | 3.8 +/- 3.0 | 420.6 +/- 281.8 | 9105.7 +/- 9322.8 | 5 |
| pedobacter_hartonius | 38.1 +/- 4.4 | 421.1 +/- 64.6 | 90532.8 +/- 17424.0 | 6 |
| bdellovibrio_bacteriovorus | 121.4 +/- 17.4 | 421.5 +/- 58.6 | 287920.3 +/- 57445.8 | 8 |
| rubritepida_flocculans | 90.8 +/- 13.2 | 428.3 +/- 90.0 | 211998.0 +/- 54204.2 | 8 |
| megasphaera_paucivorans | 118.9 +/- 10.5 | 430.8 +/- 104.6 | 275956.1 +/- 71296.4 | 5 |
| pseudoroseomonas_rhizosphaerae | 115.4 +/- 9.8 | 438.9 +/- 110.2 | 262860.4 +/- 69662.9 | 7 |

| | | | | |
|---|---|---|---|---|
| mycoplasmoides_pneumoniae | 72.7 +/- 16.9 | 446.7 +/- 153.4 | 162753.6 +/- 67479.1 | 8 |
| heliomicrobium_modesticaldum | 59.0 +/- 3.3 | 447.8 +/- 76.5 | 131868.5 +/- 23725.6 | 5 |
| gluconacetobacter_diazotrophicus | 99.0 +/- 14.6 | 450.4 +/- 74.1 | 219785.1 +/- 48606.6 | 9 |
| dactylosporangium_aurantiacum | 10.9 +/- 1.8 | 454.1 +/- 116.4 | 23993.4 +/- 7329.2 | 6 |
| adlercreutzia_mucosicola | 170.0 +/- 20.6 | 455.0 +/- 120.7 | 373734.0 +/- 109039.4 | 7 |
| pseudobdellovibrio_exovorus | 163.6 +/- 8.0 | 456.7 +/- 99.2 | 358161.7 +/- 79735.1 | 7 |
| mycoplasma_mobile | 88.3 +/- 4.8 | 460.2 +/- 84.9 | 191899.3 +/- 36892.2 | 6 |
| geobacillus_thermodenitrificans | 119.8 +/- 18.9 | 463.7 +/- 130.2 | 258436.8 +/- 83227.0 | 7 |
| salinispira_pacifica | 72.4 +/- 10.9 | 467.1 +/- 86.0 | 154897.7 +/- 36853.3 | 8 |
| streptomonospora_alba | 4.8 +/- 2.0 | 468.7 +/- 172.4 | 10339.5 +/- 5710.0 | 3 |
| histophilus_somni | 182.4 +/- 14.2 | 484.6 +/- 106.9 | 376456.5 +/- 88119.1 | 6 |
| mycoplasma_mycoides | 108.7 +/- 29.9 | 488.7 +/- 53.4 | 222515.7 +/- 65867.5 | 6 |
| pyrinomonas_methylaliphatogenes | 80.1 +/- 14.3 | 490.2 +/- 68.7 | 163332.9 +/- 37038.3 | 7 |
| methylococcus_capsulatus | 204.2 +/- 38.6 | 491.8 +/- 117.7 | 415346.2 +/- 126637.7 | 8 |
| asanoa_hainanensis | 5.2 +/- 2.4 | 494.2 +/- 257.7 | 10510.7 +/- 7294.8 | 6 |
| marinobacter_nauticus | 151.4 +/- 16.6 | 496.7 +/- 123.6 | 304854.9 +/- 82926.0 | 7 |
| shewanella_frigidimarina | 96.5 +/- 7.1 | 499.1 +/- 42.8 | 193333.9 +/- 21894.7 | 7 |
| chlorobaculum_parvum | 10.2 +/- 2.8 | 500.3 +/- 129.2 | 20443.3 +/- 7674.4 | 5 |
| fibrobacter_intestinalis | 243.6 +/- 17.0 | 503.0 +/- 144.0 | 484438.2 +/- 142722.8 | 5 |
| nonlabens_ulvanivorans | 57.5 +/- 9.3 | 519.0 +/- 41.8 | 110707.2 +/- 20038.0 | 7 |
| persephonella_hydrogeniphila | 136.5 +/- 29.5 | 534.7 +/- 71.2 | 255216.4 +/- 64730.4 | 5 |
| streptococcus_thermophilus | 69.4 +/- 11.3 | 534.8 +/- 110.6 | 129740.1 +/- 34187.8 | 7 |
| pseudomonas_entomophila | 138.2 +/- 20.0 | 535.5 +/- 55.0 | 257989.2 +/- 45776.9 | 5 |
| mycoplasmopsis_arginini | 176.0 +/- 6.4 | 535.6 +/- 99.6 | 328531.4 +/- 62282.7 | 7 |
| escherichia_coli | 193.8 +/- 37.5 | 538.5 +/- 91.6 | 359863.8 +/- 92713.9 | 7 |
| mesonia_phycicola | 309.0 +/- 26.0 | 540.6 +/- 208.8 | 571522.6 +/- 225866.0 | 6 |
| shewanella_loihica | 223.5 +/- 8.8 | 546.3 +/- 79.5 | 409053.2 +/- 61700.9 | 6 |
| agromyces_subbeticus | 128.9 +/- 23.1 | 567.9 +/- 80.0 | 227016.2 +/- 51758.7 | 7 |
| polaromonas_naphthalenivorans | 118.2 +/- 22.8 | 568.6 +/- 55.5 | 207838.0 +/- 44949.5 | 6 |
| rickettsiella_massiliensis | 16.5 +/- 3.9 | 572.0 +/- 50.4 | 28822.4 +/- 7243.2 | 6 |
| thermotoga_neapolitana | 30.9 +/- 7.8 | 581.0 +/- 63.9 | 53222.5 +/- 14598.9 | 4 |
| petrotoga_mobilis | 200.3 +/- 16.8 | 587.1 +/- 74.5 | 341178.3 +/- 51892.5 | 7 |
| aliivibrio_salmonicida | 4.3 +/- 2.1 | 590.0 +/- 519.6 | 7312.0 +/- 7324.2 | 3 |
| oleiphilus_messinensis | 9.4 +/- 4.0 | 597.9 +/- 147.2 | 15804.8 +/- 7791.4 | 6 |
| paenibacillus_bovis | 167.9 +/- 14.3 | 608.7 +/- 124.6 | 275860.0 +/- 61194.7 | 7 |
| bacillus_subtilis | 269.7 +/- 41.8 | 614.4 +/- 108.9 | 438945.6 +/- 103389.5 | 8 |
| capnocytophaga_ochracea | 5.3 +/- 0.9 | 627.5 +/- 114.2 | 8431.0 +/- 2107.4 | 5 |
| bacillus_licheniformis | 116.4 +/- 11.2 | 631.1 +/- 131.1 | 184466.9 +/- 42260.1 | 6 |
| methylophaga_frappieri | 164.6 +/- 19.9 | 635.9 +/- 170.7 | 258788.6 +/- 76152.5 | 7 |
| erwinia_tasmaniensis | 128.0 +/- 12.3 | 646.7 +/- 66.0 | 197988.4 +/- 27773.5 | 5 |
| flavobacterium_hydatis | 102.2 +/- 11.4 | 650.6 +/- 96.9 | 157054.0 +/- 29207.4 | 6 |
| streptococcus_pneumoniae | 46.8 +/- 14.2 | 656.2 +/- 157.4 | 71342.3 +/- 27584.4 | 7 |
| anaeromyxobacter_dehalogenans | 147.9 +/- 8.8 | 666.5 +/- 96.0 | 221847.7 +/- 34588.3 | 8 |
| pseudothermotoga_lettingae | 164.3 +/- 11.3 | 667.8 +/- 147.8 | 245995.4 +/- 57033.1 | 7 |
| enterococcus_faecalis | 135.8 +/- 10.9 | 671.3 +/- 92.4 | 202220.1 +/- 32267.4 | 5 |
| bacillus_cereus | 205.2 +/- 28.8 | 675.8 +/- 159.5 | 303667.8 +/- 83396.8 | 6 |
| clostridium_botulinum | 229.2 +/- 35.2 | 680.8 +/- 113.1 | 336632.5 +/- 76190.2 | 6 |
| caldanaerobacter_subterraneus | 109.4 +/- 18.7 | 696.4 +/- 80.8 | 157090.0 +/- 32454.4 | 6 |
| methanocella_arvoryzae | 228.0 +/- 29.8 | 698.6 +/- 178.1 | 326388.5 +/- 93456.8 | 7 |
| neisseria_gonorrhoeae | 102.0 +/- 13.4 | 704.9 +/- 103.1 | 144638.0 +/- 28416.9 | 7 |
| xanthomonas_campestris | 82.9 +/- 7.0 | 716.8 +/- 66.9 | 115622.7 +/- 14597.3 | 4 |
| effusibacillus_pohliae | 171.9 +/- 18.3 | 719.1 +/- 144.4 | 238989.5 +/- 54346.8 | 6 |
| geotalea_daltonii | 127.9 +/- 13.9 | 742.2 +/- 134.8 | 172382.4 +/- 36485.3 | 6 |
| consensus_adk | 5.4 +/- 1.2 | 803.9 +/- 423.1 | 6704.2 +/- 3822.9 | 5 |
| desulfitobacterium_hafniense | 263.5 +/- 28.7 | 804.1 +/- 178.0 | 327630.0 +/- 80797.7 | 6 |
| helicobacter_acinonychis | 8.1 +/- 1.6 | 850.7 +/- 279.4 | 9509.7 +/- 3668.3 | 7 |

| | | | | |
|---|---|---|---|---|
| **oenococcus_oeni** | **15.2 +/- 4.3** | **858.1 +/- 273.2** | **17661.3 +/- 7523.8** | **9** |
| **legionella_erythra** | **275.3 +/- 22.2** | **859.5 +/- 189.1** | **320350.4 +/- 75028.0** | **6** |
| **salipiger_mucosus** | **62.2 +/- 0.6** | **860.5 +/- 53.0** | **72243.0 +/- 4497.4** | **5** |
| **bordetella_petrii** | **169.6 +/- 18.9** | **903.5 +/- 157.0** | **187720.5 +/- 38720.3** | **7** |
| **acaryochloris_marina** | **5.5 +/- 2.2** | **916.4 +/- 1178.5** | **6018.6 +/- 8118.0** | **4** |
| **actinobacillus_pleuropneumoniae** | **360.6 +/- 36.7** | **925.0 +/- 270.6** | **389886.4 +/- 120786.4** | **8** |
| **verminephrobacter_aporrectodeae** | **1.4 +/- 0.3** | **932.1 +/- 633.5** | **1541.6 +/- 1112.9** | **4** |
| **acetomicrobium_flavidum** | **491.1 +/- 34.5** | **960.1 +/- 195.3** | **511538.3 +/- 110106.9** | **8** |
| **allofrancisella_guangzhouensis** | **273.9 +/- 41.7** | **970.5 +/- 209.2** | **282256.4 +/- 74480.7** | **7** |
| **helicobacter_rodentium** | **76.7 +/- 7.8** | **982.5 +/- 194.1** | **78064.5 +/- 17351.8** | **6** |
| **thermotoga_maritima** | **125.1 +/- 12.6** | **986.3 +/- 88.7** | **126818.4 +/- 17150.5** | **7** |
| **sphingopyxis_alaskensis** | **144.0 +/- 13.3** | **1014.9 +/- 244.8** | **141851.5 +/- 36636.3** | **8** |
| **legionella_pneumophila** | **318.1 +/- 59.5** | **1020.7 +/- 221.4** | **311698.5 +/- 89292.0** | **6** |
| **alkaliphilus_oremlandii** | **223.4 +/- 39.2** | **1025.2 +/- 110.0** | **217853.8 +/- 44785.6** | **7** |
| **thermosipho_melanesiensis** | **343.3 +/- 31.5** | **1027.7 +/- 282.4** | **334026.4 +/- 96749.3** | **7** |
| **mesomycoplasma_hyorhinis** | **250.2 +/- 29.7** | **1049.6 +/- 153.0** | **238412.6 +/- 44790.9** | **7** |
| **methylocella_silvestris** | **123.1 +/- 20.4** | **1066.6 +/- 97.5** | **115377.1 +/- 21818.7** | **9** |
| **sphaerochaeta_pleomorpha** | **284.4 +/- 11.5** | **1075.6 +/- 294.5** | **264402.0 +/- 73185.2** | **6** |
| **brachyspira_pilosicoli** | **215.7 +/- 19.4** | **1109.1 +/- 191.4** | **194518.7 +/- 37880.7** | **7** |
| **vibrio_cholerae** | **420.8 +/- 69.3** | **1137.7 +/- 298.1** | **369870.3 +/- 114457.3** | **8** |
| **kushneria_avicenniae** | **9.4 +/- 3.0** | **1172.2 +/- 1961.5** | **8056.3 +/- 13722.9** | **6** |
| **lactococcus_lactis** | **358.3 +/- 28.8** | **1175.2 +/- 303.7** | **304857.4 +/- 82495.7** | **7** |
| **vibrio_parahaemolyticus** | **319.8 +/- 73.0** | **1188.8 +/- 292.6** | **269006.8 +/- 90308.4** | **6** |
| **magnetospirillum_magneticum** | **502.0 +/- 43.8** | **1198.7 +/- 187.7** | **418749.4 +/- 75066.9** | **8** |
| **thermosipho_africanus** | **478.5 +/- 51.7** | **1262.5 +/- 257.0** | **379002.3 +/- 87352.2** | **10** |
| **alkalilimnicola_ehrlichii** | **488.0 +/- 27.5** | **1265.7 +/- 273.0** | **385563.0 +/- 85939.3** | **6** |
| **fervidobacterium_islandicum** | **493.2 +/- 54.6** | **1348.9 +/- 336.9** | **365631.9 +/- 99898.9** | **7** |
| **sporolituus_thermophilus** | **434.2 +/- 62.3** | **1418.0 +/- 248.6** | **306237.8 +/- 69406.3** | **7** |
| **lachnospira_multipara** | **3.8 +/- 2.8** | **1486.6 +/- 933.4** | **2583.3 +/- 2488.0** | **4** |
| **halobacterium_salinarum** | **5.6 +/- 4.6** | **1504.9 +/- 1562.5** | **3733.2 +/- 4925.2** | **5** |
| **archaeoglobus_fulgidus** | **525.7 +/- 56.8** | **1529.6 +/- 228.4** | **343690.9 +/- 63337.9** | **8** |
| **shewanella_halifaxensis** | **28.9 +/- 5.5** | **1554.5 +/- 211.5** | **18578.5 +/- 4322.9** | **7** |
| **psychromonas_ingrahamii** | **7.8 +/- 1.4** | **1572.2 +/- 979.3** | **4971.3 +/- 3214.4** | **5** |
| **treponema_putidum** | **285.7 +/- 53.5** | **1573.8 +/- 400.9** | **181535.4 +/- 57390.6** | **6** |
| **fervidobacterium_nodosum** | **522.5 +/- 27.2** | **1600.1 +/- 329.0** | **326550.4 +/- 69266.5** | **7** |
| **halorhodospira_halophila** | **472.0 +/- 38.3** | **1600.6 +/- 265.3** | **294904.6 +/- 54424.6** | **8** |
| **moorella_thermoacetica** | **448.5 +/- 46.0** | **1669.8 +/- 264.4** | **268617.5 +/- 50677.9** | **7** |
| **desulfatibacillum_alkenivorans** | **479.6 +/- 62.4** | **1702.3 +/- 334.6** | **281730.3 +/- 66409.8** | **8** |
| **spirulina_subsalsa** | **4.2 +/- 1.3** | **1712.4 +/- 2482.5** | **2477.6 +/- 3671.9** | **5** |
| **dissulfuribacter_thermophilus** | **802.8 +/- 45.5** | **1724.1 +/- 279.0** | **465636.5 +/- 79848.1** | **9** |
| **elusimicrobium_minutum** | **138.4 +/- 14.4** | **1725.2 +/- 122.4** | **80200.0 +/- 10124.4** | **7** |
| **thermodesulfatator_atlanticus** | **340.6 +/- 36.5** | **1758.3 +/- 246.5** | **193731.1 +/- 34197.6** | **8** |
| **thermodesulfobacterium_commune** | **294.6 +/- 19.1** | **1781.8 +/- 219.2** | **165322.8 +/- 22989.3** | **6** |
| **pyrococcus_furiosus** | **57.3 +/- 5.8** | **1804.6 +/- 460.9** | **31759.3 +/- 8724.3** | **5** |
| **thermodesulfovibrio_yellowstonii** | **479.1 +/- 68.3** | **1880.4 +/- 245.9** | **254806.3 +/- 49289.2** | **8** |
| **prochlorococcus_marinus** | **3.6 +/- 0.8** | **1975.5 +/- 1300.9** | **1827.1 +/- 1272.7** | **3** |
| **chlorobaculum_tepidum** | **183.4 +/- 19.6*** | **2421.1 +/- 99.4*** | **75756.5 +/- 8684.5*** | **7** |
| **desulfovibrio_desulfuricans** | **733.5 +/- 40.9*** | **3317.7 +/- 768.1*** | **221081.4 +/- 52645.4*** | **6** |
| **thermococcus_kodakarensis** | **53.1 +/- 5.2*** | **3333.7 +/- 1106.5*** | **15916.1 +/- 5507.6*** | **5** |
| **thermococcus_gammatolerans** | **30.8 +/- 4.1*** | **3558.8 +/- 762.6*** | **8663.1 +/- 2181.6*** | **6** |
| **mycoplasmopsis_canis** | **405.5 +/- 39.7*** | **4046.5 +/- 252.2*** | **100208.7 +/- 11621.9*** | **8** |

***$K_M$ outside assay bounds (>2000µM or <31.25uM)**

**Table S3. Catalytic Parameters for ADK Mutants**

| Variant | k_cat (s^-1) | K_M (uM) | k_cat/K_M (s^-1 M^-1) | Replicates |
|---|---|---|---|---|
| A_bayl_WT | 67.8 +/- 7.3 | 126.3 +/- 15.9 | 536903.0 +/- 89256.1 | 8 |
| A_ehrl_WT | 474.0 +/- 87.8 | 1533.8 +/- 255.1 | 309061.7 +/- 76909.4 | 11 |
| A_fulg_WT | 572.8 +/- 88.2 | 1421.3 +/- 200.6 | 403057.4 +/- 84182.9 | 7 |
| A_medi_WT | 39.8 +/- 8.3 | 171.4 +/- 40.5 | 231889.1 +/- 73282.3 | 8 |
| A_pleu_WT | 422.1 +/- 53.4 | 897.8 +/- 180.8 | 470145.3 +/- 111805.4 | 8 |
| B_petr_WT | 118.5 +/- 11.0 | 660.6 +/- 78.1 | 179312.7 +/- 26906.7 | 8 |
| C_japo_WT | 25.7 +/- 3.9 | 109.1 +/- 6.7 | 235911.9 +/- 38675.9 | 8 |
| C_psyc_WT | 2.1 +/- 1.2 | 501.7 +/- 476.5 | 4175.5 +/- 4620.4 | 5 |
| C_tepi_WT | 212.3 +/- 27.8* | 2355.0 +/- 118.7* | 90149.5 +/- 12649.3* | 8 |
| D_dese_WT | 99.4 +/- 11.5 | 264.1 +/- 16.6 | 376338.0 +/- 49529.3 | 9 |
| H_somn_WT | 212.0 +/- 27.0 | 482.8 +/- 49.0 | 439155.4 +/- 71514.0 | 7 |
| L_pneu_WT | 336.6 +/- 48.8 | 1095.0 +/- 170.3 | 307410.8 +/- 65357.5 | 8 |
| M_arvo_WT | 172.7 +/- 17.8 | 457.1 +/- 67.3 | 377846.5 +/- 67892.6 | 8 |
| N_gono_WT | 89.6 +/- 14.7 | 618.3 +/- 34.9 | 144894.4 +/- 25115.4 | 8 |
| P_atla_WT | 12.1 +/- 2.8 | 161.3 +/- 48.1 | 75174.9 +/- 28330.4 | 7 |
| P_ingr_WT | 5.0 +/- 2.1 | 723.0 +/- 468.7 | 6859.3 +/- 5320.0 | 8 |
| P_naph_WT | 108.1 +/- 28.4 | 595.5 +/- 74.3 | 181579.6 +/- 52806.6 | 8 |
| P_prof_WT | 6.3 +/- 1.7 | 294.8 +/- 154.2 | 21293.3 +/- 12516.1 | 7 |
| S_frig_WT | 108.7 +/- 17.6 | 426.1 +/- 34.7 | 255150.1 +/- 46188.8 | 8 |
| S_loih_WT | 213.3 +/- 31.5 | 578.2 +/- 51.9 | 368911.0 +/- 63771.5 | 10 |
| Sym_ther_WT | 32.0 +/- 2.0 | 321.8 +/- 35.3 | 99526.4 +/- 12537.5 | 7 |
| T_yell_WT | 329.5 +/- 18.6 | 1580.5 +/- 103.2 | 208485.6 +/- 17991.0 | 9 |
| V_chol_WT | 325.8 +/- 81.6 | 1135.7 +/- 214.3 | 286848.0 +/- 89953.8 | 9 |
| a_bayl_P127A | 18.1 +/- 0.6 | 68.2 +/- 4.2 | 265179.0 +/- 18730.1 | 6 |
| a_ehrl_A18E | 534.1 +/- 39.0 | 1626.3 +/- 269.5 | 328441.6 +/- 59467.1 | 8 |
| a_ehrl_A18K | 482.6 +/- 43.5 | 1439.9 +/- 260.6 | 335125.3 +/- 67752.2 | 8 |
| a_ehrl_A18Q | 196.2 +/- 25.8 | 981.3 +/- 125.8 | 199962.9 +/- 36689.7 | 8 |
| a_ehrl_E22A | 391.3 +/- 36.6 | 1310.3 +/- 144.0 | 298647.5 +/- 43101.5 | 8 |
| a_ehrl_E22K | 485.1 +/- 39.1 | 1347.2 +/- 283.2 | 360070.4 +/- 81064.3 | 7 |
| a_ehrl_P127A | 471.4 +/- 72.2 | 1420.1 +/- 228.2 | 331916.2 +/- 73662.7 | 8 |
| a_fulg_N132A | 221.4 +/- 10.6 | 1176.6 +/- 155.9 | 188170.1 +/- 26513.9 | 8 |
| a_fulg_N132P | 258.8 +/- 13.9 | 1035.3 +/- 141.5 | 249979.4 +/- 36708.9 | 8 |
| a_medi_P127A | 22.6 +/- 8.6 | 113.1 +/- 24.0 | 199982.4 +/- 87164.1 | 7 |
| a_pleu_P128A | 347.3 +/- 36.9 | 828.2 +/- 155.8 | 419305.9 +/- 90602.4 | 7 |
| b_petr_p127A | 179.2 +/- 28.6 | 751.6 +/- 93.5 | 238472.4 +/- 48246.7 | 9 |
| b_subt_ADK_ec_lid | 65.1 +/- 5.4 | 340.9 +/- 47.0 | 190862.5 +/- 30627.7 | 8 |
| b_subt_ADK_gs_lid | 147.4 +/- 8.3 | 574.9 +/- 152.5 | 256350.6 +/- 69511.9 | 7 |
| b_subt_V132A | 183.9 +/- 8.5 | 618.2 +/- 79.0 | 297473.1 +/- 40412.7 | 6 |
| b_subt_WT | 169.0 +/- 6.6 | 567.3 +/- 80.1 | 297976.2 +/- 43663.8 | 9 |
| bsADK_S131P | 308.3 +/- 32.5 | 830.0 +/- 140.6 | 371497.8 +/- 74086.5 | 8 |
| bsADK_V132P | 208.6 +/- 33.4 | 609.3 +/- 46.0 | 342342.8 +/- 60604.4 | 9 |
| c_japo_A128P | 61.0 +/- 7.4 | 171.5 +/- 24.8 | 355382.9 +/- 67106.0 | 8 |
| c_japo_E127P | 53.0 +/- 7.6 | 197.4 +/- 20.9 | 268462.2 +/- 47929.3 | 9 |
| c_psyc_G128P | 3.6 +/- 1.7 | 670.8 +/- 466.4 | 5368.9 +/- 4496.3 | 5 |
| c_tepi_A128P | 101.2 +/- 6.8* | 2554.1 +/- 330.1* | 39622.2 +/- 5767.3* | 8 |
| d_dese_ADK_bs_lid | 27.6 +/- 4.8 | 97.5 +/- 23.0 | 283420.0 +/- 82856.6 | 8 |
| d_dese_ADK_ec_lid | 5.7 +/- 0.9* | 22.3 +/- 11.9* | 254790.1 +/- 141751.1* | 8 |
| d_dese_ADK_gs_lid | 12.4 +/- 1.9 | 41.4 +/- 10.6 | 298462.4 +/- 89660.2 | 7 |
| d_dese_ADK_vc_lid | 30.6 +/- 1.9 | 157.9 +/- 13.1 | 193612.3 +/- 20163.6 | 8 |
| e_coli_ADK_bs_lid | 19.8 +/- 2.5 | 231.4 +/- 46.4 | 85570.8 +/- 20276.0 | 6 |
| e_coli_ADK_gs_lid | 10.0 +/- 1.1 | 87.9 +/- 34.8 | 113599.4 +/- 46626.8 | 6 |
| e_coli_ADK_vc_lid | 233.1 +/- 23.7 | 762.2 +/- 346.5 | 305858.3 +/- 142496.4 | 9 |
| e_coli_WT | 179.7 +/- 15.2 | 520.6 +/- 62.6 | 345273.0 +/- 50781.2 | 8 |
| ecADK_CCCC | 133.3 +/- 13.1 | 390.5 +/- 63.6 | 341298.7 +/- 64956.0 | 9 |

| | | | | |
|---|---|---|---|---|
| ecADK_CCCT | 117.7 +/- 17.0 | 342.5 +/- 44.9 | 343775.1 +/- 66991.8 | 9 |
| ecADK_CCDC | 96.9 +/- 4.6 | 235.2 +/- 22.8 | 411722.7 +/- 44507.0 | 8 |
| ecADK_CCDT | 45.1 +/- 9.7* | 3419.7 +/- 925.2* | 13196.5 +/- 4552.7* | 7 |
| ecADK_CSCC | 195.8 +/- 17.9 | 527.9 +/- 59.5 | 370993.9 +/- 53798.0 | 8 |
| ecADK_CSCT | 72.4 +/- 13.6 | 1124.7 +/- 74.5 | 64406.6 +/- 12841.9 | 8 |
| ecADK_CSDC | 55.6 +/- 9.6 | 1820.2 +/- 224.9 | 30540.9 +/- 6506.9 | 8 |
| ecADK_CSDT | 24.7 +/- 5.6* | 2561.9 +/- 596.6* | 9658.8 +/- 3137.7* | 6 |
| ecADK_HCCC | 126.7 +/- 23.0 | 401.5 +/- 41.3 | 315481.4 +/- 65865.1 | 8 |
| ecADK_HCCT | 136.0 +/- 22.0 | 908.8 +/- 112.3 | 149627.2 +/- 30464.6 | 8 |
| ecADK_HCDC | 88.0 +/- 33.6 | 1315.3 +/- 330.7 | 66920.9 +/- 30567.7 | 8 |
| ecADK_HCDT | 84.5 +/- 8.8* | 5545.7 +/- 1304.0* | 15246.0 +/- 3921.9* | 7 |
| ecADK_HSCC | 107.6 +/- 21.3* | 3793.2 +/- 263.6* | 28360.3 +/- 5959.4* | 9 |
| ecADK_HSCT | 76.7 +/- 7.8 | 1567.5 +/- 237.5 | 48961.3 +/- 8922.8 | 8 |
| ecADK_HSDC | 208.6 +/- 10.0 | 1154.7 +/- 30.4 | 180647.4 +/- 9894.1 | 7 |
| g_stea_A22E | 49.9 +/- 4.0 | 192.0 +/- 32.1 | 259823.0 +/- 48067.2 | 7 |
| g_stea_A22K | 72.0 +/- 11.2 | 244.7 +/- 50.6 | 294301.5 +/- 76131.9 | 8 |
| g_stea_A22Q | 43.2 +/- 4.0 | 199.5 +/- 15.0 | 216485.5 +/- 25905.8 | 6 |
| g_stea_ADK_bs_lid | 79.5 +/- 7.3 | 236.1 +/- 18.3 | 336842.3 +/- 40625.3 | 8 |
| g_stea_ADK_ec_lid | 101.7 +/- 7.7 | 309.9 +/- 23.5 | 328198.1 +/- 35182.3 | 7 |
| g_stea_ADK_vc_lid | 103.1 +/- 15.5 | 1291.9 +/- 64.6 | 79776.3 +/- 12637.1 | 9 |
| g_stea_E18A | 75.7 +/- 6.4 | 240.3 +/- 26.6 | 315187.8 +/- 43974.0 | 7 |
| g_stea_E18K | 106.1 +/- 11.1 | 394.7 +/- 58.2 | 268906.7 +/- 48579.1 | 8 |
| g_stea_E18Q | 16.0 +/- 2.1 | 187.3 +/- 41.3 | 85215.4 +/- 21984.5 | 8 |
| g_stea_N132P | 94.0 +/- 8.4 | 326.2 +/- 63.6 | 288048.3 +/- 61711.9 | 7 |
| g_stea_WT | 78.9 +/- 10.4 | 212.7 +/- 28.0 | 370994.7 +/- 69299.1 | 9 |
| gsADK_CCCC | 74.6 +/- 11.0 | 220.5 +/- 17.8 | 338436.1 +/- 56978.7 | 10 |
| gsADK_CCCT | 31.2 +/- 2.5 | 218.2 +/- 24.6 | 143170.5 +/- 19680.1 | 8 |
| gsADK_CCDC | 34.8 +/- 3.9 | 132.8 +/- 18.4 | 262475.6 +/- 46721.1 | 8 |
| gsADK_CSCC | 73.2 +/- 13.8 | 322.0 +/- 56.6 | 227445.9 +/- 58660.8 | 9 |
| gsADK_CSCT | 5.4 +/- 1.6 | 897.7 +/- 390.2 | 5994.8 +/- 3186.4 | 4 |
| gsADK_CSDC | 5.2 +/- 1.7 | 909.9 +/- 342.3 | 5719.4 +/- 2853.1 | 6 |
| gsADK_CSDT | 2.9 +/- 1.8 | 396.4 +/- 257.2 | 7283.2 +/- 6631.2 | 6 |
| gsADK_HCCC | 39.8 +/- 9.7* | 3832.2 +/- 746.2* | 10384.0 +/- 3248.9* | 9 |
| gsADK_HCDC | 6.3 +/- 2.5 | 1421.7 +/- 852.1 | 4454.9 +/- 3207.1 | 5 |
| gsADK_HCDT | 5.7 +/- 1.3 | 1054.0 +/- 842.2 | 5439.5 +/- 4516.5 | 4 |
| gsADK_HSCC | 9.0 +/- 2.9 | 1624.6 +/- 925.7 | 5542.5 +/- 3627.3 | 7 |
| gsADK_HSCT | 2.8 +/- 0.2 | 310.7 +/- 56.1 | 9021.0 +/- 1770.4 | 3 |
| gsADK_HSDC | 22.8 +/- 0.7 | 595.9 +/- 70.6 | 38201.0 +/- 4691.0 | 3 |
| gsADK_HSDT | 3.5 +/- 1.4 | 907.6 +/- 683.0 | 3836.3 +/- 3256.2 | 5 |
| h_somn_P127A | 131.3 +/- 27.3 | 265.9 +/- 43.8 | 493806.6 +/- 130887.8 | 9 |
| hbond_consensus | 255.8 +/- 27.2 | 1174.1 +/- 73.7 | 217832.7 +/- 26893.1 | 10 |
| jm_ADK | 31.1 +/- 4.1 | 160.9 +/- 18.8 | 192936.2 +/- 34110.4 | 7 |
| l_pneu_P128A | 128.5 +/- 13.8 | 804.8 +/- 145.8 | 159606.8 +/- 33595.2 | 8 |
| m_arvo_E22A | 19.3 +/- 3.9 | 220.9 +/- 40.0 | 87559.6 +/- 23831.1 | 8 |
| m_arvo_E22K | 107.5 +/- 20.1 | 306.2 +/- 53.5 | 351235.2 +/- 89850.5 | 9 |
| m_arvo_E22Q | 131.7 +/- 15.2 | 354.9 +/- 44.2 | 371170.7 +/- 62991.9 | 7 |
| m_arvo_K18A | 157.8 +/- 12.6 | 651.9 +/- 63.4 | 242046.0 +/- 30482.5 | 8 |
| m_arvo_K18E | 215.5 +/- 30.4 | 804.6 +/- 85.1 | 267814.5 +/- 47235.0 | 10 |
| m_arvo_K18Q | 149.2 +/- 12.1 | 612.4 +/- 44.3 | 243693.8 +/- 26477.4 | 7 |
| m_arvo_S132A | 77.9 +/- 7.0 | 349.4 +/- 56.4 | 222988.7 +/- 41213.0 | 9 |
| m_arvo_S132P | 204.4 +/- 14.0 | 620.9 +/- 96.4 | 329232.6 +/- 55867.8 | 8 |
| m_naut_E127P | 191.4 +/- 24.2 | 490.5 +/- 49.6 | 390168.8 +/- 63242.8 | 8 |
| m_naut_G128P | 133.3 +/- 12.4 | 466.2 +/- 48.4 | 285822.7 +/- 39887.0 | 8 |
| n_gono_A128P | 103.1 +/- 12.9 | 664.6 +/- 42.3 | 155112.4 +/- 21818.8 | 8 |
| o_acum_ADK_ec_lid | 3.1 +/- 0.8 | 1030.1 +/- 767.1 | 3019.7 +/- 2387.3 | 3 |
| o_acum_ADK_gs_lid | 1.5 +/- 0.8 | 260.5 +/- 460.1 | 5822.6 +/- 10786.8 | 6 |

| | | | | |
|---|---|---|---|---|
| o_acum_ADK_vc_lid | 3.1 +/- 2.3 | 172.2 +/- 114.5 | 18222.6 +/- 18081.5 | 4 |
| p_atla_G22A | 12.6 +/- 2.9 | 296.9 +/- 120.4 | 42303.6 +/- 19763.9 | 8 |
| p_atla_G22E | 11.3 +/- 2.2 | 472.8 +/- 208.2 | 23981.0 +/- 11513.6 | 8 |
| p_atla_G22K | 17.9 +/- 3.1 | 177.0 +/- 33.9 | 100961.9 +/- 25958.8 | 7 |
| p_atla_G22Q | 9.4 +/- 3.5 | 312.8 +/- 126.2 | 30129.3 +/- 16591.3 | 9 |
| p_atla_P127A | 4.3 +/- 1.4 | 404.4 +/- 353.0 | 10660.5 +/- 9941.3 | 4 |
| p_atla_Q18A | 9.3 +/- 1.9 | 332.4 +/- 95.6 | 28128.5 +/- 9991.8 | 7 |
| p_atla_Q18E | 13.3 +/- 4.0 | 608.3 +/- 430.7 | 21922.4 +/- 16829.7 | 7 |
| p_atla_Q18K | 10.8 +/- 2.2 | 449.9 +/- 190.8 | 24067.6 +/- 11292.9 | 7 |
| p_naph_A128P | 84.0 +/- 17.7 | 597.8 +/- 82.8 | 140430.2 +/- 35466.8 | 9 |
| p_prof_A22E | 8.7 +/- 2.1 | 439.9 +/- 241.0 | 19694.4 +/- 11795.4 | 8 |
| p_prof_A22Q | 10.1 +/- 1.7 | 258.2 +/- 43.0 | 38968.2 +/- 9337.0 | 8 |
| p_prof_Q18A | 4.2 +/- 1.6 | 529.5 +/- 300.6 | 7916.2 +/- 5383.6 | 7 |
| p_prof_Q18K | 9.0 +/- 4.7 | 501.2 +/- 424.5 | 17885.7 +/- 17797.7 | 6 |
| s_frig_G128P | 189.8 +/- 20.3 | 720.5 +/- 68.4 | 263356.2 +/- 37613.5 | 9 |
| s_loih_P127A | 134.6 +/- 16.3 | 597.0 +/- 31.7 | 225385.6 +/- 29880.1 | 9 |
| sym_therm_G22A | 27.7 +/- 2.9 | 447.7 +/- 159.9 | 61966.0 +/- 23088.2 | 8 |
| sym_therm_G22E | 27.2 +/- 2.7 | 315.3 +/- 59.6 | 86372.9 +/- 18407.1 | 7 |
| sym_therm_G22K | 40.0 +/- 5.6 | 372.2 +/- 47.0 | 107518.2 +/- 20308.8 | 8 |
| sym_therm_G22Q | 39.0 +/- 8.9 | 345.3 +/- 94.3 | 112871.8 +/- 40197.4 | 9 |
| sym_therm_S132A | 30.2 +/- 5.0 | 344.8 +/- 63.2 | 87633.9 +/- 21622.9 | 9 |
| sym_therm_S132P | 31.0 +/- 3.2 | 411.3 +/- 79.3 | 75428.3 +/- 16542.0 | 8 |
| sym_therm_V18A | 43.8 +/- 3.5 | 341.9 +/- 59.1 | 128160.1 +/- 24434.7 | 7 |
| sym_therm_V18E | 31.7 +/- 2.5 | 378.5 +/- 80.3 | 83633.9 +/- 18899.0 | 7 |
| sym_therm_V18K | 92.8 +/- 8.2 | 484.9 +/- 43.6 | 191338.0 +/- 24095.5 | 8 |
| sym_therm_V18Q | 46.5 +/- 6.0 | 291.5 +/- 31.0 | 159358.2 +/- 26592.2 | 7 |
| t_yell_E22A | 219.8 +/- 12.6 | 1580.5 +/- 77.0 | 139088.7 +/- 10467.3 | 6 |
| t_yell_E22K | 1094.9 +/- 126.7* | 2799.0 +/- 358.2* | 391170.0 +/- 67496.8* | 9 |
| t_yell_E22Q | 888.3 +/- 123.4* | 2350.0 +/- 265.6* | 377990.1 +/- 67712.7* | 9 |
| t_yell_K18A | 262.5 +/- 36.6* | 2751.6 +/- 255.3* | 95400.3 +/- 15975.3* | 8 |
| t_yell_K18E | 50.5 +/- 11.5* | 2501.6 +/- 1662.4* | 20202.0 +/- 14196.7* | 5 |
| t_yell_K18Q | 332.7 +/- 46.8* | 2914.8 +/- 264.1* | 114138.5 +/- 19096.5* | 9 |
| t_yell_S132A | 468.9 +/- 37.0 | 1820.0 +/- 124.0 | 257623.9 +/- 26858.8 | 8 |
| t_yell_S132P | 596.0 +/- 78.9 | 1924.9 +/- 299.1 | 309605.9 +/- 63200.3 | 8 |
| v_chol_ADK_bs_lid | 51.8 +/- 9.6 | 206.7 +/- 39.3 | 250670.6 +/- 66681.7 | 9 |
| v_chol_ADK_ec_lid | 105.6 +/- 9.9 | 513.1 +/- 216.3 | 205797.1 +/- 88884.8 | 8 |
| v_chol_ADK_gs_lid | 10.3 +/- 1.0 | 109.1 +/- 60.8 | 94182.4 +/- 53361.7 | 7 |
| v_chol_E22A | 178.9 +/- 33.5 | 705.9 +/- 128.1 | 253437.0 +/- 66107.7 | 8 |
| v_chol_E22K | 221.2 +/- 26.9 | 737.4 +/- 106.4 | 299936.5 +/- 56630.1 | 8 |
| v_chol_E22Q | 163.8 +/- 41.4 | 733.1 +/- 177.1 | 223451.9 +/- 78108.8 | 7 |
| v_chol_Q18A | 466.3 +/- 94.8 | 1393.6 +/- 241.6 | 334588.0 +/- 89387.9 | 7 |
| v_chol_Q18E | 311.2 +/- 43.1 | 1658.0 +/- 100.1 | 187670.5 +/- 28357.4 | 8 |
| v_chol_Q18K | 329.2 +/- 31.9 | 1092.6 +/- 78.8 | 301334.3 +/- 36431.6 | 7 |
| zinc_consensus | 9.0 +/- 1.2 | 277.7 +/- 91.7 | 32499.5 +/- 11599.6 | 8 |

***$K_M$ outside assay bounds (>2000µM or <31.25uM)**

### Supplementary Figures

**A**

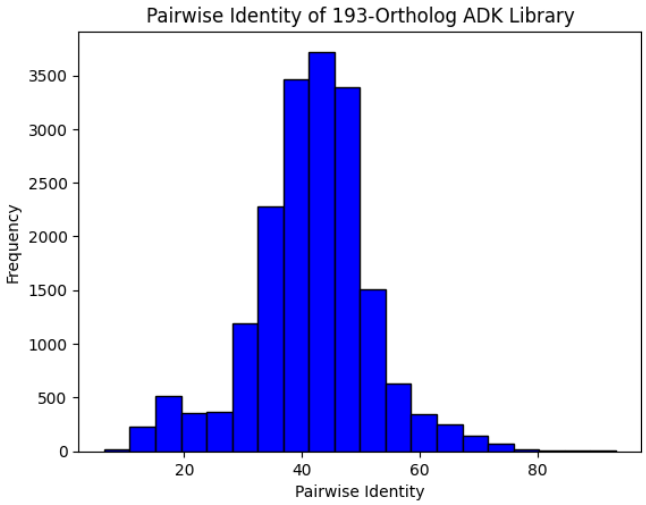


**B**

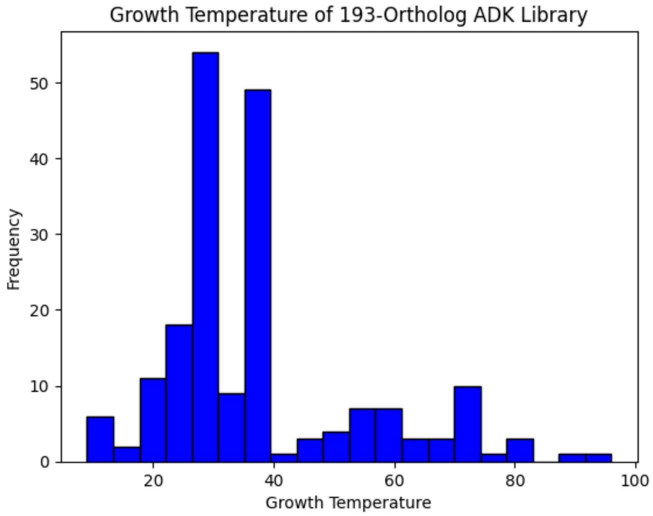


**Fig S1. Distribution of sequence identity and growth temperature of naturally-occurring ADK variants.** (**A**) Histogram of pairwise sequence identities obtained from a multiple sequence alignment (see Methods). (**B**) Histogram of median optimal growth temperatures for the organism associated with each ADK ortholog.

A

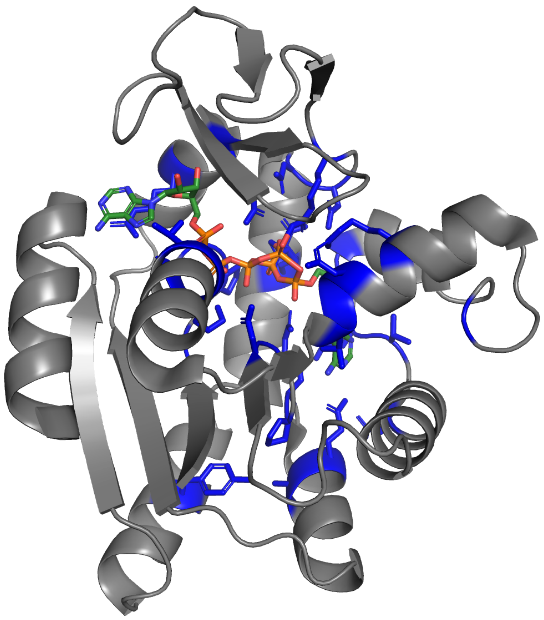

B

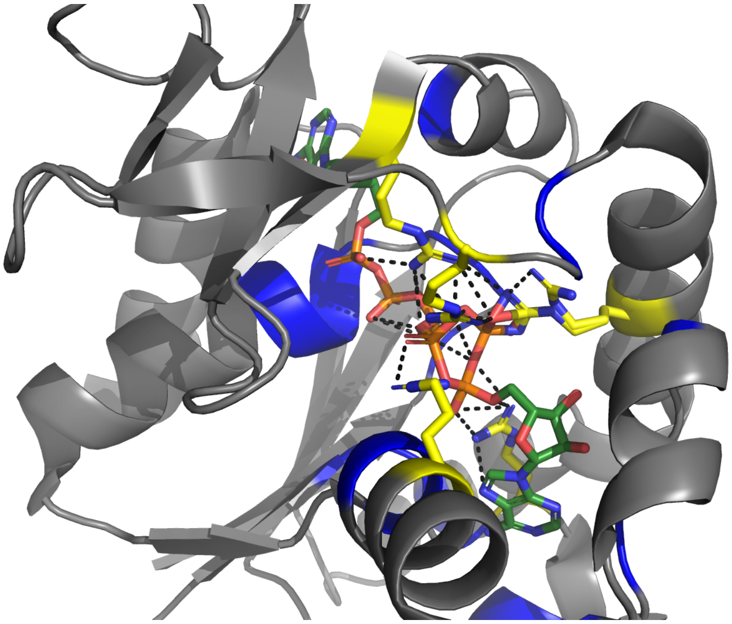

**Fig S2. ADK active site residues are conserved across naturally occurring ADKs assayed herein.** (**A**) Residues conserved in >90% of aligned ADK sequences from the ortholog library assayed herein are colored in blue with sidechains shown on ecADK (PDB: 1AKE) with substrate analog AP5 bound (green). (**B**) Structural zoom-in showing key catalytic arginines (yellow sticks).

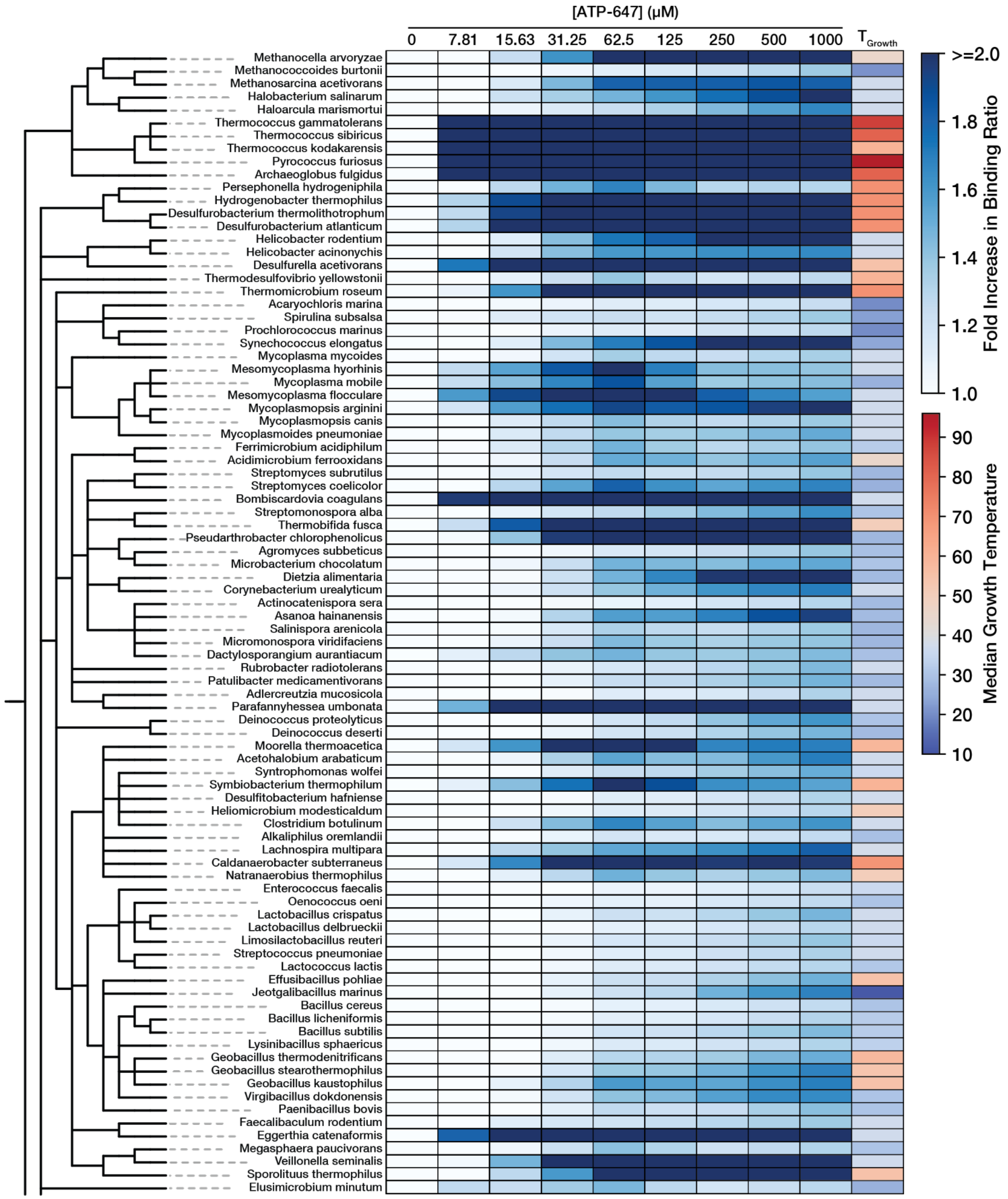


**Fig S3. Fluorescent ATP analog binding on-chip with ADK orthologs.** Fold increase in the ratio of ATP-647:ADK-eGFP summed RFU (CY5:eGFP) over buffer alone (0M fluorescent probe) as a heatmap organized by a taxonomical tree. Optimal growth temperature is also plotted.

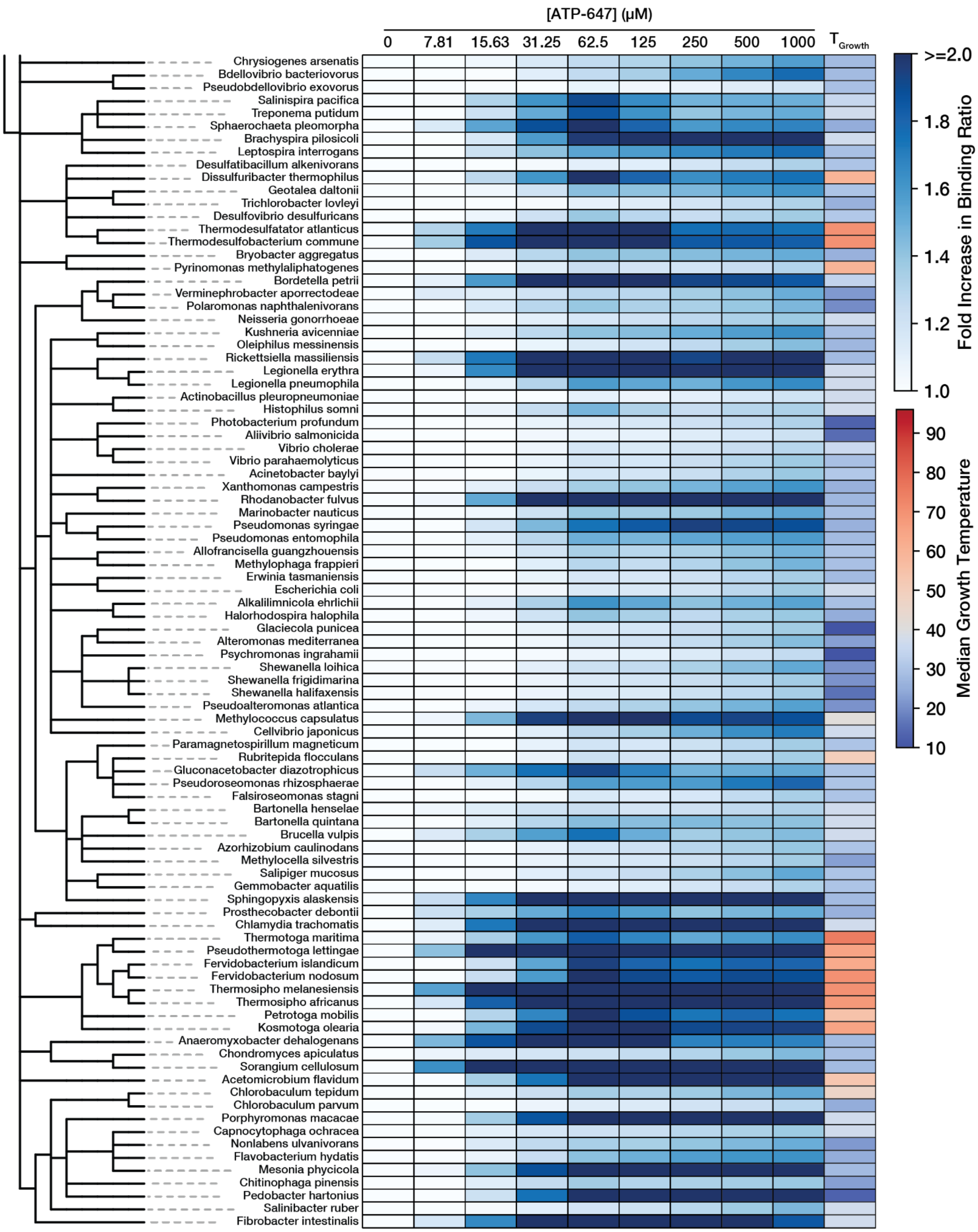


**Fig S3. (cont.) Fluorescent ATP analog binding on-chip with ADK orthologs**

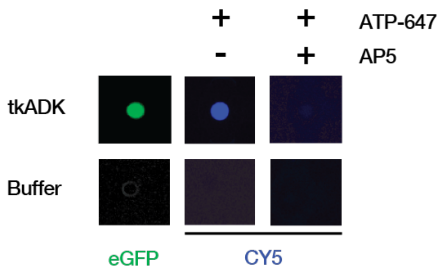


**Fig S4. ATP-647 is competed off by substrate analog on-chip.** A comparison of a buffer chamber and an exemplary ortholog *T. koda* ADK shows an AlexaFluor647 labeled ATP analog binding specifically to ADK and completed off when excess AP5 is introduced.

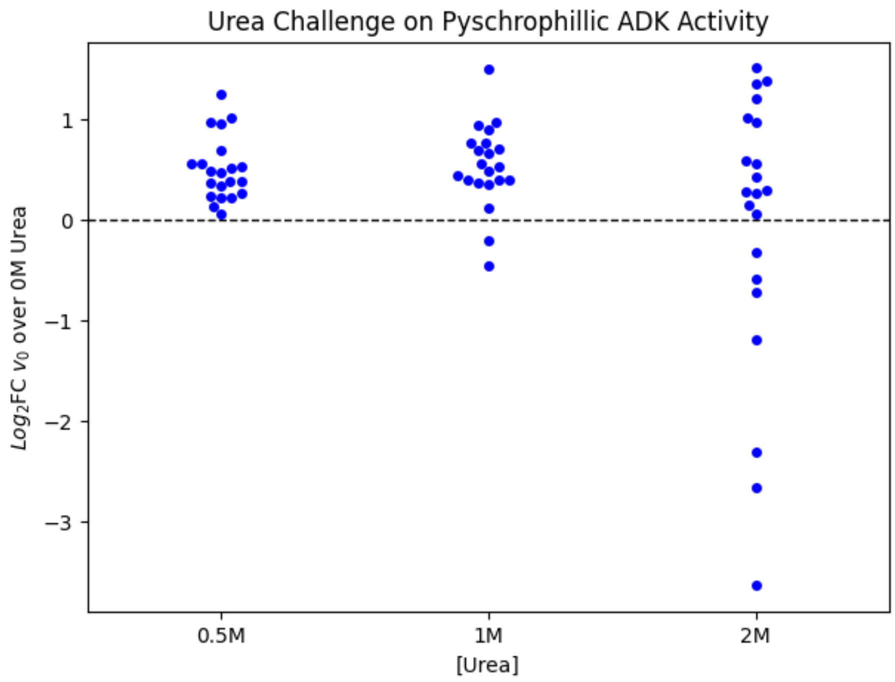


**Fig S5. Psychrophilic ADKs retain or gain activity at 0.5 M urea.** $Log_2$ fold-change in initial rate over 0M urea initial rate for ADKs with an associated growth temperature below 25C. A dashed line at 0 would represent no change in activity.

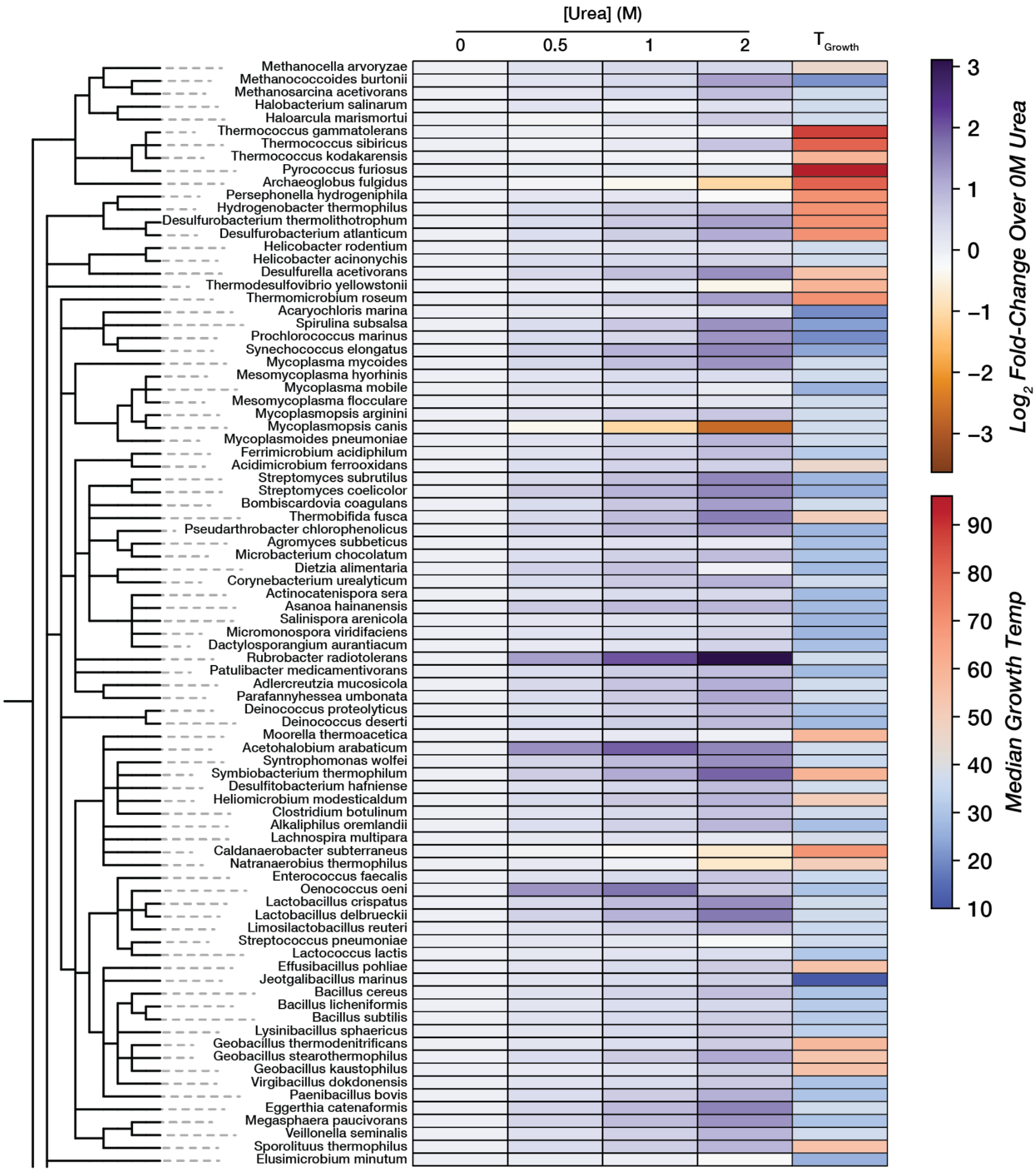


**Fig S6. Urea challenge on ADK ortholog activity.** Heat map of $\log_2$ fold-change over activity in 0M organized by taxonomy. The optimal growth temperature is also plotted.

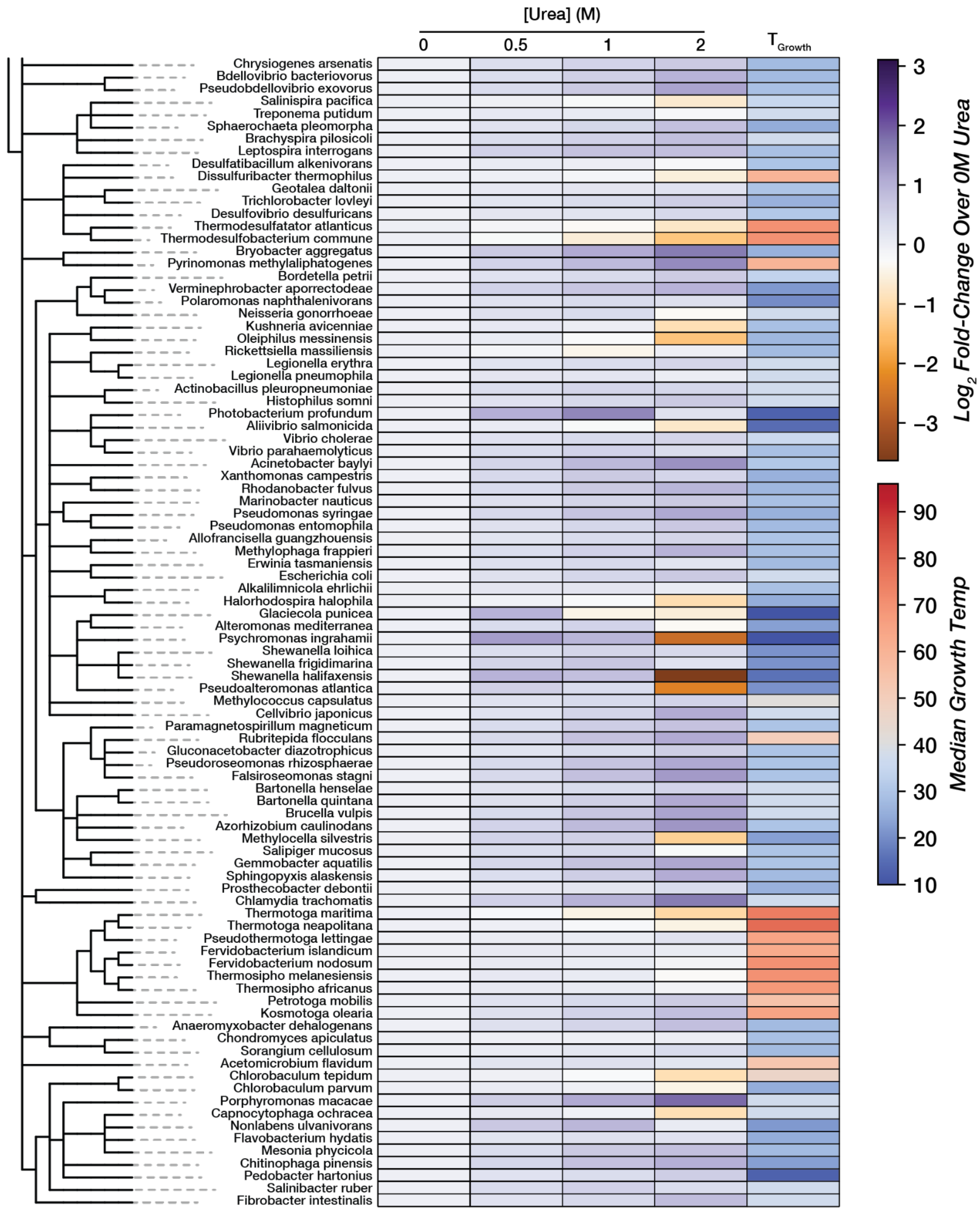


**Fig S6. (cont.) Urea challenge on ADK ortholog activity**

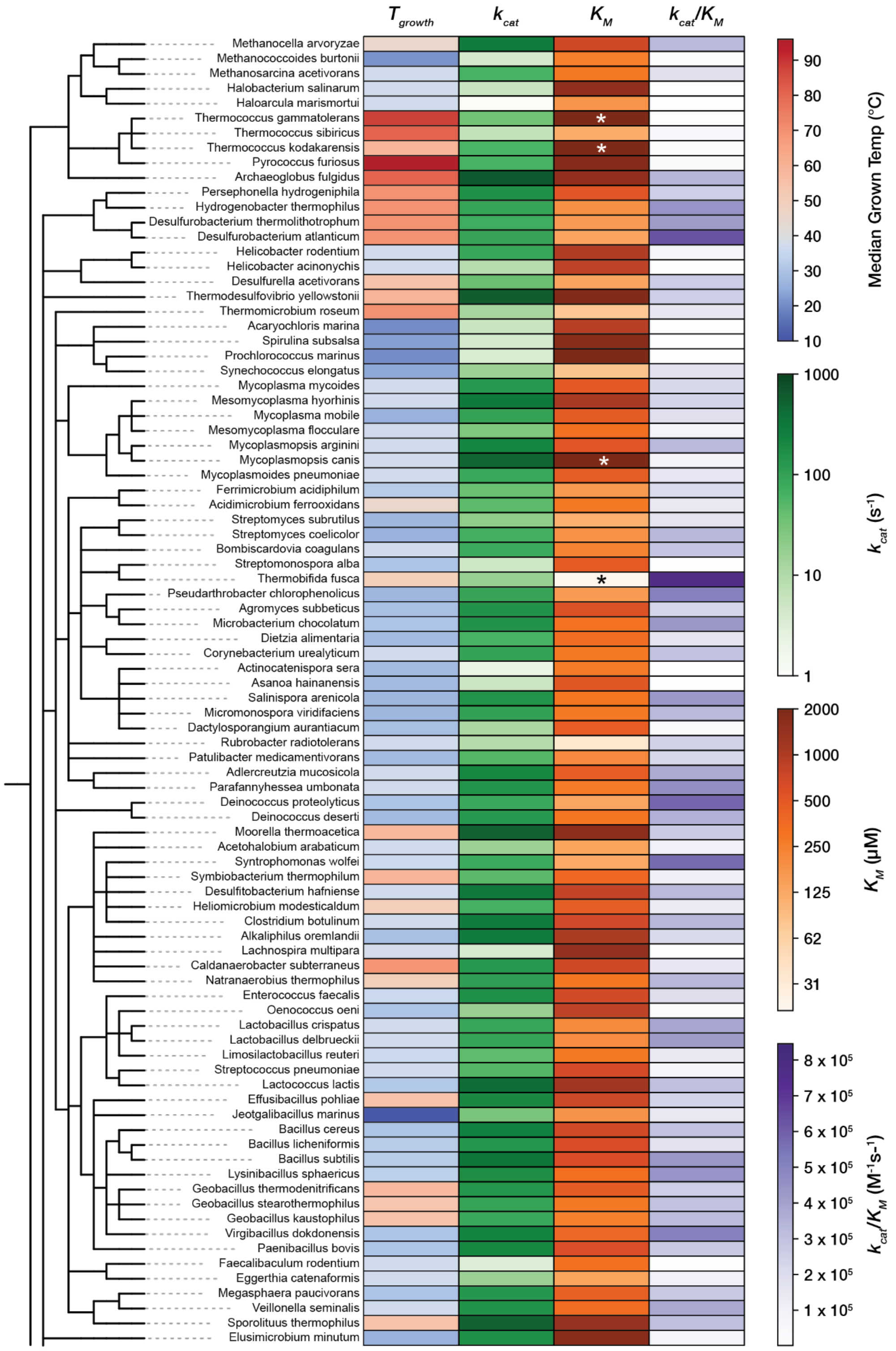


**Fig S7. Full activity and growth temperature heatmap.** Catalytic parameters and $T_{Growth}$ values for ADK orthologs are displayed as a heatmap across a taxonomic tree (Methods). $k_{cat}$ and $K_M$ are colored on log-scale. Orthologs with $K_M$ outside the range of the assay are labeled with asterisks (*).

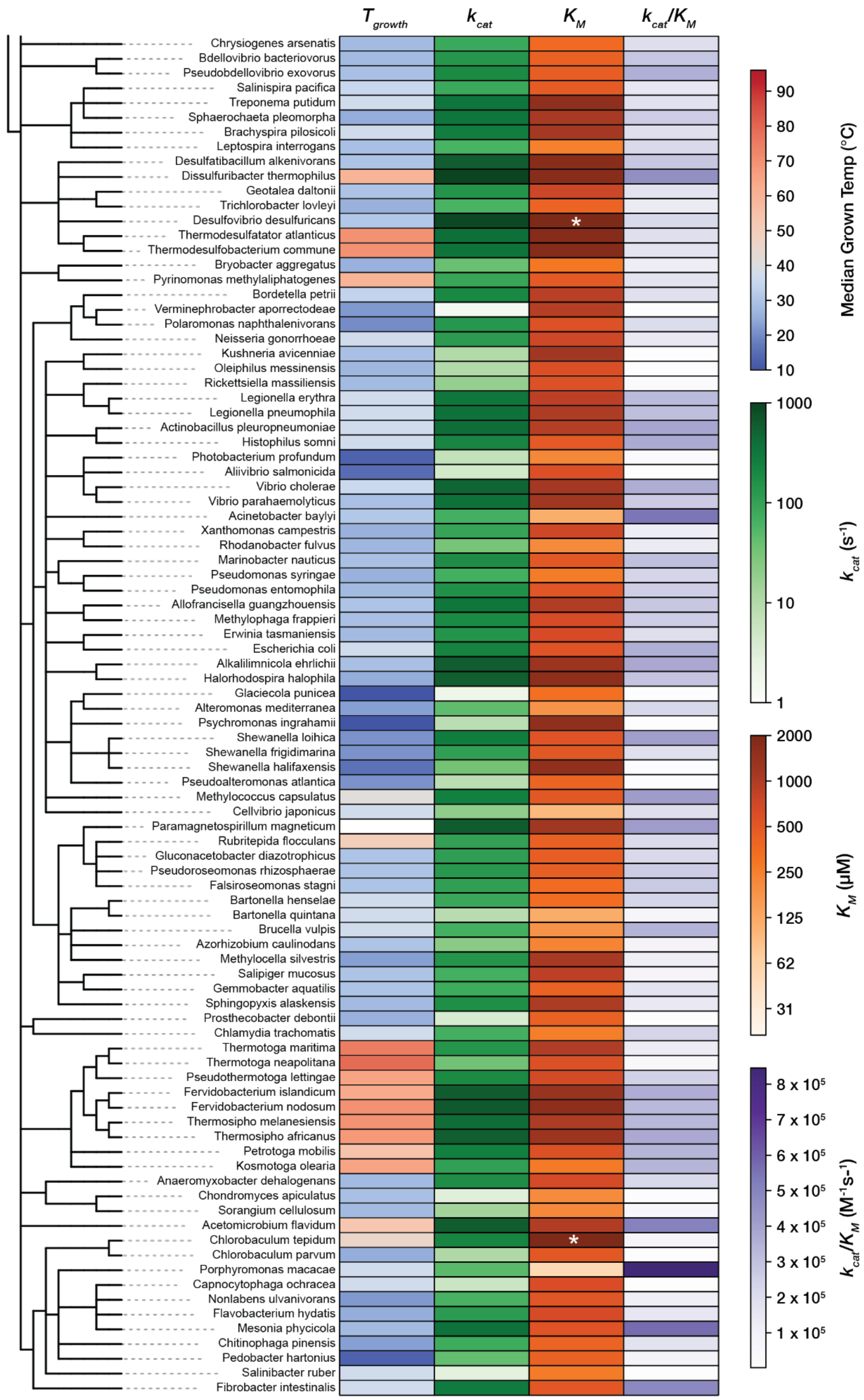


**Fig S7. cont. Full activity and growth temperature heatmap**

A

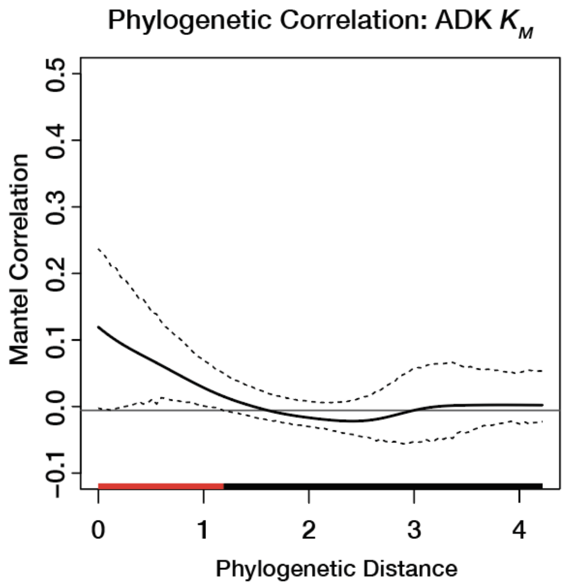


B

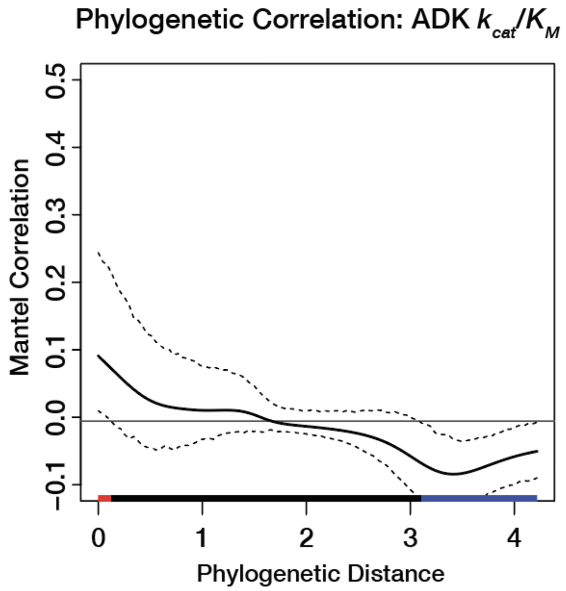


**Fig S8. Phylogenetic correlation for ADK $K_M$ and $k_{cat}/K_M$.** Phylogenetic signal computed using phylosignal R-package for (**A**) $K_M$ and (**B**) $k_{cat}/K_M$. Moran's I index of autocorrelation is plotted as a solid black line, with 95% confidence interval outlined by dashed black lines. The colored bar at the bottom encodes the significance of autocorrelation, where black=nosignficant, blue=signficant negative autocorrelation, and red=signficant positive autocorrelation.

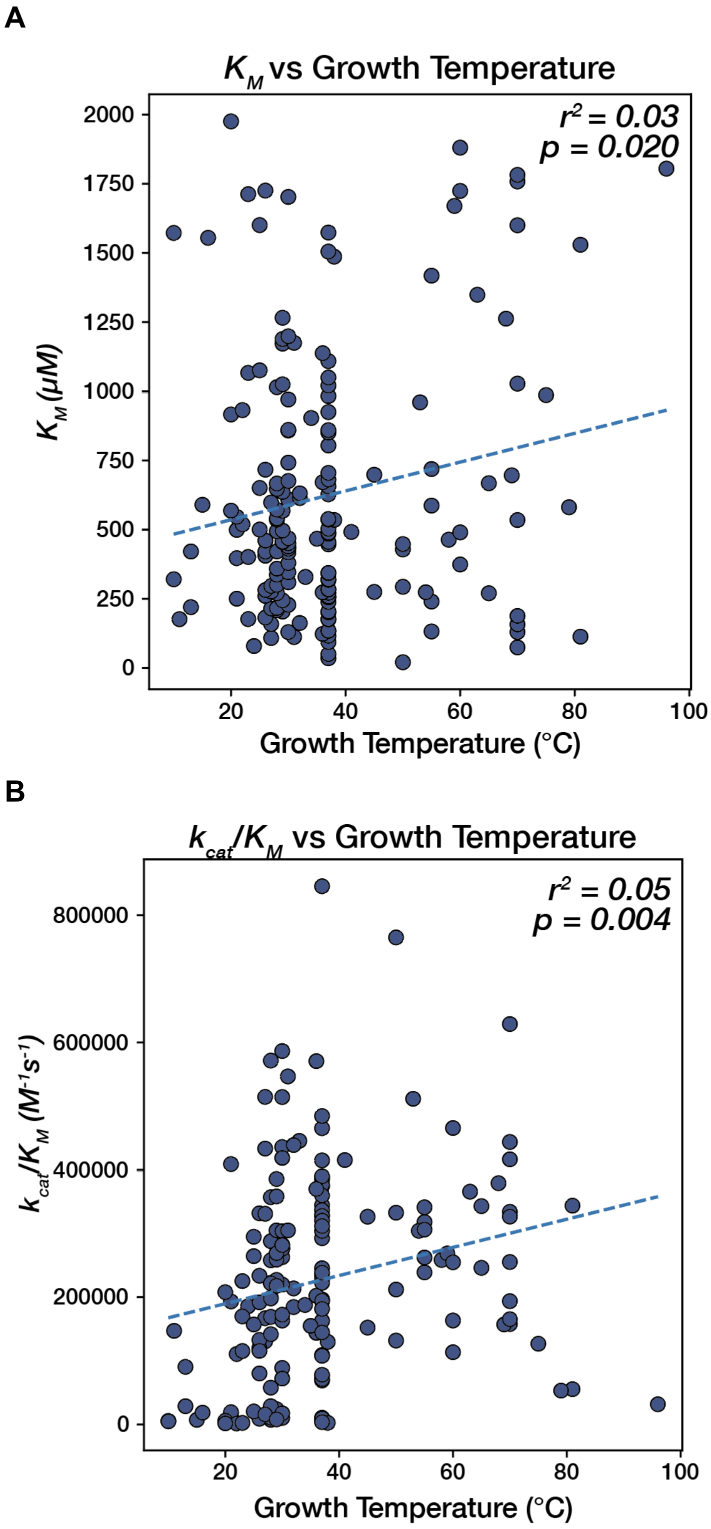


**Fig S9. Correlation of $K_M$ and $k_{cat}/K_M$ with growth temperature.** Linear regression of (**A**) Growth Temperature vs. $K_M$ and (**B**) Growth Temperature vs. $k_{cat}/K_M$. Regression lines are plotted as dashed blue lines, and $r^2$ and p-values are labelled on the plots.

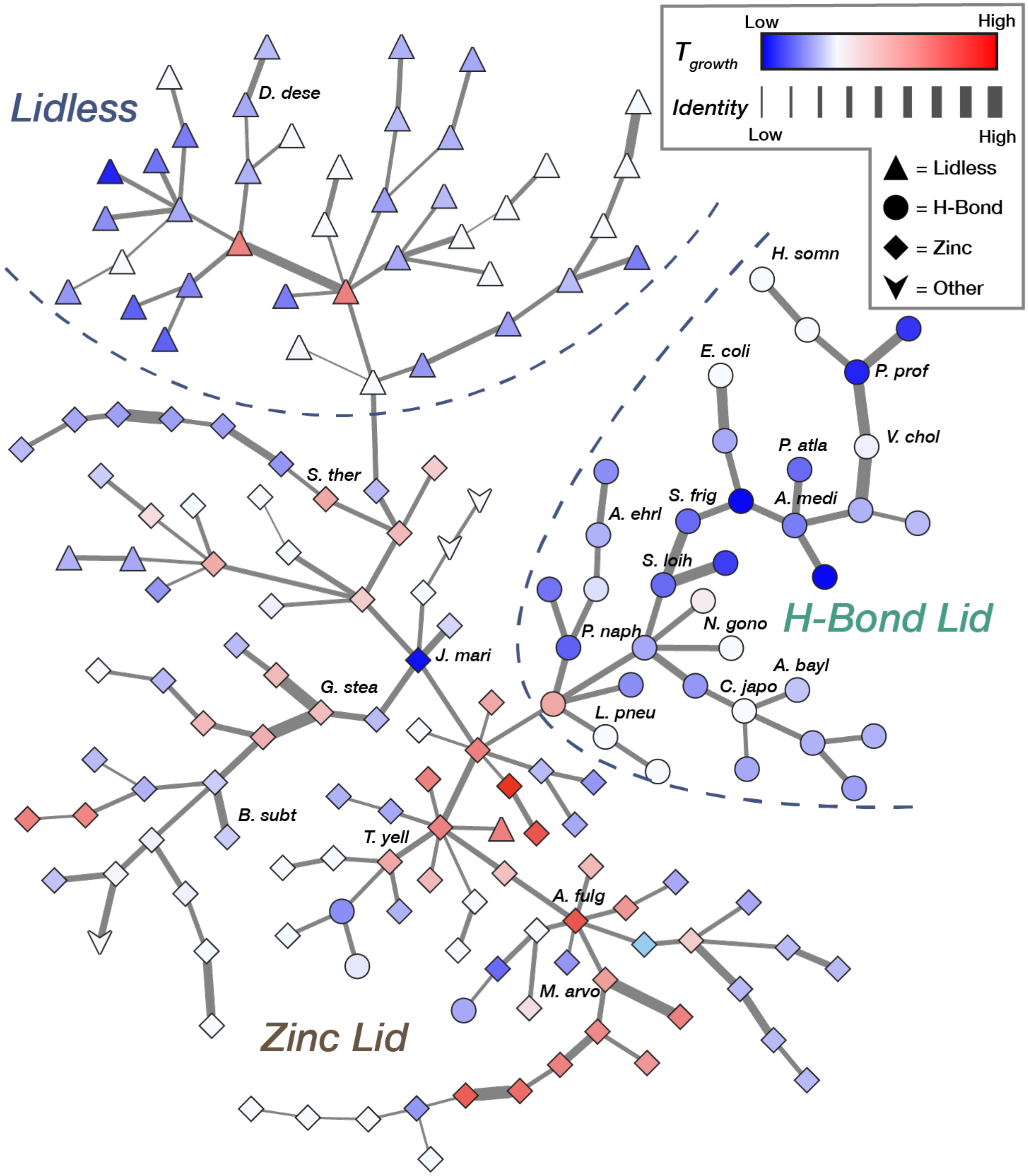


**Fig S10. ADK minimum spanning tree colored by temperature.** A minimum spanning tree constructed from a graph formed by connecting 175 ADK ortholog sequences (nodes) to each other by edit distance (edge weight) and traversing the graph to form a tree that minimizes edge weight and does not contain cycles. Node color corresponds to the median optimal growth temperature of the organism, and node shape encodes lid type. Edge thickness corresponds to the identity between two nodes. Rough lid-type neighborhoods are outlined with dashed lines.

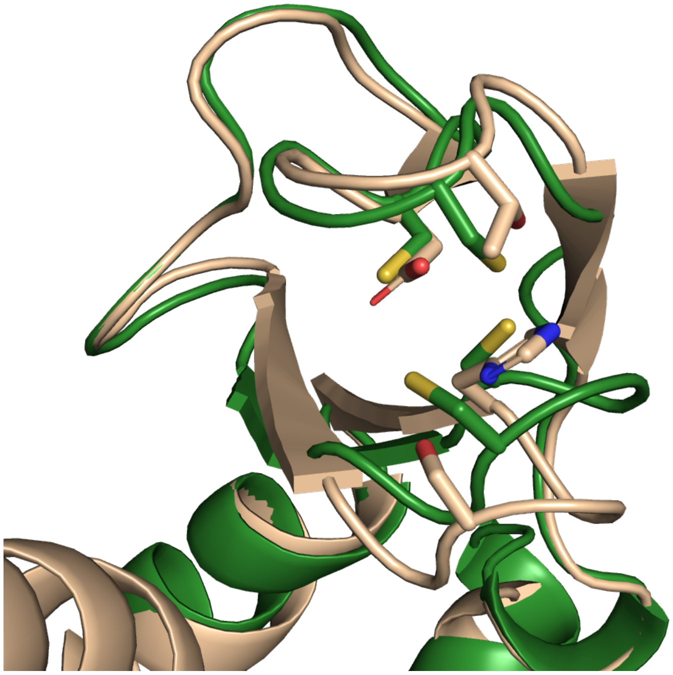

**Fig S11. Structural alignment of LID domain from $Zn^{2+}$/H-Bond ADKs.** Experimental structures of ecADK (PDB: 1AKE, tan) and gsADK (PDB: 1ZIP, green) are displayed, zoomed in on the LID domain. Key residues in the structural H-Bond motif of ecADK (His126, Ser129, Asp146, and Thr149, shown as sticks) align well with corresponding residues in the structural $Zn^{2+}$ binding motif of gsADK (Cys130, Cys133, Cys150, and Cys153, shown as sticks.

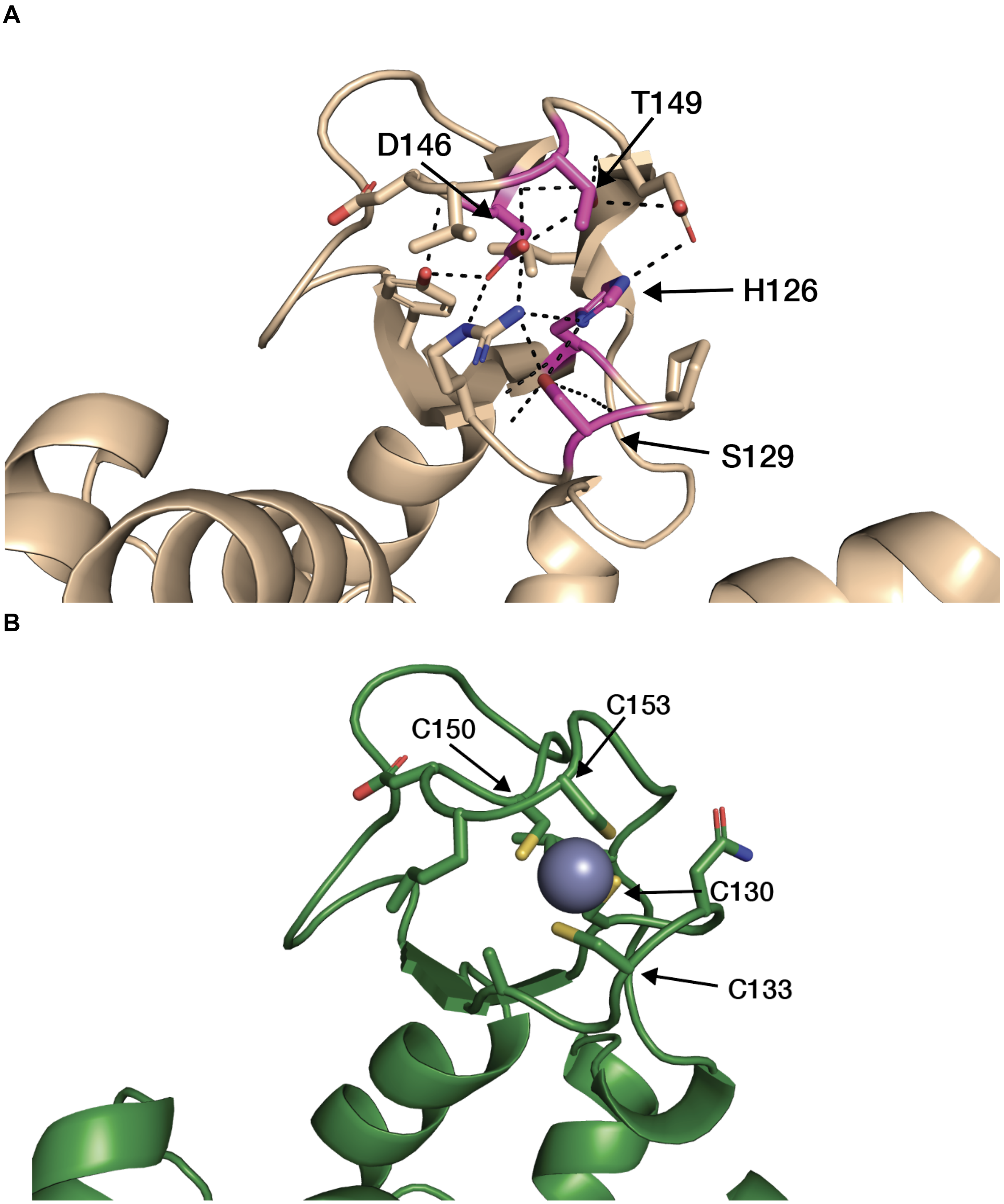


**Fig S12. Comparison of interactions in H-Bond and $Zn^{2+}$ lids. (A)** ecADK (PDB: 1AKE) with HSDT motif (pink) highlighted and hydrogen bonds (black dashed lines) displayed with neighboring residues. **(B)** gsADK with CCCC motif and neighboring residues shown as sticks, and $Zn^{2+}$ ion shown as a gray sphere.

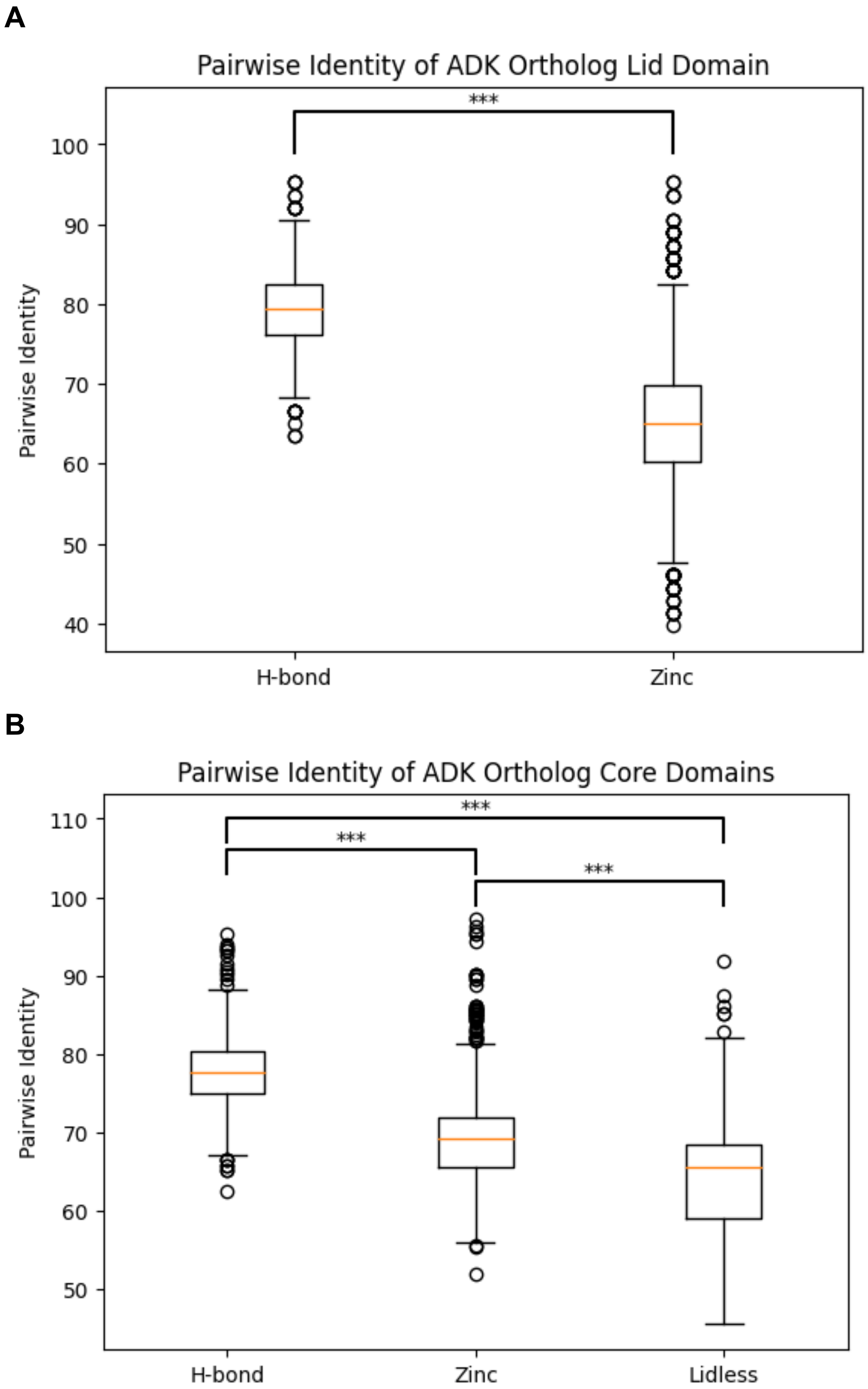


**Fig S13. Pairwise identity of ADK ortholog LID and CORE domains.** Domain regions are defined as by *Bae and Phillips* (55) (**A**) The LID domain H-Bond LID ADKs show higher pairwise sequence identity than the LID of $Zn^{2+}$ LID ADKs for ADK orthologs in the dataset (t-test, p-val < 1.0e-16). (**B**) Distribution of pairwise sequence identities within the CORE domain for each ADK lid type (ANOVA, F=1280.79, p-val<1.0e-16; Tukey-HSD, all comparisons p-adj < 1.0e-16).

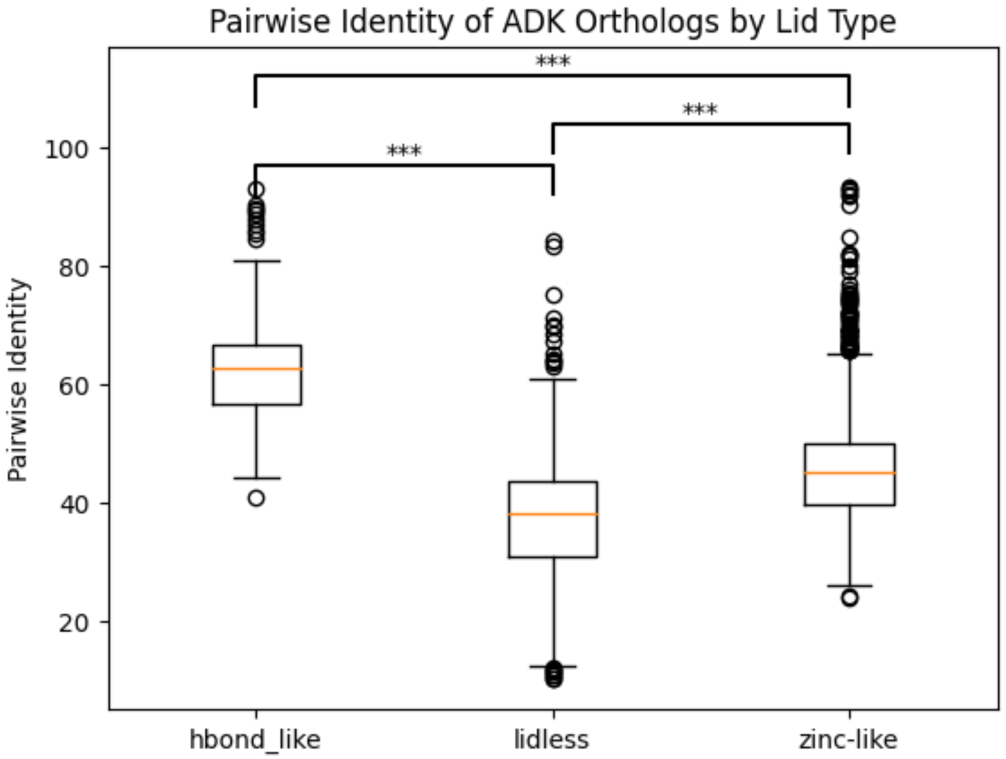


**Fig S14. Pairwise identity of ADK orthologs by lid type.** Box plot of pairwise sequence identity by lid-type for 175 ADK orthologs (ANOVA, F=1671.88, p-val<1.0e-16; Tukey-HSD, all comparisons p-adj < 1.0e-16).

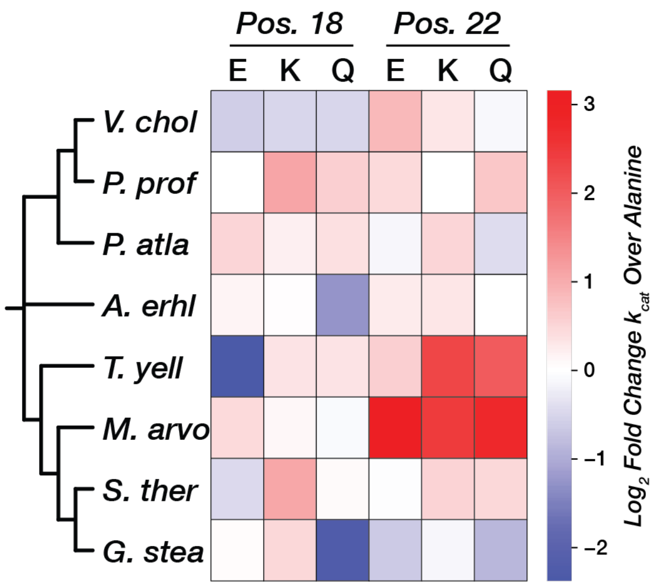


**Fig S15. Effect of charged or larger residues on $k_{cat}$ relative to alanine at positions 18 and 22.** $Log_2$ fold-change in $k_{cat}$ for glutamate, lysine, and glutamine mutants over alanine at positions 18 and 22 (ecADK numbering) are displayed as a divergent heatmap for 8 ADK orthologs, organized by a phylogenetic tree (branch length not encoded).

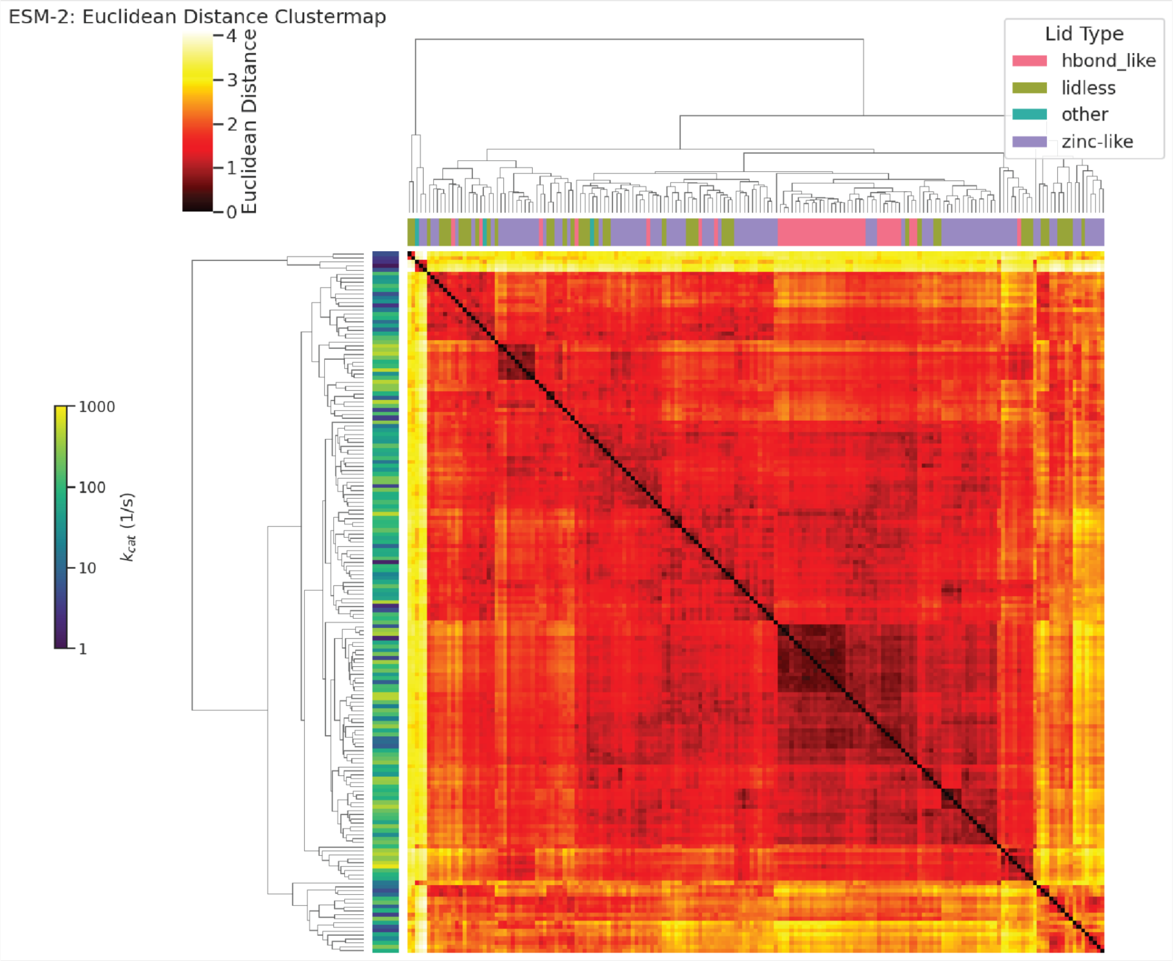


**Fig S16. Clustermap of ESM-2 embedded ADKs by euclidean distance.** Rows and columns are hierarchichally linked by euclidean distance. Columns are colored by lid type, and rows are colored by $k_{cat}$ on log scale.

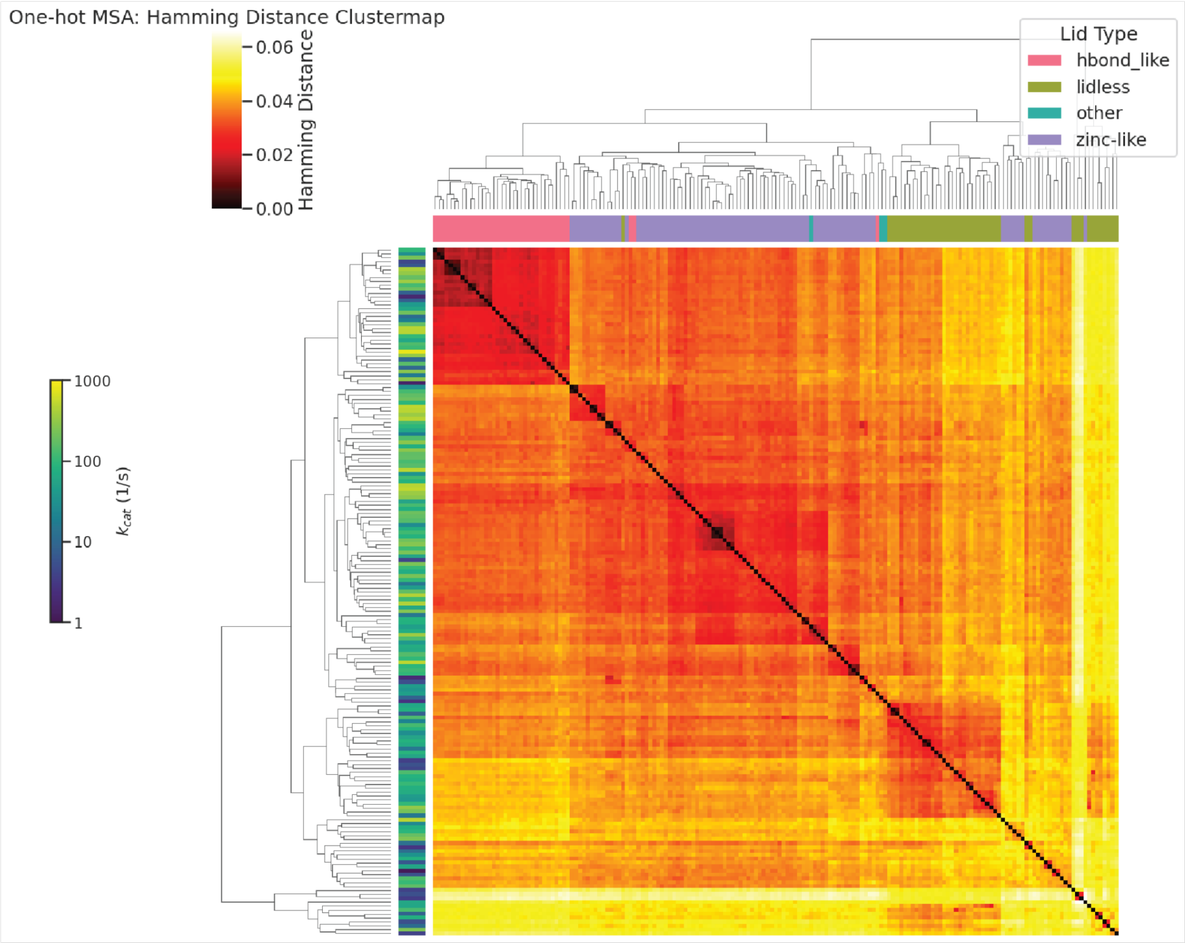


**Fig S17: Clustermap of One-Hot MSA encoded ADKs by Hamming distance.** Rows and columns are hierarchichally linked by euclidean distance. Columns are colored by lid type, and rows are colored by $k_{cat}$ on log scale.

**A**

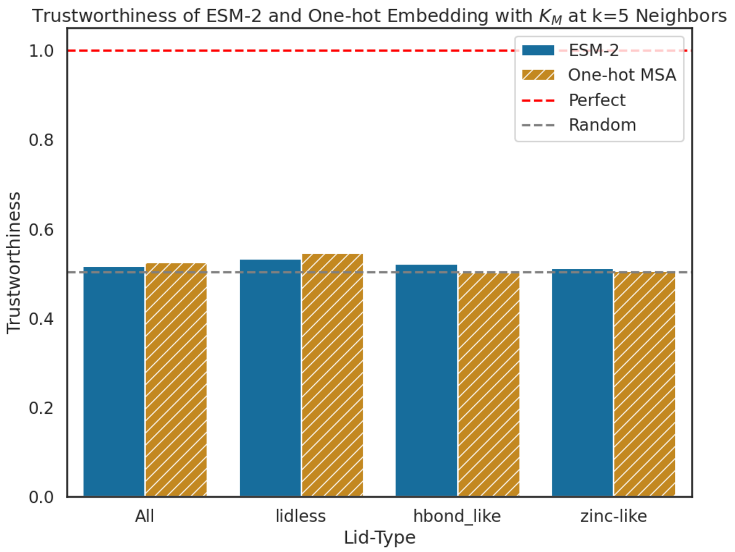


**B**

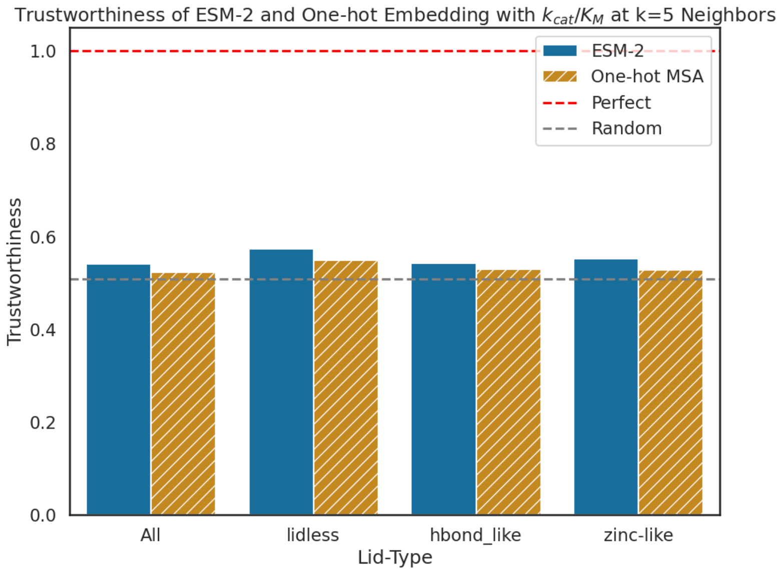


**Fig S18. Trustworthiness of ESM-2 embeddings vs. one-hot encodings for $K_M$ and $k_{cat}/K_M$.** Grouped barplot of trustworthiness of ESM-2 ADK embeddings (by Euclidean distance) and one-hot encoded MSA (by Hamming distance) with respect to $K_M$ computed at k=5 neighbors for (**A**) $K_M$ and (**B**) $k_{cat}/K_M$. Trustworthiness is computed for all sequences, as well as by lid-type. Perfect trustworthiness is plotted as a red dashed line, and random as a grey dashed line (average of 30 shuffles of corresponding embeddings).

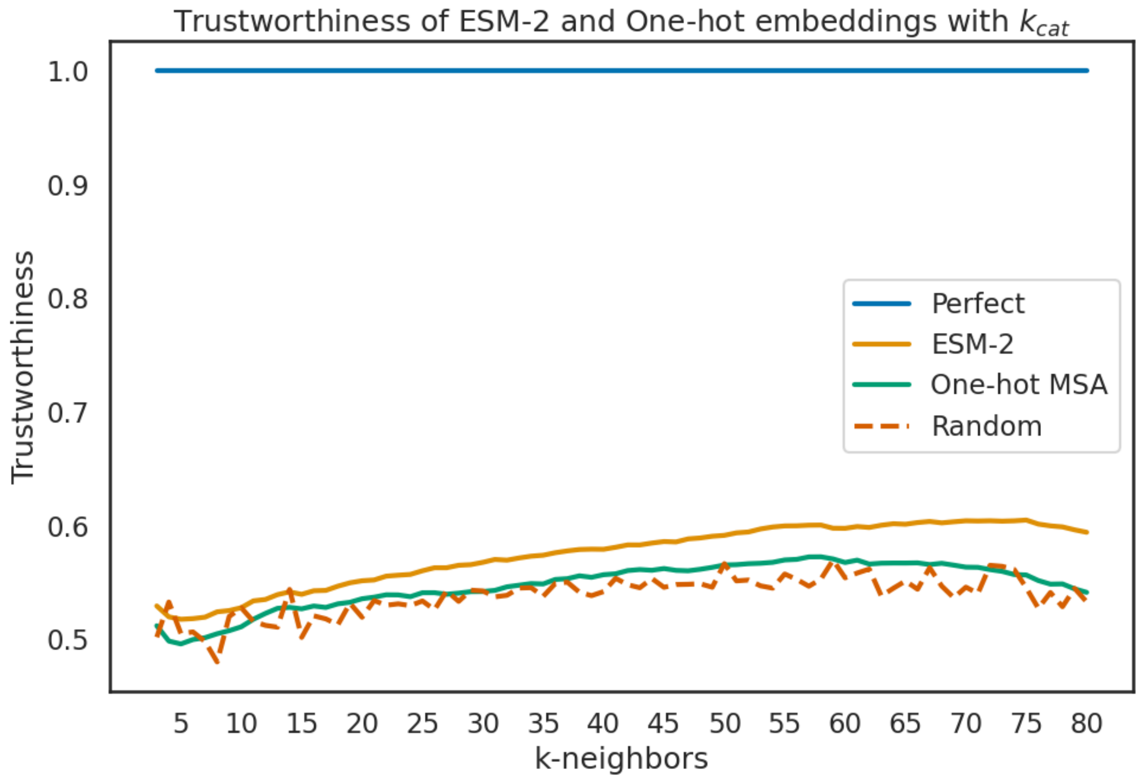


**Fig S19. $k_{cat}$ trustworthiness as a function of k-neighbors for ESM-2 embeddings and one-hot encoded MSA.** Trustworthiness is plotted as function of k-neighbors for ESM-2 embeddings and a one-hot encoded MSA with respect to $k_{cat}$ (based on euclidean and hamming distances, respectively). Perfect trustworthiness is plotted as a blue line ($k_{cat}$ vs. $k_{cat}$), and random as a red dashed line ($k_{cat}$ vs. average of 30 shuffles of corresponding embeddings).

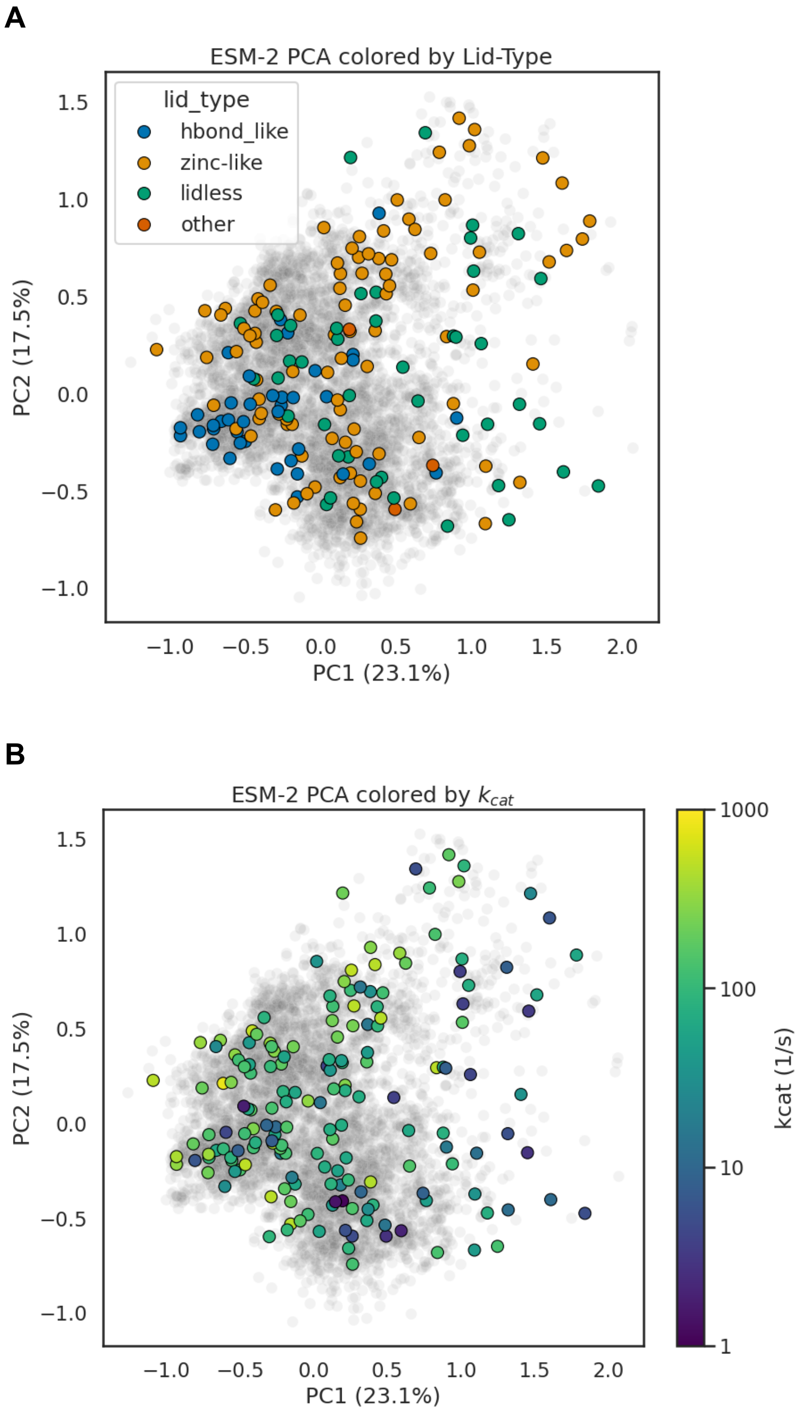


**Fig S20. Lidtype and $k_{cat}$ mapped onto ESM-2 embedding PCA.** (**A**) The first two principle components of a PCA of ESM-2 embeddings are plotted for all ~5,000 representative ADK sequences in grey, and sequences with measured catalytic parameters are colored by lid type. Explained variance for each component is annotated on the axis label. (**B**) The first two principle components of a PCA of ESM-2 embeddings are plotted for all ~5,000 representative ADK sequences in grey, and sequences with measured catalytic parameters are colored by $k_{cat}$.

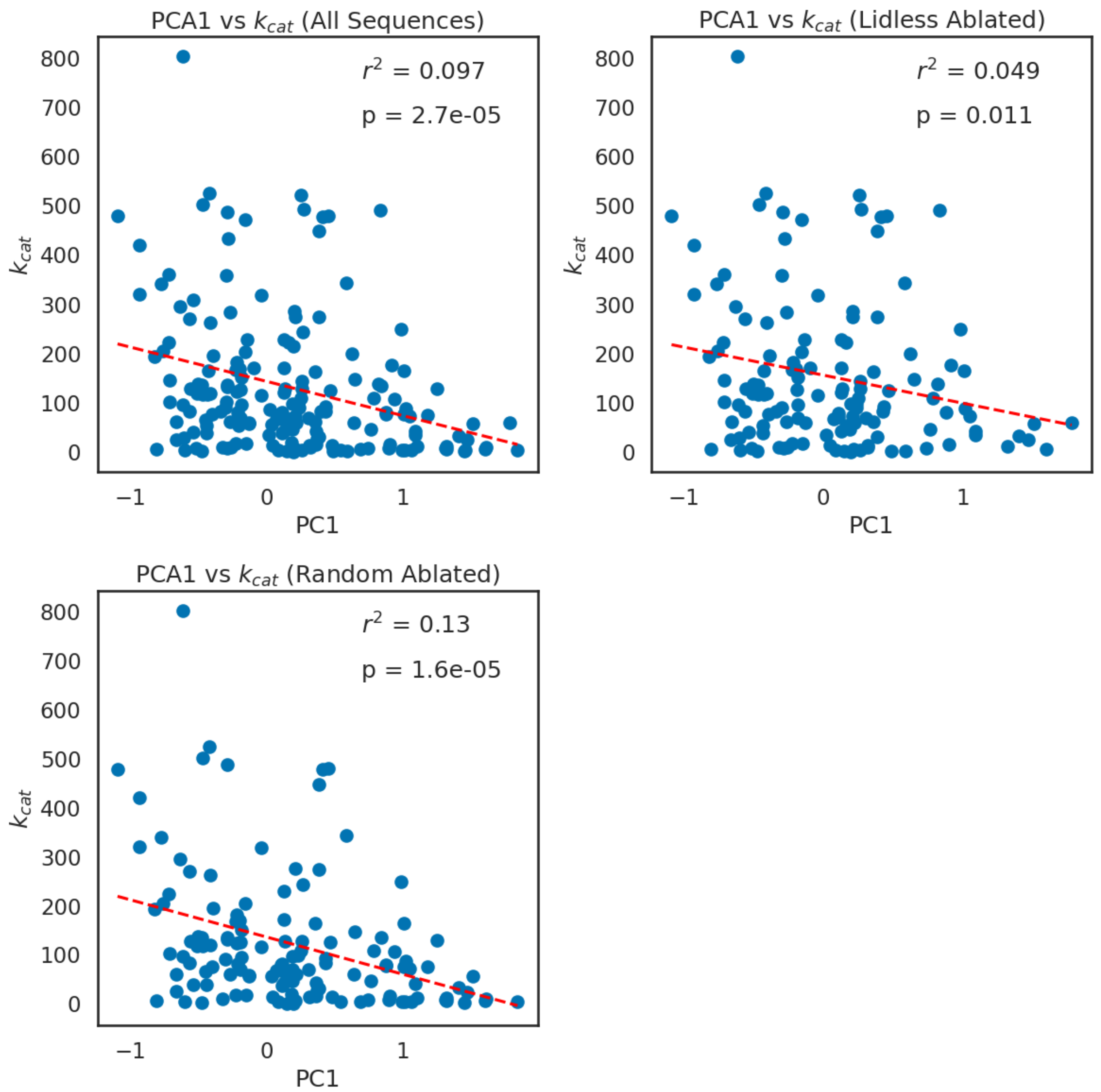


**Fig S21. Linear regression between first principle component of ADK ESM-2 embeddings and $k_{cat}$.** Linear regression between the first principle component of ADK ESM-2 embeddings and $k_{cat}$ for all 175 ADK sequences, all sequences except lidless, and all sequences except a random subset (of the same size as lidless). Regression lines are plotted as dashed red lines, and $r^2$ and p-values are labelled on the plots.

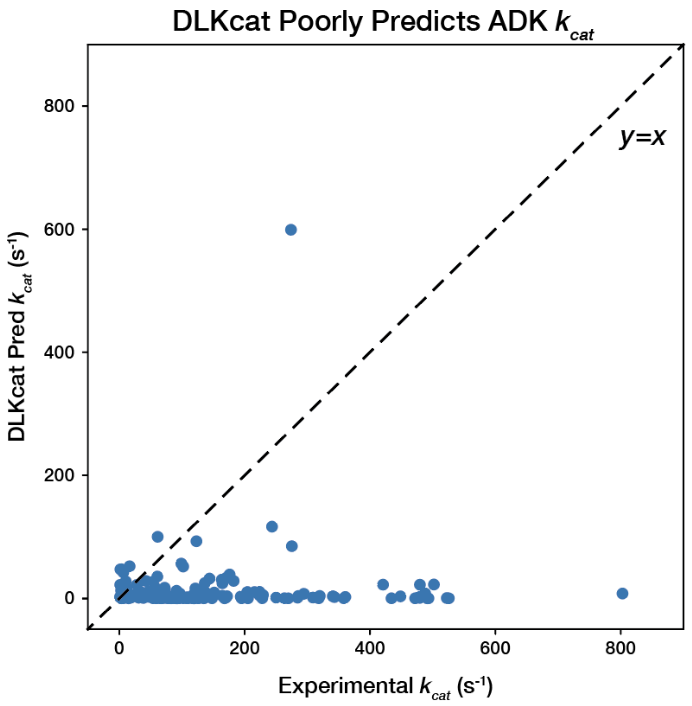


**Fig S22. Evaluation of DLKcat on 175 ADK sequences.** True values of $k_{cat}$ measured for 175 orthologs are plotted on the x-axis against predicted values from DLKcat on the y-axis (spearman rho=-0.09). A dashed black line represents perfect prediction (y=x).

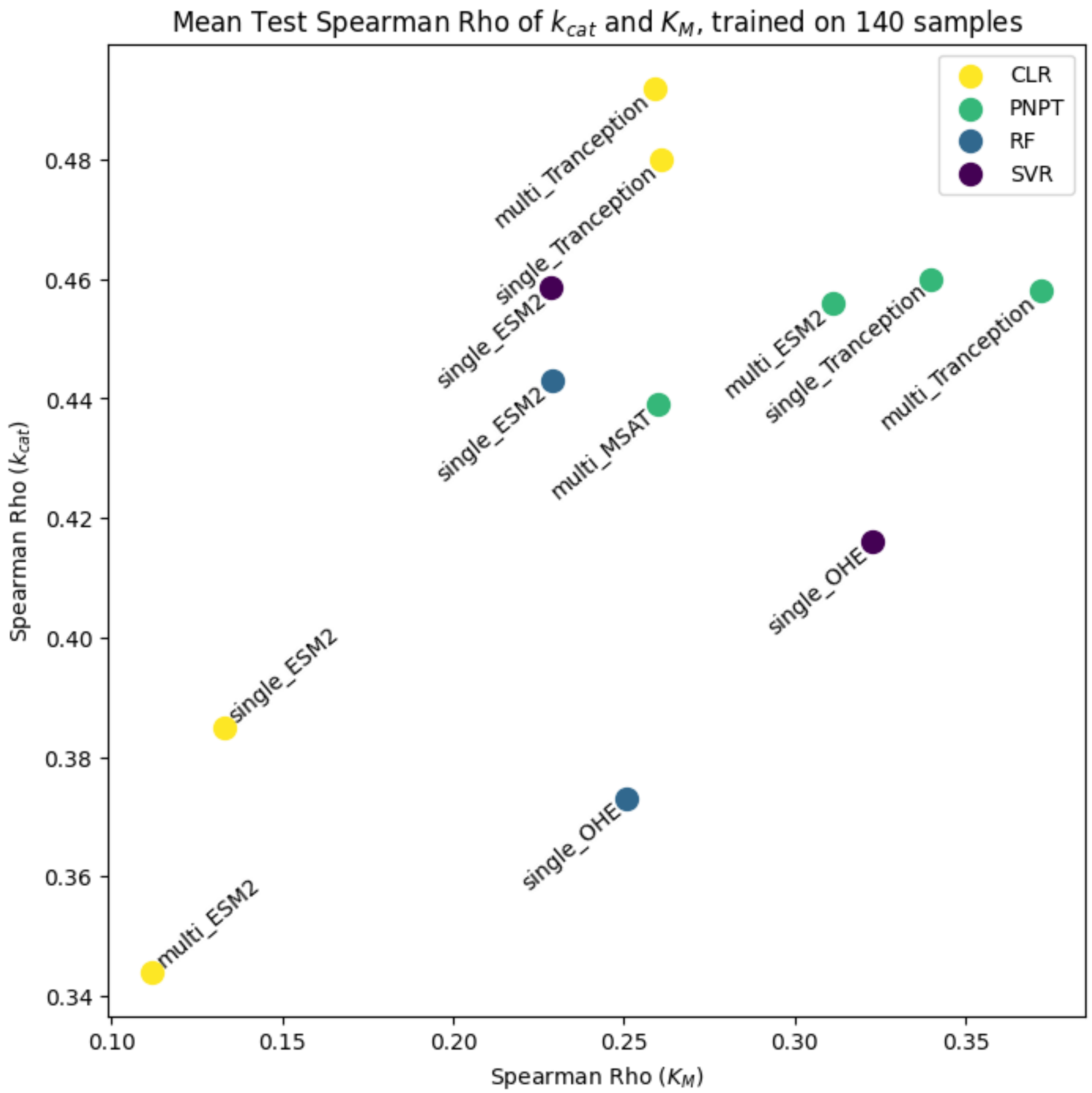


**Fig S23. Comparison of $k_{cat}$ and $K_M$ prediction performance for different model architectures, input embeddings, and hyperparameters.** Mean spearman rho across 30 samples for prediction of $k_{cat}$ and $K_M$ on a held out test set. Architecture is encoded by color (CLR=convolutional linear regression, PNPT=Protein NPT, RF=Random Forest, SVR=Support Vector regression). Text labels include whether or not a single target is predicted or multiple ($k_{cat}$, $K_M$, lid-type, and $T_{Growth}$), and the underling embedding (OHE=One-hot encoded MSA, MSAT=MSA Transformer).

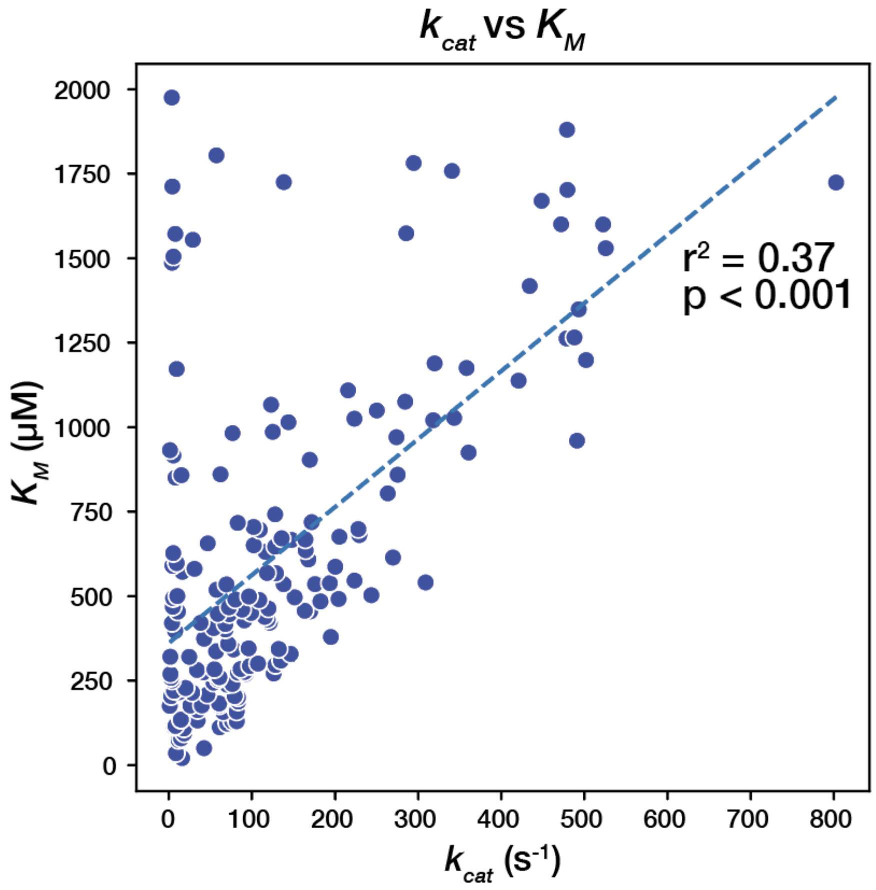


**Fig S24. Scatter plot of $k_{cat}$ vs. $K_M$ for ADK orthologs.** Linear regression line is plotted as a dashed blue line, with $r^2$ and p-values labelled on plot.

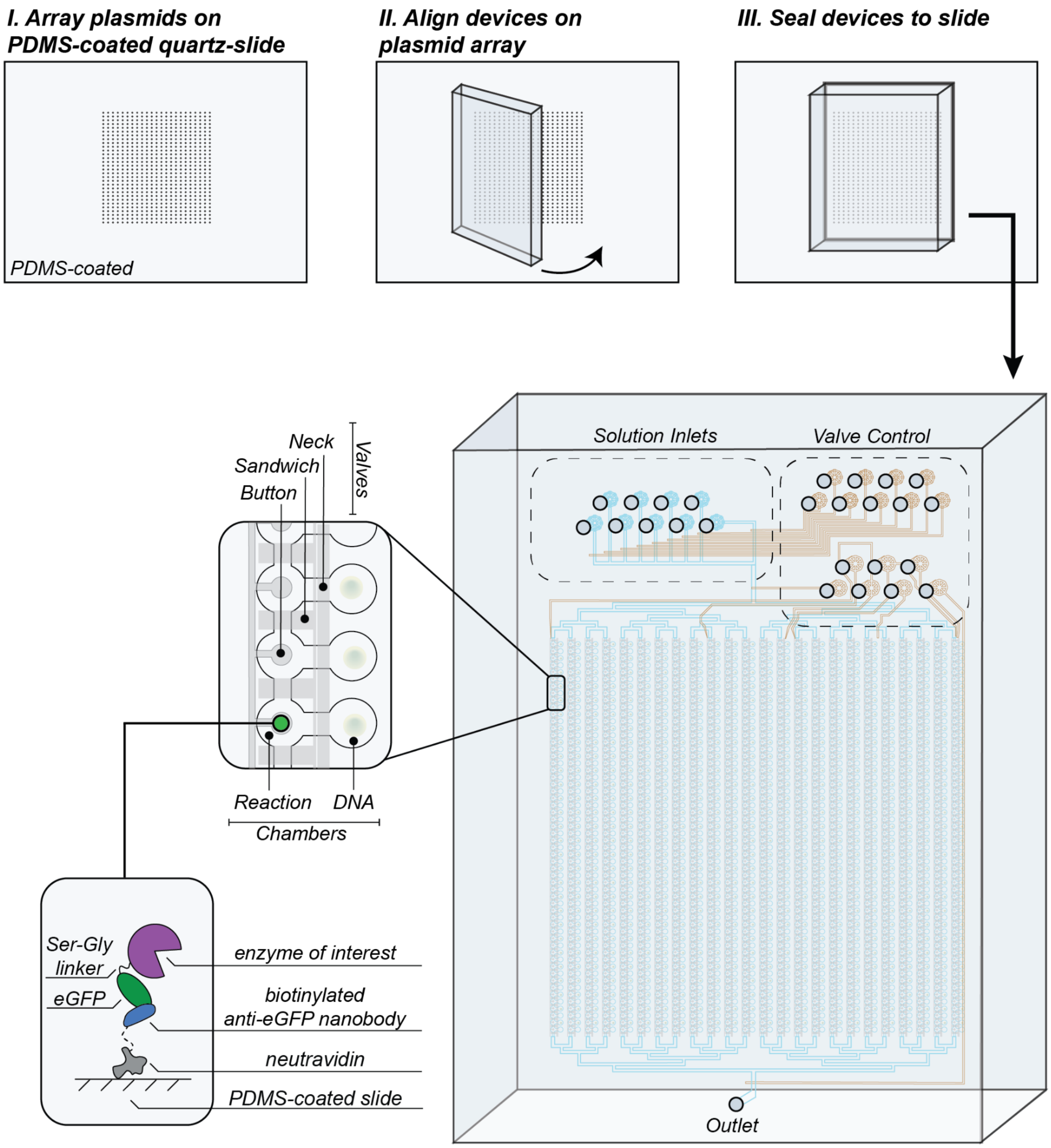


**Fig S25. Device architecture and alignment.** The process of plasmid arraying, and device assembly on a microscope slide is depicted. An enlarged device highlights the solution inlets used to introduce substrate and other reagents, and the valve control lines that manipulate the various valves used throughout the device (see inset for highlighted Neck, Sandwich, and Button valve). An example "pedestal" is then displayed in a reaction chamber, showing the capture strategy for enzymes of interest.

A

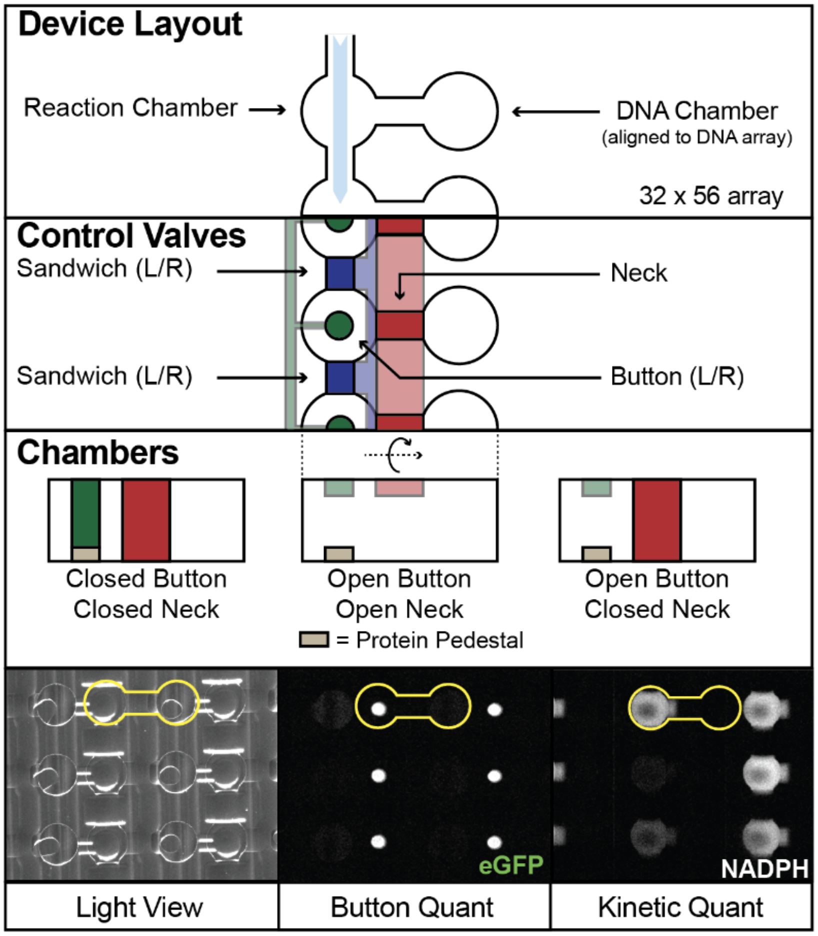


B

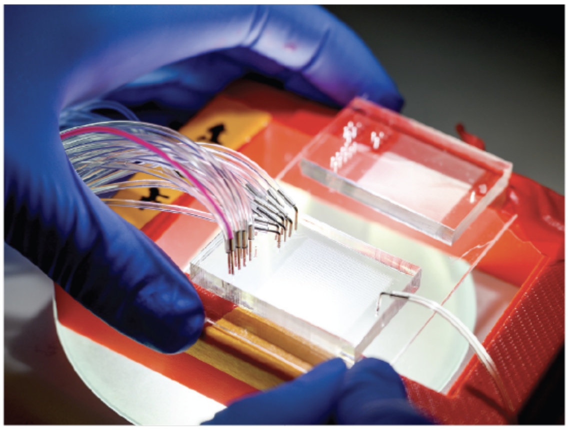

**Fig S26. Device valve configuration and example images.** (A) The layout of the reaction and DNA chambers is shown with the direction of flow shown with a light blue arrow. Sandwich, Button, and Neck valves allow for the protection of the protein pedestal and isolation of each reaction and DNA chamber from one another. Example microscope images are shown (from left to right) with room lights on visualizing device features, in the eGFP channel displaying ADK variant “Buttons”, and in the NADPH channel showing activity in reaction chambers. (B) Macroscopic image of an exemplary HT-MEK device.

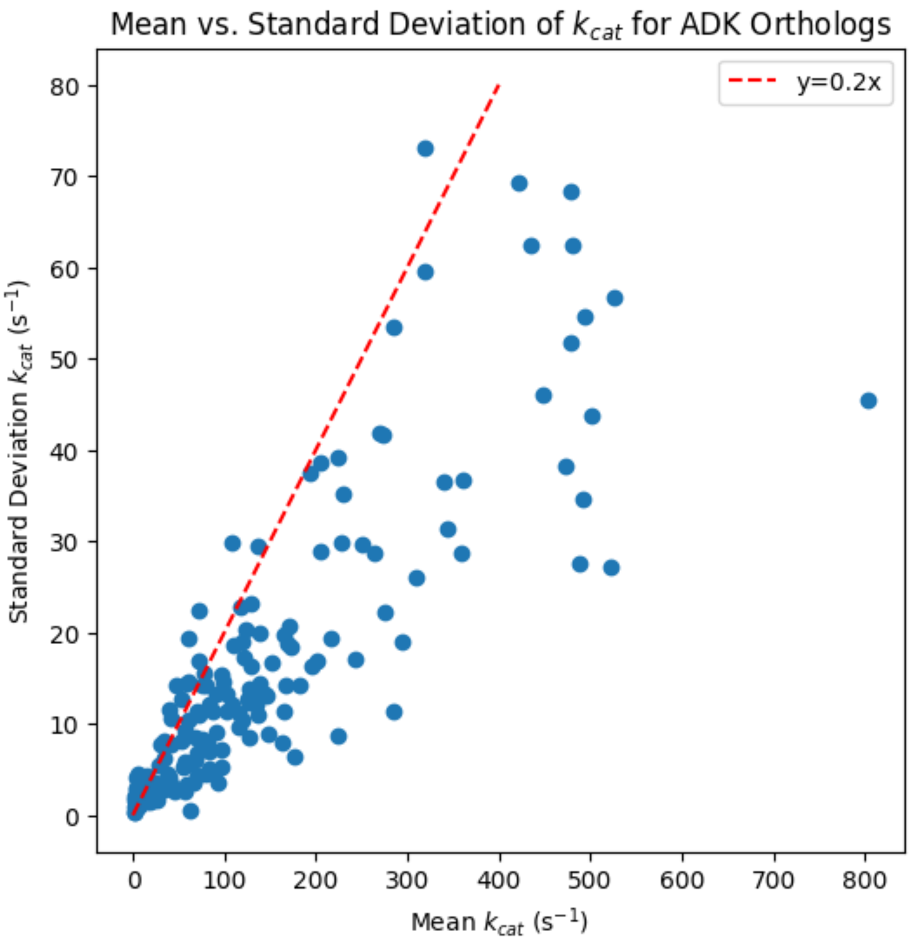


**Fig S27. Mean vs. standard deviation for ADK ortholog $k_{cat}$.** The mean versus standard deviation of $k_{cat}$ for 175 ADK orthologs is scatterplotted. A red dashed line shows that many ADK orthologs have standard deviations less than 20% of the magnitude of their associated mean value.

**A**

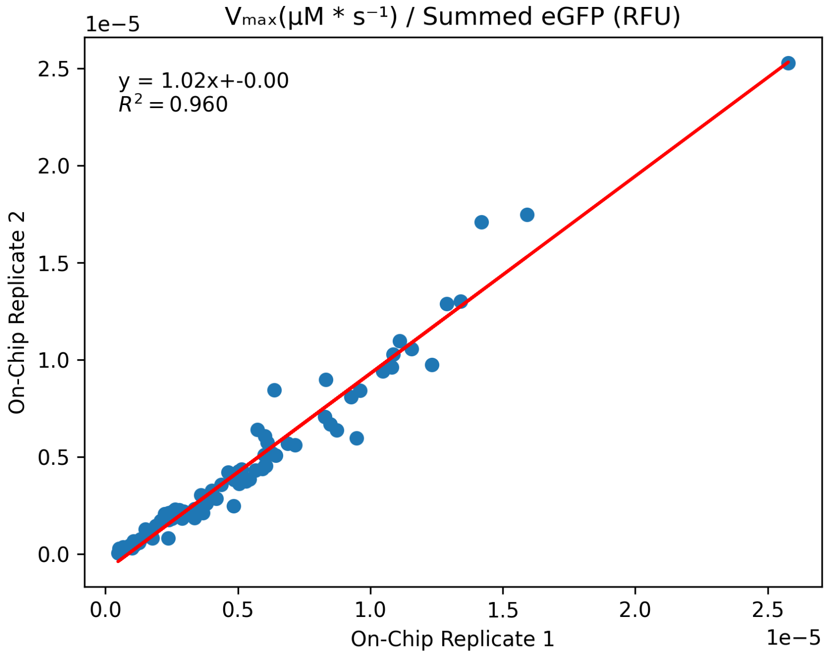


**B**

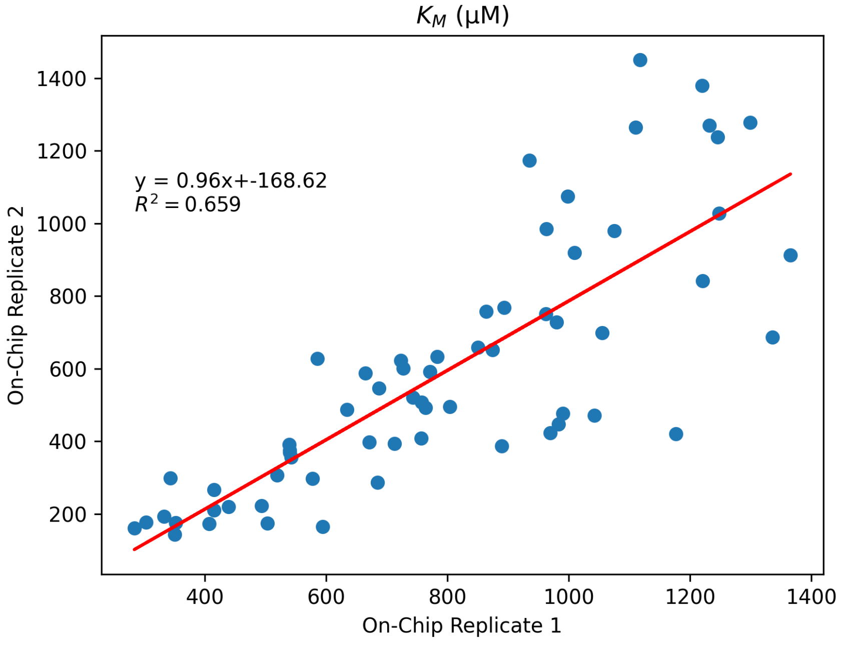


**Fig S28. Reproducibility of ADK catalytic parameters across multiple chips.** **(A)** Identical prints of an ADK ortholog library show highly consistent eGFP-normalized $V_{max}$ between two experiments run on different days ($r^2 = 0.96$). (**B**) $K_M$ for the same experiments is also comparable, though slightly noisier ($r^2 = 0.66$), likely due to challenges of detecting low concentrations of NADPH.
